# Large differences in collateral blood vessel abundance among individuals arise from multiple genetic variants

**DOI:** 10.1101/2023.05.28.542633

**Authors:** James E Faber, Hua Zhang, James G Xenakis, Timothy A Bell, Pablo Hock, Fernando Pardo-Manuel de Villena, Martin T Ferris, Wojciech Rzechorzek

**Author notes:** Corresponding author: James E. Faber, Department of Cell Biology and Physiology, 6309 MBRB, University of North Carolina, Chapel Hill, NC 27599-7545.

## Abstract

Collateral blood flow varies greatly among humans for reasons that remain unclear, resulting in significant differences in ischemic tissue damage. A similarly large variation has also been found in mice that is caused by genetic background-dependent differences in the extent of collateral formation, termed collaterogenesis—a unique angiogenic process that occurs during development and determines collateral number and diameter in the adult. Previous studies have identified several quantitative trait loci (QTL) linked to this variation. However, understanding has been hampered by the use of closely related inbred strains that do not model the wide genetic variation present in the “outbred” human population. The Collaborative Cross (CC) multiparent mouse genetic reference panel was developed to address this limitation. Herein we measured the number and average diameter of cerebral collaterals in 60 CC strains, their 8 founder strains, 8 F1 crosses of CC strains selected for abundant versus sparse collaterals, and 2 intercross populations created from the latter. Collateral number evidenced 47-fold variation among the 60 CC strains, with 14% having poor, 25% poor-to-intermediate, 47% intermediate-to-good, and 13% good collateral abundance, that was associated with large differences in post-stroke infarct volume. Genome-wide mapping demonstrated that collateral abundance is a highly polymorphic trait. Subsequent analysis identified: 6 novel QTL circumscribing 28 high-priority candidate genes harboring putative loss-of-function polymorphisms (SNPs) associated with low collateral number; 335 predicted-deleterious SNPs present in their human orthologs; and 32 genes associated with vascular development but lacking protein coding variants. This study provides a comprehensive set of candidate genes for future investigations aimed at identifying signaling proteins within the collaterogenesis pathway whose variants potentially underlie genetic-dependent collateral insufficiency in brain and other tissues.

## Introduction

Collaterals are vessels that reside within the watershed region between adjacent arterial trees where they cross-connect (anastomose) a fraction of the trees’ distal-most arterioles. They thus serve as alternative routes for perfusion when the trunk or a large branch of one of the trees experiences a slowly developing atherogenic or other structural narrowing, or a sudden occlusion caused by an embolic or athero-thrombotic clot. Most tissue types in healthy individuals have a network of native pre-existing collaterals. However, the amount of blood flow that they provide, which can be assessed following pathologic or diagnostic occlusion of a major artery supplying the tissue, exhibits wide variation among individuals^1–10^ For example, the level of flow mediated by the brain’s pial (leptomeningeal) collateral network, which can be estimated (ie, scored) in patients presenting with acute occlusion of the proximal middle cerebral artery (MCA)—the most common cause of ischemic stroke—partitions into ∼25% with low, ∼50% intermediate, and 25% high collateral flow, with the rate of progression and severity of subsequent ischemic injury following accordingly.^1–6^ Pre-treatment collateral score also correlates directly with the efficacy of thrombolysis, thrombectomy, and extension of the therapeutic time window for revascularization, and inversely with the risk for hemorrhagic transformation after treatment.^1–6^ The mechanisms responsible for this large variation, which extends to the collateral network in other tissues of the same individual,^11–19^ are not well understood and have become an area of increasing interest.

A similar wide variation in collateral flow has also been identified in different strains of mice and shown to result primarily from genetic background-dependent differences in the number and diameter of collaterals present in a given strain’s tissues.^12–19^ These differences arise from variation in the formation of collateral vessels, a unique endothelial cell-led process termed collaterogenesis, that occurs late during gestation after formation of the arterial and venous trees and capillary networks, ie, after developmental angiogenesis has occurred.^19–21^ This is followed by investment of the newly-formed collaterals with smooth muscle cells and subsequent growth in diameter, termed collateral maturation, that occurs during the first several weeks after birth— at which time the number and diameter of collaterals that will be present in the adult are established.^19–21^ While understanding of the pathways driving these processes is in its infancy, reverse-genetic studies of gene-targeted mice have identified involvement of several signaling proteins that are known to play primary roles in developmental angiogenesis, for example, VEGF-A, Flk1/Kdr, Notch1 and EphA4.^see^ ^references^ ^in^ ^Supplemental^ ^table^ ^4^ As well, unbiased forward-genetic studies employing F2 mice derived from intercrossing two common classical laboratory-bred strains, C57BL/6 (B6) and BALB/cBy, with abundant versus sparse collaterals respectively, identified four quantitative trait loci (QTL), denoted *Canq1-4*, located at genomic regions where none of the above-mentioned “angiogenesis” genes reside.^23, 24^ The causal gene for the largest-effect locus, *Canq1*, was recently identified^22^ and confirmed by others^25, 26^ as *Rabep2*, a novel gene involved in VEGF-A➔Flk1 downstream signaling within endothelial cells but not required for angiogenesis of non-collateral vessels.^22^ Identification of other naturally occurring variant genes that underly differences in the vigor of collaterogenesis has been hampered by the use of the above and other closely related classical strains that capture only a small fraction of the naturally occurring polymorphisms present in the mouse species (*Mus musculus*)^27–29^ and thus do not fully model the “outbred” human population.

The Collaborative Cross mouse genetic reference panel (CC) was recently developed to address this limitation.^30, 31^ The CC was created by reciprocal mating of five genetically diverse classical strains plus three strains derived from the three major wild sub-species of *Mus musculus*. This resulted in the segregation of greater than 50 million SNPs and structural variants that account for more than 90% of the general genetic variation in the mouse. This delineation of a much wider range of variation than achieved using F2 intercrosses of closely related strains greatly increases the ability to identify polymorphic genes that are responsible for differences in complex traits like collaterogenesis. Given that mice and humans share a high degree of genetic similarity and since the collaterogenesis pathway is presumably well-conserved among vertebrates, it is likely that *Rabep2* and other genes involved in collaterogenesis that harbor naturally occurring functional variants in mice are also determinants of collateral variation in humans.

Herein we performed genome-wide QTL mapping, bioinformatics-based SNP analysis, and functional assays within the CC to identify polymorphic loci and genes that underlie genetic-dependent variation in the abundance of collaterals. Such information is needed to better understand the fundamental mechanisms and key signaling proteins within the collaterogenesis pathway, the genetic basis for differences in collaterals among individuals, and to translate these findings going forward.

## Methods

### Mice

1434 mice were phenotyped for QTL mapping studies, 35 for infarct volume, 19 for abdominal wall and intestine angiography, ∼400 for in vivo assays, and ∼360 for generation of knockouts and other procedures. Founder (parental) strains of the CC were obtained from Jackson Laboratories (Bar Harbor, ME), and CC strains were obtained from the University of North Carolina Systems Genetics Core Facility (SGCF, Chapel Hill) between 2016 and 2018 for the initial screen, and again between 2020 and 2021 for the F1 validation experiments. C57BL/6J (B6), BALB/cByJ and *Rabep2*^-/-^ (B6.*Rabep2^em2Jef^*/J, JAX #029463)^22^ mice were from the first author’s colonies that were rejuvenated at 2-year intervals with breeders from Jackson Laboratories. Endothelial cell (EC)-specific inducible *C1galt1* knockdown mice were created by crossing B6.*Cdh5(PAC)-Cre-ER^T2^* mice (provided by Ralf Adams) to B6.129S1- *C1galt1^tm1.Rpmc^*/LxJ conditional/floxed mice^32^ (provided by Lijun Xia). C57BL/6N-*A^tm1Brd^ Cyb5r1^tm1a(KOMP)Wtsi^*/MbpMmucd (RRID:MMRRC_047271-UCD) cryo-recovered knockout mice, bred to a FLP-expressing strain to convert to the floxed allele, were from the Mutant Mouse Resource and Research Center (MMRRC) at the University of California at Davis (UCD), donated by The USC KOMP Repository, and originated from Kent Lloyd, UCD Mouse Biology Program. Ubiquitous and EC-specific inducible *Cyb5r1* knockdown mice were obtained by crossing this strain onto B6.*CAG-Cre* or B6.*Cdh5(PAC)-Cre-ER^T2^*, respectively. B6.*Igfn1^em1(IMPC)J^*/Mmjax cryo-recovered knockout mice (#042400-JAX) were from Jackson Laboratories. Cre activity was induced by tamoxifen (#T5648, Sigma) dissolved in 100% ethanol, diluted 1:9 with filtered corn oil, sonicated (final concentration 40mg/ml), and injected intraperitoneally at 100ul/30 gram body weight (4mg/30g) into pregnant dams beginning on embryonic day E15-16 for 3 consecutive days.

Both sexes were studied in ∼equal numbers, however n-sizes were not powered to test for sexual dimorphism. No sex-dependent differences have been observed for pial collateral number or diameter.^33^ Neonatal pups were postnatal day-0 (P0), young adults were 6 weeks-old, and adults were 10 weeks-old. CC mice, which exceeded 25 generations and were thus mostly inbred,^34^ were maintained under standard conditions at the SGCF or UNC vivariums; animal husbandry to obtain newborn pups for in vivo assays (see below) were performed by the Faber lab. All procedures were approved by UNC’s Institutional Animal Care and Use Committee and the NIH Guide for the Care and Use of Laboratory Animals (IACUC# 18-123.0-A, 04/2019).

### Generation of F2 populations

Following our initial study of 60 CC strains, we selected 12 with divergent collateral number but identical *Rabep2* haplotypes (this was done since we had already identified that *Rabep2* variants contribute to variation in collaterals^22^). We generated cohorts of F1s from each of these pairs and measured collaterals as described above. We selected two pairs of CC strains where our analysis of the strains themselves and their F1 phenotypes indicated we would be well-powered to identify genetic variants influencing their collateral differences in F2 intercrosses. Namely, we sought to identify strain-pairs with: (a) large between-strain differences in phenotype, (b) small within-strain variation in phenotype, and (c) F1 progeny whose phenotypes evidenced a largely additive nature. Subsequent analysis indicated that with ∼300 mice per intercross, we would be well-powered (genome-wide α = 0.05, β = 0.9) to identify loci contributing approximately 20% of the between-strain variation in collateral number. We contracted with the SGCF to set up two crosses in 2019 (CC049/TauUnc x CC053/Unc, CC055/TauUnc x CC036/Unc) based on these analyses. We targeted receiving ∼400 mice from each cross, with the expectation of roughly equal numbers of male and female mice per cross (confirmed). Inbred breeders for each pair were set up in reciprocal directions, and as much as possible. F2 mice were generated from all four possible F1-breeding combinations (eg, AxB dam crossed to AxB sire, etc) to minimize any potential mitochondrial or maternal effects, while maximizing the representation of X chromosome genotypes/diplotypes). F2 mice were received between 2019 and 2020.

### Angiography and morphometry of pial collaterals

As previously detailed,^35^ animals were deeply anesthetized with ketamine and xylazine and heparinized. The distal thoracic aorta was cannulated with pulled-out PE50, right atrium perforated, and phosphate-buffered saline (PBS) containing freshly prepared sodium nitroprusside (10^-4^M, for maximal dilation, maintained at 4°C in the dark) and Evan’s blue dye (for staining brain and adlumenal surface of ECs) were infused at ∼100 mmHg. Care was taken to prevent the introduction of bubbles. After exposing the neocortex through a craniotomy and positioning it under a stereomicroscope, Microfil (Flowtech Inc, Carver, MA) was infused until it began to appear in pial venules, to assure complete filling of precapillary pial vessels, followed by clamping the infusion line. After at least 20 minutes to allow the Microfil to set, brains were placed in 4% paraformaldehyde, and collaterals were imaged within 24h using a Leica fluorescent stereomicroscope. All collaterals between the anterior cerebral (ACA) and middle cerebral (MCA) artery trees of both hemispheres were counted by the same individual (HZ). Lumen diameter of each collateral was determined at midpoint from 50X images using ImageJ (NIH) and averaged for each mouse.

### Permanent middle cerebral artery occlusion and determination of infarct volume

As previously described,^35^ mice were anesthetized with ketamine and xylazine (100 and 10mg/kg, ip, respectively) and rectal temperature was maintained at 37±0.5°C. The right temporalis muscle was retracted along a 4 mm skin incision. The oblique edge of a 2.1 mm drill bit (#19007-21, FST, Foster City, CA) was used to thin an approximately 1 mm circle of bone overlying the distal M1-MCA. The thinned bone and dura were incised with a 27-gauge needle tip and reflected. The M1-MCA was cauterized (#18010-00, FST, Foster City, CA, custom modified tip) just distal to the lenticulostriate branches. The incision was closed with suture (∼15 min total surgery time), intramuscular cefazolin (50 mg/kg) and buprenorphine (0.1 mg/kg, repeated 12h later) were administered, and the animal was monitored in a warmed cage that maintained rectal temperature. Mice were euthanized 24h after MCA occlusion. Brains were sliced into ∼1 mm coronal sections and incubated in 1% 2,3,5-triphenyltetrazolium chloride in PBS at 37°C. Left and right forebrain and infarcted tissue in the right hemisphere were imaged on both sides of each slice with a stereomicroscope (ImageJ, NIH), average areas were determined for each slice, and tissue volumes were calculated. Percent infarct volume was normalized to forebrain volume, where infarct volume = the sum of the [(lesion area divided by total forebrain area) X 100] determined for 7 slices from each side of the brain, multiplied by slice thickness.

### SNP genotyping

Genomic DNA was extracted from tail snips using the DNeasy Tissue kit (Qiagen, Hilden, Germany). 500 ng of DNA was sent to Neogen, Inc (Lincoln, NE) for genotyping on the MiniMUGA array.^36^ Upon return, raw data were processed into relevant SNP calls (either homozygous for allele 1, heterozygous, homozygous for allele 2 or an N-call). Before further processing, we utilized mouse breeding records to ensure genotypes were consistent with expectations (ie, that the sex, mitochondrial origin and X-chromosome genotypes were consistent with the expected information based on the breeding records). Following this processing, we utilized the CC parental (inbred), F1s and F2 genotypes within each intercross to further filter to informative markers. Namely, we kept markers which (a) segregated between the two inbred parent pairs, (b) were heterozygous in the F1 samples, and (c) appeared as well behaving mendelian markers in the F2 group, with ∼1:2:1 ratios of genotypes at a marker, and ∼1:1 allele ratio in the population. These filtering steps took us from 10,819 total genomic markers to X for the CC049xCC053 cross and Y for the CC055xCC036 cross.

### Quantitative trait locus analysis

We conducted two types of QTL mapping: haplotype based mapping in the CC population, and interval mapping in our intercross populations. For the CC population, we utilized R/QTL2. We imported 8-state haplotype probability data at MegaMUGA density (∼77,000 positions across the autosomes and X chromosome) for each of the CC strains (http://csbio.unc.edu/CCstatus/index.py?run=FounderProbs)^34,37^ We calculated kinship for these strains, and mapped strain-average trait values against these data (trait ≈ haplotype + kinship + error versus a null model of trait ≈ kinship + error), and set significance thresholds based on 1,000 haplotype-phenotype permutations giving us empirical thresholds for p=0.01, 0.05, and 0.1). For the intercross populations, we imported our per-animal genotype and phenotype data into R/qtl. We initially utilized the b38 genome coordinates of the MiniMUGA markers^36^ to seed our genetic map, but then utilized the estimate.map() function to utilize the observed number of crossover events to recalculate the linkage map. We utilized standard Haley-Knott interval mapping in the scanone() function to assess the strength of the association (Logarithm of the Odds, or LOD score) between any interval and collateral numbers in each intercross. We did 1,000 permutations for each trait/intercross to generate an empirical distribution of expected spurious LOD scores, and used this distribution to calculate p=0.01, 0.05, and 0.1 thresholds. Once significant QTL were identified, we used the bayesint() function to identify the Bayesian credible intervals for the genomic positions of the causative variant(s) driving the locus. The peaks with the largest LOD values identified, *Canq6* and *Canq7* (see Results) were further investigated for possible interaction by fitting a model using the *Canq7* genotypes as a fixed effect, and then mapping the residuals as the new phenotypes for *Canq6*; the same was done after fixing *Canq6*. The distribution of the residuals was bell-shaped in both models. Rescans of the genotype-phenotype data showed that the peak locations of the two loci remained the same, the LOD score slightly increased and interval narrowed for *Canq7* when *Canq6* was fixed, and the LOD score slightly decreased and interval slightly increased for *Canq6* when *Canq7* was fixed. Peak locations for the other 5 significant loci (*Canq8*-*Canq12*) were also unaffected and LOD scores and intervals minimally affected after fixing either *Canq6* or *Canq7.* These findings indicate that the loci largely act independently and additively (see also **Supplemental figure I**).

### SNP analysis

We focused on a ±10 Mb region of the peak marker at each QTL to further interrogate protein coding genes as putative causal drivers (except for −5 to +10 Mb for *Canq9* due to haplotype heterogeneity). Based on the CC founder haplotypes present in a given intercross, we considered the following SNPs/Indel types: missense, in-frame deletion, in-frame insertion, initiator codon variant, stop gained/nonsense, stop lost, using both the Mouse Genomes Query (https://www.sanger.ac.uk/sanger/Mouse_SnpViewer/ rel-1505) and Mouse Phenome Database Sanger4 dataset (https://phenome.jax.org/projects/CGD-MDA1). SNPs that caused an amino acid change were analyzed for a potential detrimental effect on protein function using *in silico* prediction algorithms: Protein Variation Effect Analyzer (PROVEAN, http://provean.jcvi.org),^38^ Sorting Intolerant From Tolerant (SIFT) as implemented within PROVEAN,^39^ and SIFT as implemented within the Ensemble browser (http://sift.bii.a-star.edu//sg/). Protein-coding genes within a QTL interval were also queried for the presence of private SNPs/Indels in the CC.^34, 40^ No private missense mutations predicted to be deleterious or stop gained/lost mutations were found, except *Magea1* which has a private missense snp (C/T) in CCXXX at location 155089004 (p < 0.05) (see Results).

### In vivo collaterogenesis assay and AAV probes

Neonatal (P0) *Rabep2^-/-^* and various CC strain pups were anesthetized by placing on wet ice covered with a kimwipe for 2-3 min, then transferred to a 60×90×12 mm stainless steel block pre-chilled to 4°C and positioned under a stereomicroscope. Recombinant AAV9 harboring an expression plasmid for a given gene to be tested was injected (10 ul of 1×10^13^ vg/ml rAAV9) retro-orbitally using a 31-gauge needle attached to a 0.3 ml insulin syringe (BD Ultra-fine^TM^ II, Becton Dickinson, Franklin Lakes, NJ).^41, 42^ Successful retro-orbital injection was confirmed by flushing of blood from the superficial facial vein. E-coli plasmids were constructed with the following features: CMV (ubiquitous) promotor, ORF of the given gene, P2A linker, enhanced green fluorescence protein (*GFP*) and WPRE post-transcriptional regulatory element (VectorBuilder, Chicago, IL). The *GFP* element was not included for genes whose ORFs exceeded 798 amino acids to assure robust packaging and expression by AAV9 (*Pld1, Usp13, Cfhr4, Sh3rf1*). Vectors for genes whose ORFs exceeded 1036 were not studied because of these limitations, with the exception of *Jak3* (ORF 1100) wherein twice the standard dose was used (see Results). The sequence of the ORF for candidate genes was for the C57BL/6J reference strain to provide the full-length protein to compete with the predicted LoF protein for binding to its partner(s) and/or activation of downstream effectors. The base-sequence at the indicated SNPs for the genes listed in Table 1 for the high-collateral parental strains of the two F2 populations were the same as present in the C57BL/6J reference strain (unless indicated otherwise), justifying using the latter’s ORF in the AAV9 constructs.

**Table 1.**
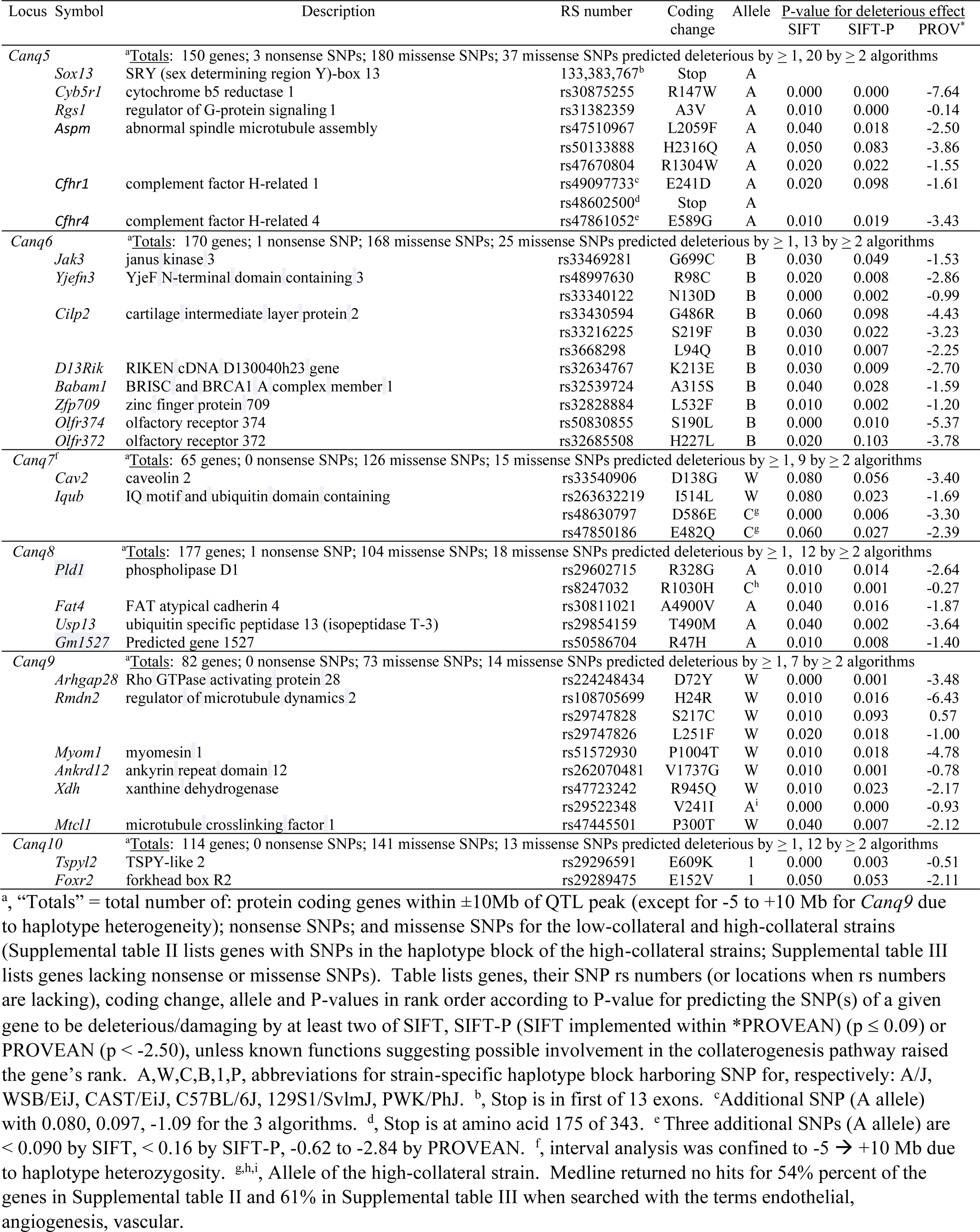
Candidate genes underlying *Canq5*-to-*Canq10* for collateral insufficiency/low collateral number. Genes with SNP(s) in the haplotype block of the low-collateral strains in the crosses shown in Figure 4A/Supplemental table 1 are ranked highest-to-lowest based on P-values for presence of non-sense SNPs, missense SNPs predicted to be deleterious, and known functions suggesting possible involvement in the collaterogenesis pathway.

| Locus | Symbol | Description | RS number | Coding change | Allele | P-value for deleterious effect |  |  |
| --- | --- | --- | --- | --- | --- | --- | --- | --- |
|  |  |  |  |  |  | SIFT | SIFT-P | PROV<sup>a</sup> |
| *Canq5* | | <sup>a</sup>Totals: 150 genes; 3 nonsense SNPs; 180 missense SNPs; 37 missense SNPs predicted deleterious by $\geq 1$, 20 by $\geq 2$ algorithms | | | | | | |
|  | *Sox13* | SRY (sex determining region Y)-box 13 | 133,383,767<sup>b</sup> | Stop | A |  |  |  |
|  | *Cyb5r1* | cytochrome b5 reductase 1 | rs30875255 | R147W | A | 0.000 | 0.000 | -7.64 |
|  | *Rgs1* | regulator of G-protein signaling 1 | rs31382359 | A3V | A | 0.010 | 0.000 | -0.14 |
|  | *Aspm* | abnormal spindle microtubule assembly | rs47510967 | L2059F | A | 0.040 | 0.018 | -2.50 |
|  |  |  | rs50133888 | H2316Q | A | 0.050 | 0.083 | -3.86 |
|  |  |  | rs47670804 | R1304W | A | 0.020 | 0.022 | -1.55 |
|  | *Cfhr1* | complement factor H-related 1 | rs49097733<sup>c</sup> | E241D | A | 0.020 | 0.098 | -1.61 |
|  |  |  | rs48602500<sup>d</sup> | Stop | A |  |  |  |
|  | *Cfhr4* | complement factor H-related 4 | rs47861052<sup>e</sup> | E589G | A | 0.010 | 0.019 | -3.43 |
| *Canq6* | | <sup>a</sup>Totals: 170 genes; 1 nonsense SNP; 168 missense SNPs; 25 missense SNPs predicted deleterious by $\geq 1$, 13 by $\geq 2$ algorithms | | | | | | |
|  | *Jak3* | janus kinase 3 | rs33469281 | G699C | B | 0.030 | 0.049 | -1.53 |
|  | *Yjefn3* | YjeF N-terminal domain containing 3 | rs48997630 | R98C | B | 0.020 | 0.008 | -2.86 |
|  |  |  | rs33340122 | N130D | B | 0.000 | 0.002 | -0.99 |
|  | *Cilp2* | cartilage intermediate layer protein 2 | rs33430594 | G486R | B | 0.060 | 0.098 | -4.43 |
|  |  |  | rs33216225 | S219F | B | 0.030 | 0.022 | -3.23 |
|  |  |  | rs3668298 | L94Q | B | 0.010 | 0.007 | -2.25 |
|  | *D13Rik* | RIKEN cDNA D130040h23 gene | rs32634767 | K213E | B | 0.030 | 0.009 | -2.70 |
|  | *Babam1* | BRISC and BRCA1 A complex member 1 | rs32539724 | A315S | B | 0.040 | 0.028 | -1.59 |
|  | *Zfp709* | zinc finger protein 709 | rs32828884 | L532F | B | 0.010 | 0.002 | -1.20 |
|  | *Olfr374* | olfactory receptor 374 | rs50830855 | S190L | B | 0.000 | 0.010 | -5.37 |
|  | *Olfr372* | olfactory receptor 372 | rs32685508 | H227L | B | 0.020 | 0.103 | -3.78 |
| *Canq7<sup>f</sup>* | | <sup>a</sup>Totals: 65 genes; 0 nonsense SNPs; 126 missense SNPs; 15 missense SNPs predicted deleterious by $\geq 1$, 9 by $\geq 2$ algorithms | | | | | | |
|  | *Cav2* | caveolin 2 | rs33540906 | D138G | W | 0.080 | 0.056 | -3.40 |
|  | *Iqub* | IQ motif and ubiquitin domain containing | rs263632219 | I514L | W | 0.080 | 0.023 | -1.69 |
|  |  |  | rs48630797 | D586E | C<sup>g</sup> | 0.000 | 0.006 | -3.30 |
|  |  |  | rs47850186 | E482Q | C<sup>g</sup> | 0.060 | 0.027 | -2.39 |
| *Canq8* | | <sup>a</sup>Totals: 177 genes; 1 nonsense SNP; 104 missense SNPs; 18 missense SNPs predicted deleterious by $\geq 1$, 12 by $\geq 2$ algorithms | | | | | | |
|  | *Pld1* | phospholipase D1 | rs29602715 | R328G | A | 0.010 | 0.014 | -2.64 |
|  |  |  | rs8247032 | R1030H | C<sup>h</sup> | 0.010 | 0.001 | -0.27 |
|  | *Fat4* | FAT atypical cadherin 4 | rs30811021 | A4900V | A | 0.040 | 0.016 | -1.87 |
|  | *Usp13* | ubiquitin specific peptidase 13 (isopeptidase T-3) | rs29854159 | T490M | A | 0.040 | 0.002 | -3.64 |
|  | *Gm1527* | Predicted gene 1527 | rs50586704 | R47H | A | 0.010 | 0.008 | -1.40 |
| *Canq9* | | <sup>a</sup>Totals: 82 genes; 0 nonsense SNPs; 73 missense SNPs; 14 missense SNPs predicted deleterious by $\geq 1$, 7 by $\geq 2$ algorithms | | | | | | |
|  | *Arhgap28* | Rho GTPase activating protein 28 | rs224248434 | D72Y | W | 0.000 | 0.001 | -3.48 |
|  | *Rmdn2* | regulator of microtubule dynamics 2 | rs108705699 | H24R | W | 0.010 | 0.016 | -6.43 |
|  |  |  | rs29747828 | S217C | W | 0.010 | 0.093 | 0.57 |
|  |  |  | rs29747826 | L251F | W | 0.020 | 0.018 | -1.00 |
|  | *Myom1* | myomesin 1 | rs51572930 | P1004T | W | 0.010 | 0.018 | -4.78 |
|  | *Ankrd12* | ankyrin repeat domain 12 | rs262070481 | V1737G | W | 0.010 | 0.001 | -0.78 |
|  | *Xdh* | xanthine dehydrogenase | rs47723242 | R945Q | W | 0.010 | 0.023 | -2.17 |
|  |  |  | rs29522348 | V241I | A<sup>i</sup> | 0.000 | 0.000 | -0.93 |
|  | *Mtcl1* | microtubule crosslinking factor 1 | rs47445501 | P300T | W | 0.040 | 0.007 | -2.12 |
| *Canq10* | | <sup>a</sup>Totals: 114 genes; 0 nonsense SNPs; 141 missense SNPs; 13 missense SNPs predicted deleterious by $\geq 1$, 12 by $\geq 2$ algorithms | | | | | | |
|  | *Tspyl2* | TSPY-like 2 | rs29296591 | E609K | 1 | 0.000 | 0.003 | -0.51 |
|  | *Foxr2* | forkhead box R2 | rs29289475 | E152V | 1 | 0.050 | 0.053 | -2.11 |

Collateral phenotyping for collateral number and diameter was performed at 6 weeks-age unless indicated otherwise. Preliminary studies showed that new collaterals can be induced to form, in addition to those that form naturally late during gestation, by intravenous (periorbital) injection of rAAV9-*Rabep2* into newborn (P0) *Rabep2^-/-^* mice, and that they mature by 6 weeks-age to yield the number and diameter present in the 10 week-old adult (see Results)—findings which confirm our previous identification of the interval when cerebral collaterogenesis occurs and native pial collaterals mature (E16.5 to P21).^19–21^ Access of AAV9 to the brain and its vasculature is aided by the blood brain barrier remaining open through post-natal day-0.^43^

### Quantitative RT-PCR

Total RNA was isolated from lung using the Qiagen RNeasy Kit per manufacturer’s protocol. 0.5 ug of RNA was processed with SuperScript^TM^ IV Reverse Transcriptase (#18090050, Invitrogen) following the manufacturer’s instructions. Amplification of targets was by Taqman assays on the QuantStudio^TM^ 7 RT-PCR system (Taqman IDs: *Rabep2* Mm00518884_m1, *Rgs1* Mm00450170_m1, *Gapdh* Mm 99999915_g1). *Gapdh* was used as the housekeeping gene. Three samples per group were analyzed in triplicate using the delta-delta method.

### Statistical analysis

Values are mean ± SEM. Statistical tests are given in the figure legends, with p < 0.05 denoting significance. Experiments were conducted according to the STAIR and ARRIVE guidelines:^44, 45^ This is an exploratory descriptive investigation using a forward genetics unbiased design, thus investigators were not blinded to strain/group; No data points were identified as outliers and excluded; The discussion and citation of the literature were unbiased; Number of mice per group were based on our previous studies which demonstrated sufficient power to test hypotheses regarding the variables measured.

## Results

### CC strains exhibit large differences in collateral number, diameter and stroke severity

Collateral number varied by 47-fold and anatomic (maximally dilated) diameter by 3-fold among the 60 CC strains (**Figure 1**). Both traits correlated weakly (r^2^ = 0.23, p < 0.001) (**Supplemental figure II**). These data agree with what we reported previously comparing 15 classical strains^18^ and 221 C57BL/6 x BALB/cBy F2 mice.^23^ Calculated heritability (as in Xing et al^46^) was 0.82 for number and 0.78 for diameter. To confirm the expected impact of this wide variation on stroke severity, we determined infarct volume 24h after permanent MCA occlusion for 6 CC strains with high, intermediate and low collateral number and diameter, as well as for B6 and BALB/cBy strains studied previously (**Figure 2A**). Strain CC044 with the greatest number of collaterals sustained—remarkably—no infarction, while strain CC031 with the largest diameter and a large number of collaterals evidenced a small infarction. Strain CC039, with number and diameter comparable to B6 mice, had a smaller infarction than the latter, confirming the well-known contribution of other factors, in addition to the abundance of collaterals, to cell loss.^25, 47^ Strain CC036 with the least number of collaterals sustained a large infarction similar to BALB/cBy mice that have comparable collateral number and diameter.^18, 47^ The number (and diameter, see figure legend) of PCA-to-MCA collaterals obtained for a subset of CC strains followed the same relative abundance as their ACA-to-MCA collaterals (Figure 2B). Collateral abundance in skeletal muscle of the abdominal wall and small intestine of selected high- and low-collateral strains evidenced the same relative abundance as in brain (Figure 2C,D). This agrees with our previous findings for classical mice.^12–16^

**Figure 1.**
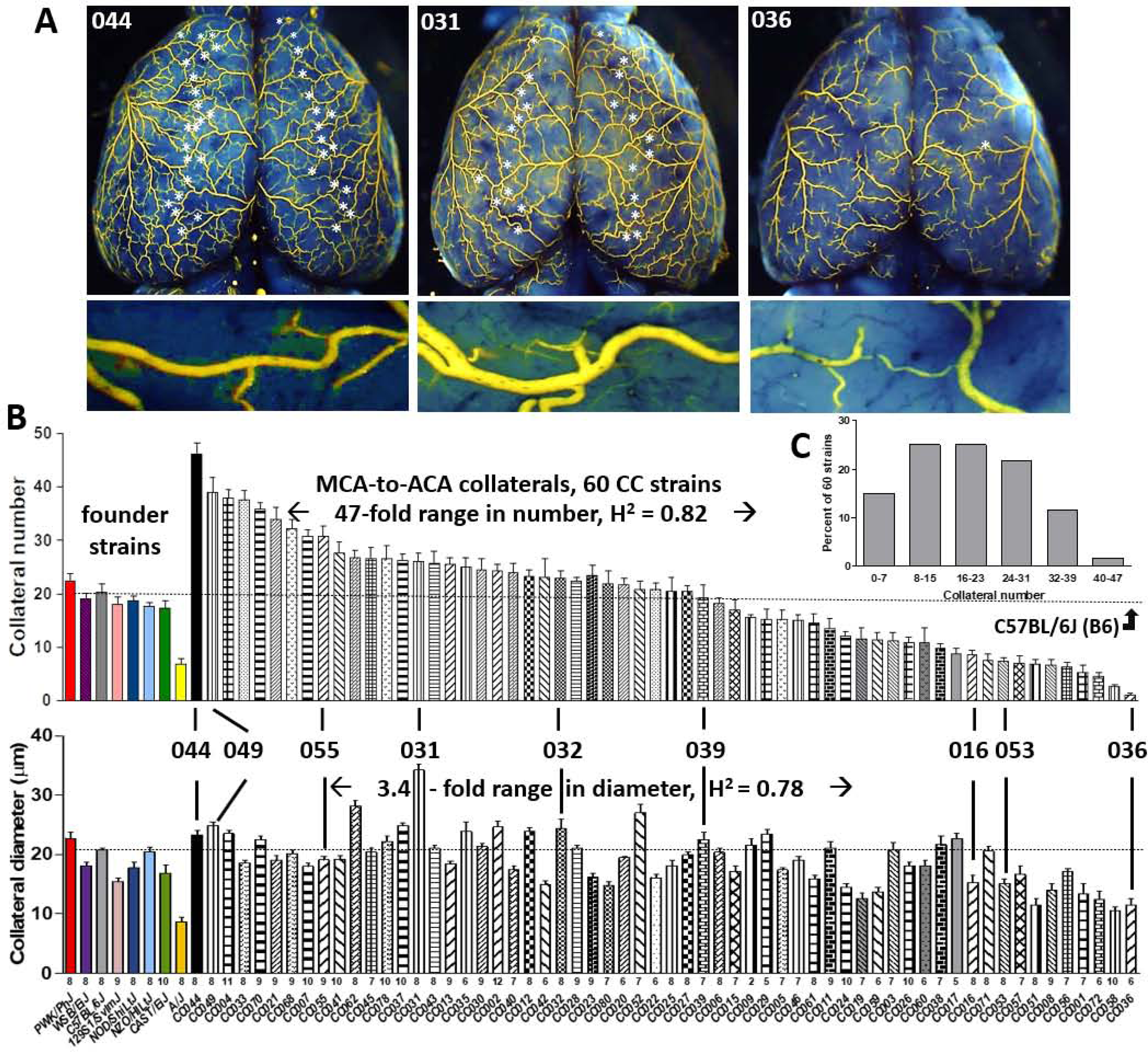
Wide variation in collateral number and diameter among Collaborative Cross strains. **A**, Collaterals between the right and left MCA and ACA trees of 3 strains (stars; representative collateral shown in lower panels). **B**, Strain names and number of animals per strain given below each bar (ẍ = 7.9 ± 0.2 per strain; 537 mice total). Strain CC044 has the largest number, CC031 the largest diameter, and CC036 the smallest number and diameter. CC049, CC055, CC053, CC036 were used in F2 crosses (Figure 4), CC036, CC032, CC016 in AAV9-expression assays (Figure 6). CC039 has number and diameter similar to B6 mice. Among the 60 CC strains: fold ranges and heritability values are given, distribution of collateral number is shown (**C**), and diameter correlates with number (r^2^ = 0.23, p < 0.001, Supplemental figure I).

**Figure 2.**
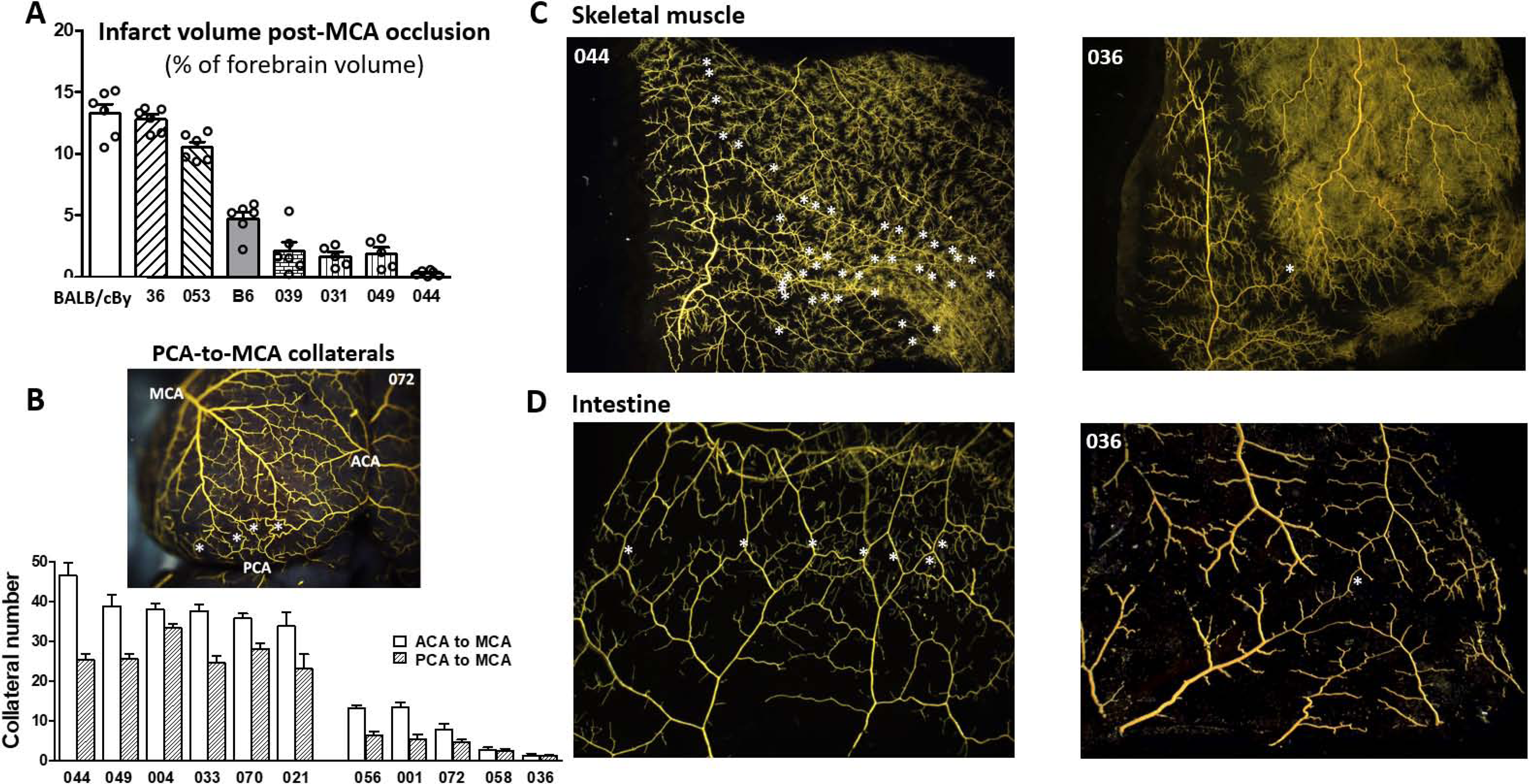
**Among selected CC strains: A, Stroke severity is inversely associated with collateral number and diameter; B, PCA-to-MCA collaterals (stars) have same relative strain-dependent differences in abundance as ACA-to-MCA collaterals; C, D, Collaterals (stars) in skeletal muscle and intestine have the same relative strain-dependent differences in abundance as in brain (Figure 1A).** A, Infarct volume determined 24h after permanent distal M1-MCA occlusion. B, Number of PCA-to-MCA collaterals of both hemispheres in the 6 CC strains with the highest and 5 strains with the lowest collateral number shown in Figure 1 (n = 5-11 per bar). Diameter of PCA-MCA collaterals for CC021 and CC036 (19.3 ± 1.0, 9.3 ± 0.8) do not differ from diameter of their ACA-MCA collaterals shown in Figure 1 (18.4 ± 0.5, 11.2 ± 1.3). C, D Angiographic images of abdominal wall skeletal muscle viewed from peritoneal side and small intestine.

### Quantitative trait loci for variation in collateral number and diameter

To identify QTL for collateral variation, we performed genome-wide linkage analysis using the available consensus haplotype reconstruction probabilities for the 60 CC strains (http://csbio.unc.edu/CCstatus/index.py?run=FounderProbs).^37^ Despite their wide range in collateral number and diameter, we did not identify any genomic loci reaching even suggestive (p<0.1) levels of significance (**Figure 3**). Importantly, we did not find any overlap with the loci we identified previously in a B6 X BALB/cBy F2 population that were on chromosomes (Chr) 7, 1, 3 and 8 (*Canq1, Canq2, Canq3* and *Canq4*).^23^ In that cross, *Canq1* was the major locus, and we narrowed it to the *Rabep2* gene using gene-editing (7:126 Mb)^22^. However in this study, we found no evidence for a major effect of *Rabep2* haplotype on collateral number (arrows, Figure 3). This is not wholly unexpected when comparing QTL for a highly polygenic trait mapped in the genetically diverse CC versus in an intercross of the closely related B6 and BALB/cBy strains.

**Figure 3.**
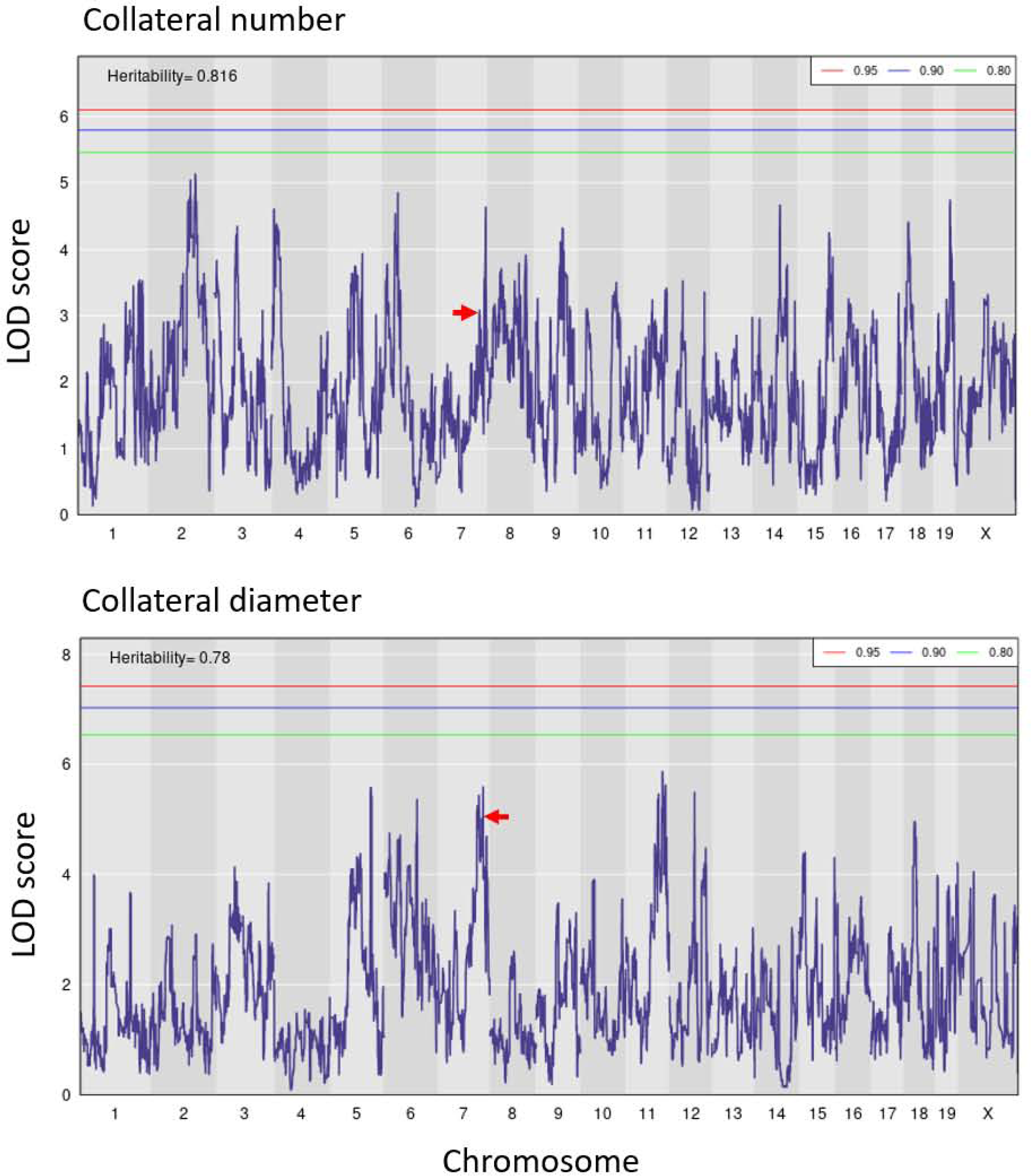
QTL mapping of collateral number and diameter in CC mice. LOD plots show the genomic position of QTL peaks for collateral number and diameter data in Figure 1. Relative chromosome length is shown by shading across the abscissae. Thresholds are for genome-wide significance based on 1000 permutations in this and subsequent figures. Eight peaks exceed LOD 4.5 for number and 8 for diameter; none is centered on the same marker SNP for both traits. Red arrows, peaks coincide with location of *Rabep2*.

Given the small trait-range for collateral diameter, together with the greater influence of collateral number than diameter on the distribution of collateral blood flow following permanent MCA occlusion,^47^ the remainder of our investigation focused on identifying loci associated with differences in collateral number.

The data in Figures 1 and 3 indicate that variation in collaterals is a highly polygenic trait. To increase our ability to identify a subset of the causal loci, we generated eight F1 crosses of 12 CC strains selected for having high-versus low-collateral number in each strain-pair to simplify the genetic structure of the trait and also increase our sample size (ie, both power and locus interval refinement) (**Figure 4A,B**). The strains were also selected for having homologous alleles for *Rabep2* to strengthen detection of additional QTL by preventing re-identification of *Rabep2.*^14, 22^ Two of the eight crosses (Figure 4A,C) were selected for generation of F2 populations, based on their trait distributions (see Methods).

**Figure 4.**
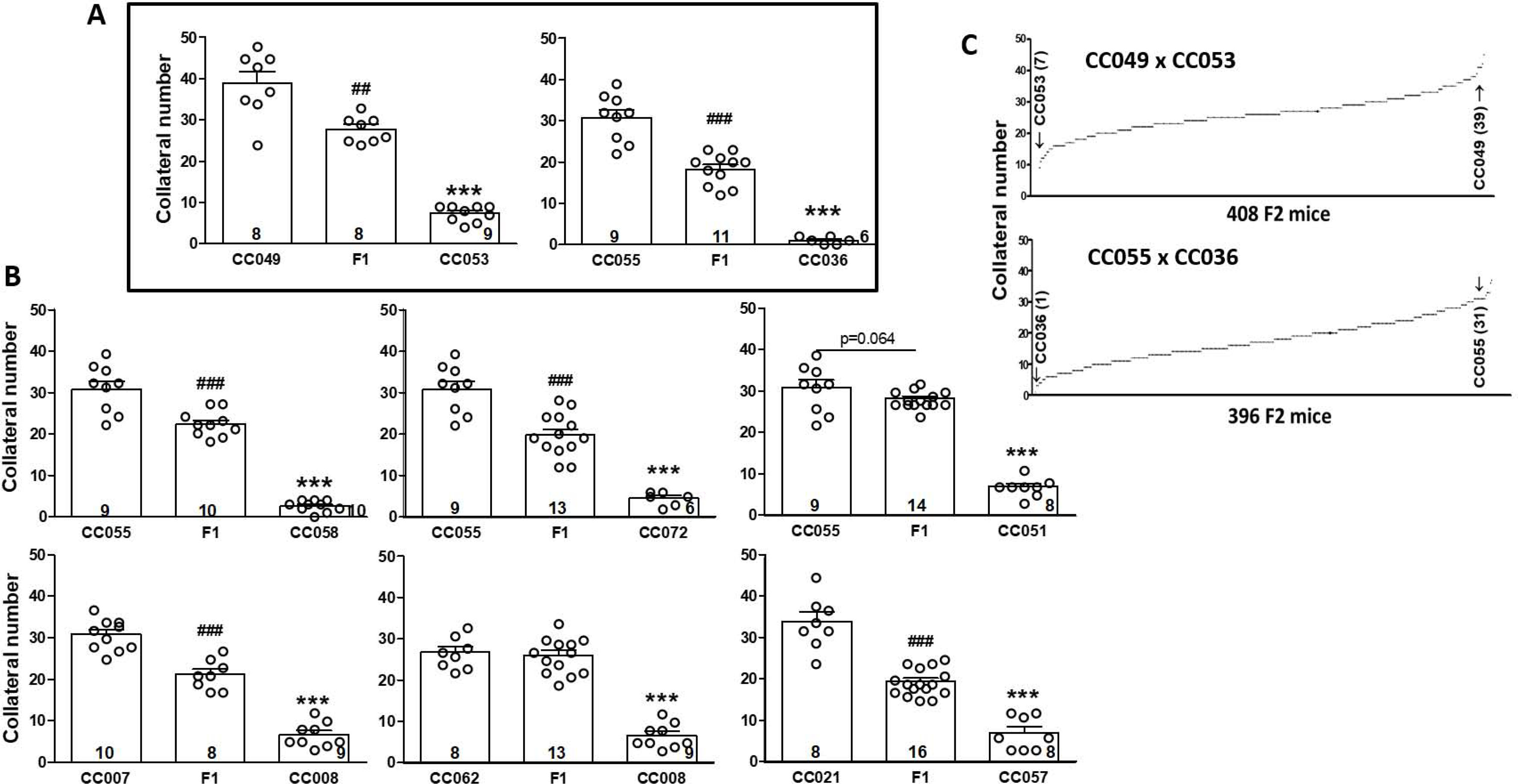
F1 crosses of 12 CC strains. Strains were chosen to give progeny with heterozygous genomes except at *Rabep2,* since variants of *Rabep2* were previously shown to cause large effects on collateral number and diameter in a panel of classical strains.^22^ **A**, F1 lines CC049 x CC053 and CC055 x CC036 were selected, among the 8 crosses (**A, B**), to generate two F2 populations (**C**) based on their *a priori* power ranking (see Methods) and minimum evidence of dominance. N-sizes in base of bars. ##, ### p<0.01, <0.001 vs. 1^st^ bar, ***, p<0.001 vs. 2^nd^ bar; 1-sided *t-*tests.

Concurrent with phenotyping the two F2 populations, we genotyped them using the MiniMUGA array.^36,see^ ^Methods^ For the CC049xCC053 intercross, we identified a single QTL on Chr 1 (128-157 Mb, **Figure 5A-C, Supplemental table I**) and named it *Canq5.* Additionally, seven significant (p<0.05) and two suggestive (0.10<p<0.05) peaks were identified in the CC055xCC036 population (**Figure 5D-F**, Supplemental table I). Given this large number of loci, we conducted a variety of analyses to confirm the loci (see Methods). We also estimated the phenotypic effects of each QTL, ie, the number of collaterals determined by each locus expressed as a percentage difference in collateral number between the parental strains: 15% for *Canq5* (4.7/31.0 collaterals, Figure 5C), 19% for *Canq6* (Figure 5E,F), 20% for *Canq7*, and 13%, 13%, 5%, 14% and 12%, respectively, for *Canq8-Canq12*. *Canq8* does not align with *Canq3* identified previously.^23^

**Figure 5.**
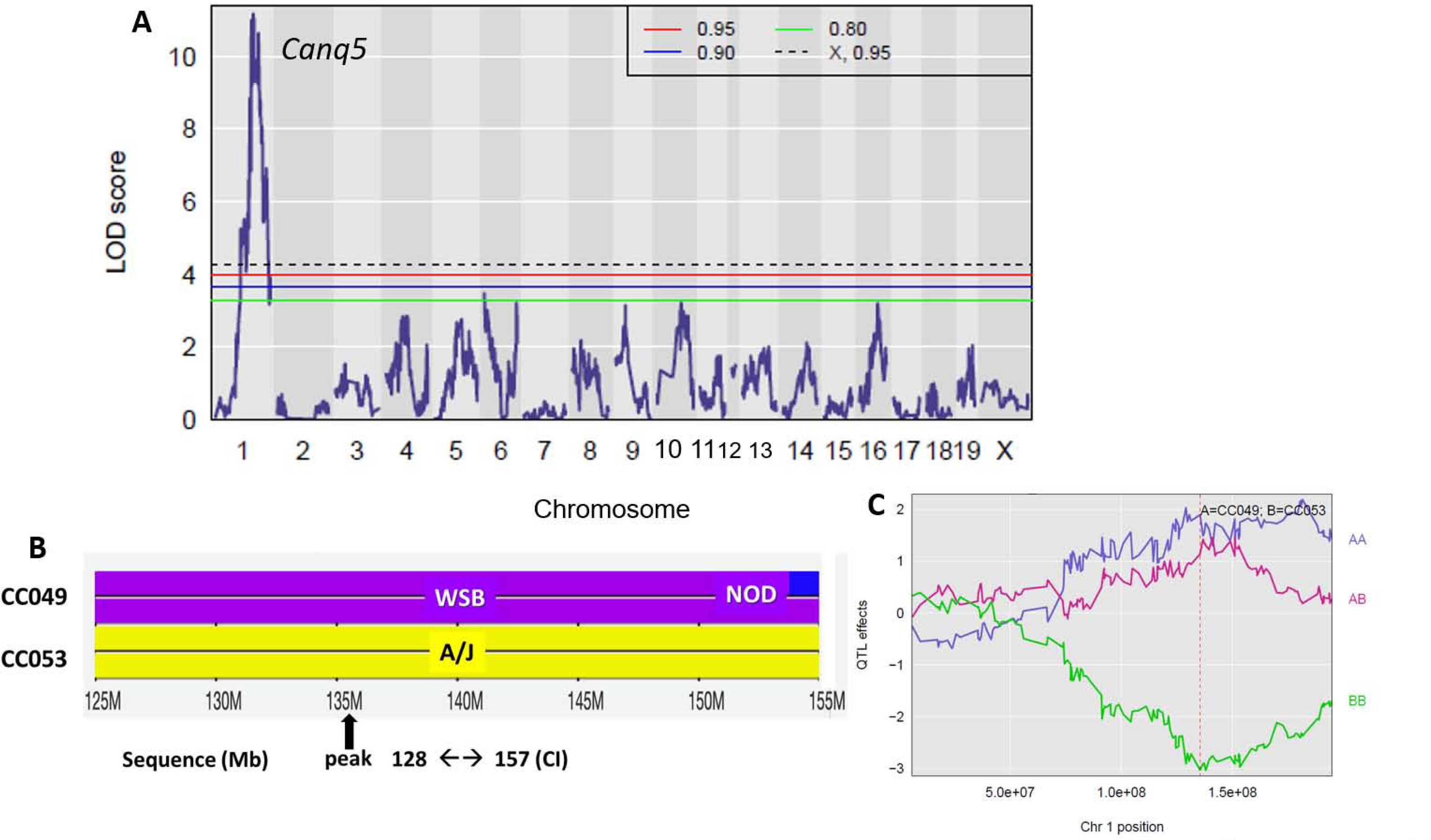
Mapping of CC049 x CC053 F2 progeny identifies a significant QTL on chromosome 1. **A**, LOD plot. **B**, Recombination blocks, peak, and confidence interval (CI) of *Canq5*. **C**, The *Canq5* allele accounts for 15% of the difference in collateral number between CC0049 and CC053 shown in Figure 4 (4.7/31 collaterals). Supplemental table I gives additional characteristics of *Canq5.*

**Figure 5.**
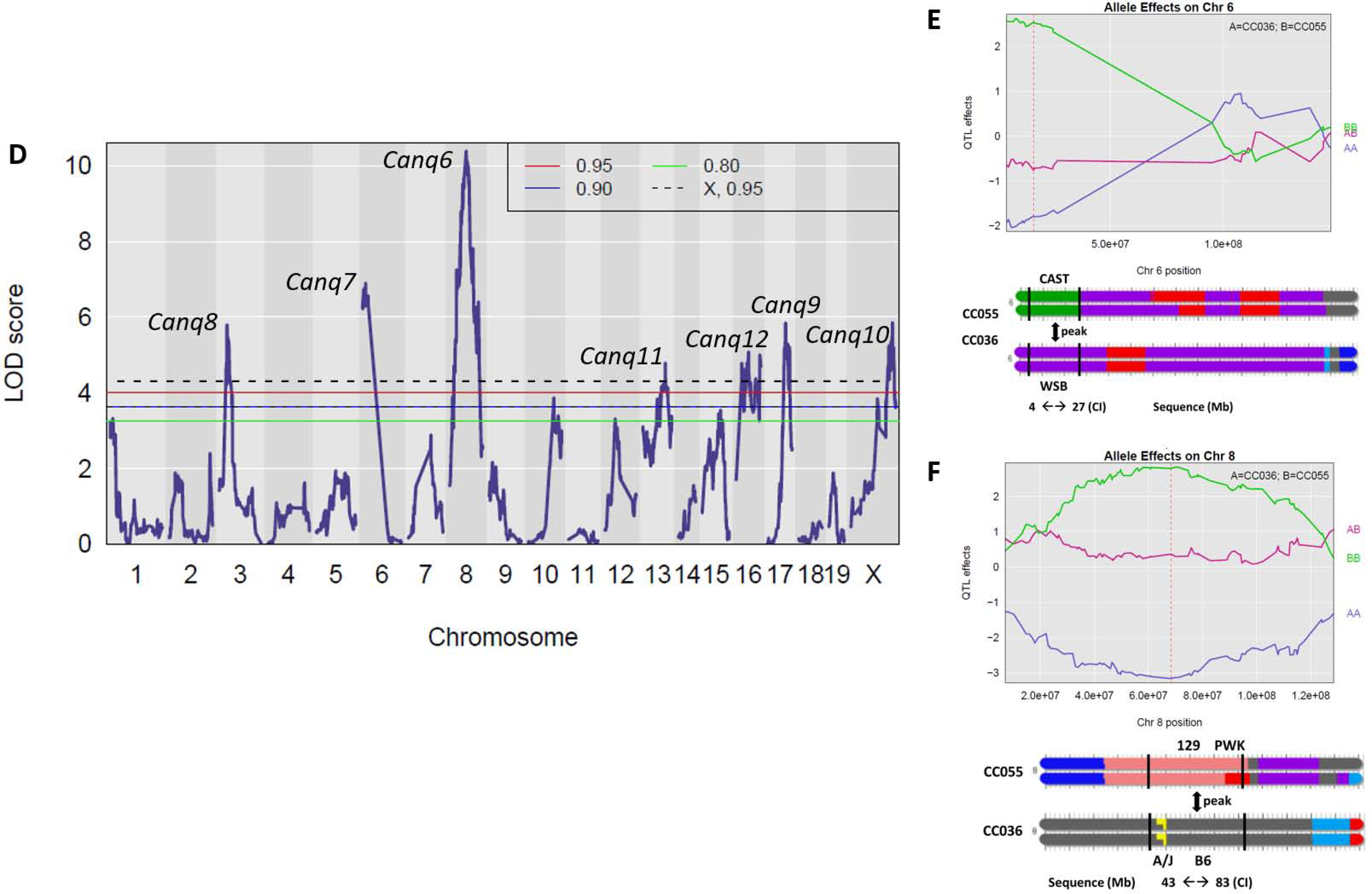
Mapping of CC055 x CC036 F2 progeny identifies 7 significant QTL. **D**, LOD plot. **E, F,** Recombination blocks, peaks, confidence intervals (CI) and allele effects of *Canq7 and Canq6*. Their alleles account for 19% and 20%, respectively, of the difference in collateral number between CC055 and CC036 shown in Figure 3. Supplemental table I gives additional characteristics of *Canq6-12*.

### Identification of candidate genes for variation in collateral number

Unlike crosses between classical inbred strains, crosses between CC strains can (and likely often do) point to large haplotype blocks arising from different mouse subspecies. Therefore, in addition to a large number of non-coding (intronic, intergenic, untranslated regions) variants that might alter regulation of genes, many genes within a QTL interval may possess one or more missense/non-synonymous SNPs (amino acid substitution/coding SNPs), as well as an occasional nonsense SNP(s) or insertion(s) or deletion(s) that create a premature stop-gained/lost codon. Such a variant(s) may have a functional effect (ie, be causal) for the QTL. We defined, *a priori*, candidate genes for proteins potentially involved in collaterogenesis according to the hypothesis that a deficiency in the protein’s function that is caused by a LoF variant(s) results in impaired collaterogenesis, ie, a reduction in the number of collaterals that form during development, resulting in a reduced collateral number in newborns and thus adults of the low-collateral strain of our F1 intercross populations. Therefore, we identified all protein-coding genes within ±10 Mb of the peak for each of *Canq5-10. Canq11* and *Canq12* were not investigated because their peaks exhibited lower LOD scores and poor differentiation from surrounding signal (Figure 5). We then identified nonsense and missense SNPs segregating between each pair of strains used in the cross (based on the sequences of the founder strains contributing haplotype blocks to these CC strains) within the above genes. Lastly, we subjected these missense and nonsense SNPs to three in silico prediction algorithms (see Methods) to determine their potential for deleterious effects on protein function.

For *Canq5* we ascertained the following genes with protein-affecting variants (see “Totals” in **Table 1**): 150 genes are present within ±10 Mb of *Canq5’s* peak that among them harbor 3 nonsense and 180 missense SNPs between CC049 (high-collateral WSB allele) and CC053 (low-collateral A/J allele). Thirty-seven of the missense SNPs were predicted to be deleterious by at least one and 20 by at least two prediction algorithms. Among these, Table 1 lists the 6 top candidate genes underlying *Canq5* ranked highest-to-lowest based on: 1) presence of the nonsense or missense SNP(s) in the haplotype block of the low-collateral parental strain; 2) pLoF overall p-value among the 3 prediction algorithms for the given SNP; 3) evidence in the literature (see “Bioinformatics analysis” in Supplement) linking the gene’s function to one or more of the following Medline search terms: angiogenesis, vascular development, endothelial cell (EC) function, endocytic vesicle trafficking (per that *Rabep2—*the only variant gene thus far shown to be involved in collaterogenesis^22^—is involved in this process). Table 1 also lists the top 22 genes, ranked as above, that underly *Canq6-Canq10*, and their associated data. **Supplemental table II** lists genes underlying *Canq5-10* with nonsense or missense SNPs in the haplotype block of the high-collateral parental strain. These were not viewed as high-priority candidate genes because they did not fit our above-stated *a priori* hypothesis (see also below).

### Development of an in vivo collaterogenesis assay

Identifying genes underlying QTL within the CC for involvement in collaterogenesis presents a number of difficulties: Knockouts, floxed alleles, other types of spontaneous mutations, and transgenic mice have been constructed or identified most often in B6 and otherwise infrequently in other classical strains and used to identify causal genes underlying QTL, including *Rabep2* underlying *Canq1*.^22^ However, such genetic resources are not available for CC mice. Since the findings in Figure 3 indicate that collaterogenesis is a polygenic process wherein multiple loci contribute to collaterogenesis, the impact of a causal gene varies depending on genetic background. As such, it was not feasible to address this dilemma by creating the dozens of ubiquitous and/or cell-specific conditional KOs in the high-collateral parental CC strains (CC049 and CC055) or transgenic mice in the low-collateral parentals (CC053 and CC036) that would be needed to screen the genes listed in Table 1. Even if this were feasible, breeding difficulties encountered with some of these strains (discussed below) present significant impediments for creating gene-targeted mice in a timely manner. In addition, collaterogenesis in the mouse neocortex—wherein collateral number (and diameter) can be robustly phenotyped—occurs late during gestation (E15.5-18.5, E20-21=birth) in the subarachnoid space above the pial/leptomeningeal membrane that overlies the watershed regions between adjacent arterial trees.^19–21^ The process involves proliferation of tip- and stalk-like ECs that originate from distal arterioles and migrate across the watershed regions to cross-connect adjacent arterial tree branches, followed by lumen formation. There is no method—and we were unsuccessful in developing one—for in utero delivery of vectors carrying antisense or transgenic sequences for candidate genes to ECs and/or other cell types within the watershed regions during the above embryonic time interval to test for inhibition or augmentation of the number of collaterals that form (ie, collaterogenesis). Lastly, understanding of the collaterogenesis process is in its infancy, thus no in vitro model system has been devised.

Given these constraints, we sought to develop an in vivo assay for use in early postnatal mice to test the candidate genes underlying *Canq5-Canq10* based on two observations: First, collaterogenesis in mouse brain concludes just before birth, followed by collateral maturation during the first 4 postnatal weeks to yield the number and diameter that are present in the adult.^19–21^ Second, the blood brain barrier is open for the first ∼24h after birth in mice.^43^ We therefore postulated that intravenous administration on postnatal day-0 (P0) of an expression plasmid harboring the ORF of a gene in the collaterogenesis pathway carried by a virus with a broad range of cell-type infectivity, including brain ECs (eg, AAV9 used herein), may be capable of re-activating collaterogenesis to induce formation of additional collaterals—and thereby provide an assay to test our candidate genes. To examine this hypothesis, we tested *Rabep2*—a key contributor to collateral formation in B6 and a number of other classical strains^22^—as a “positive control”. AAV9-*Rabep2* (ORF=C57BL6/J sequence) was injected at P0 in B6.*Rabep2*^-/-^ knockout mice. As reported previously, these mice have 60% fewer collaterals with smaller diameters than wildtype mice.^22, 47^ We used AAV9 rather than lentivirus because the latter has narrower tropism for the multiple cell types likely involved in collateral formation, and because it directs incorporation of plasmid DNA into the host genome which creates a heightened risk to personnel when used in live animals.

Administration of AAV9-*Rabep2* (1×10^13^ vg/ml, 4 μl) on P0 increased collateral number by 37% (p<0.002) over control (ie, AAV9 carrying the identical plasmid construct but without the *Rabep2* ORF) when examined 4 weeks later (**Figure 6A**). Collateral diameter was also increased (**Supplemental figure III**). Values for control were not different from naïve *Rabep2*^-/-^ mice (data not shown) reported previously.^22, 47^ Administration of 2.5-fold higher AAV9-*Rabep2* (10 μl) followed by examination at 10 weeks-age did not cause an additional increase in collaterogenesis over that shown in Figure 6A: number=12.0±0.8, diameter=13.7±0.4 μm, n=7. Neither did 5-fold higher AAV9-*Rabep2* (20 μl) measured at 6 weeks-age: number=12.4±1.8, diameter=13.9±0.7 μm, n=9. These findings are consistent with our previous evidence that collateral formation and maturation are complete by 4 weeks-age.^19–21^ They also demonstrate that maximal effect is achieved at the lower dose, with a 5-fold higher dose causing no adverse effect. Preliminary experiments also found that injection of AAV9-*Rabep2* on P1 resulted in formation of fewer and at P2 or later no additional collaterals over control, consistent with the time-course for blood brain barrier maturation.^43^ We also examined AAV9-mediated *Rabep2* expression in lung of *Rabep2^-/-^* mice because of its high proportion of ECs (the primary cell type involved in collaterogenesis^19–22^) and accessibility for rapid removal into liquid nitrogen without manipulation injury and RNA degradation. qPCR of *Rabep2* at 6 weeks-age after administration of 10 μl of AAV9-*Rabep2* at P0 increased expression 3.5-fold (0.91±0.11 control AAV, 3.17±0.71 *Rabep2-*AAV9; n=3 each; note the above control/baseline expression value reflects amplification of the truncated nonsense-mediated decay transcript expressed in the CRISPR-derived *Rabep2* knockout mouse^22^). The above results affirm the hypothesis stated in the preceding paragraph and the use of our assay to assess candidate genes in the low-collateral F2 parental strains (Table 1). Based on them, we adopted a dose of 10 μl (1 x10^13^ vg/ml titer) and assessment at 6 weeks-age for all tests, with the assumption that the ORFs of the Table 1 genes were expressed at levels similar to the above for *Rabep2* (∼3.5-fold) since all were inserted into the same plasmid construct with identical promoter and other regulatory and insertion elements (see Methods); we confirmed this for *Rgs1* (4.9-fold increase, see below). An exception was *Jak3*; the WPR cassette was not included in its construct because its ORF of 1100 is slightly above borderline for AAV9 cargo size. To adjust for this, we injected 10 ul of higher titer (3.4×10^13^; n=13) and also tested a 3-fold higher concentration (n=4); both doses were without effect (see below).

**Figure 6.**
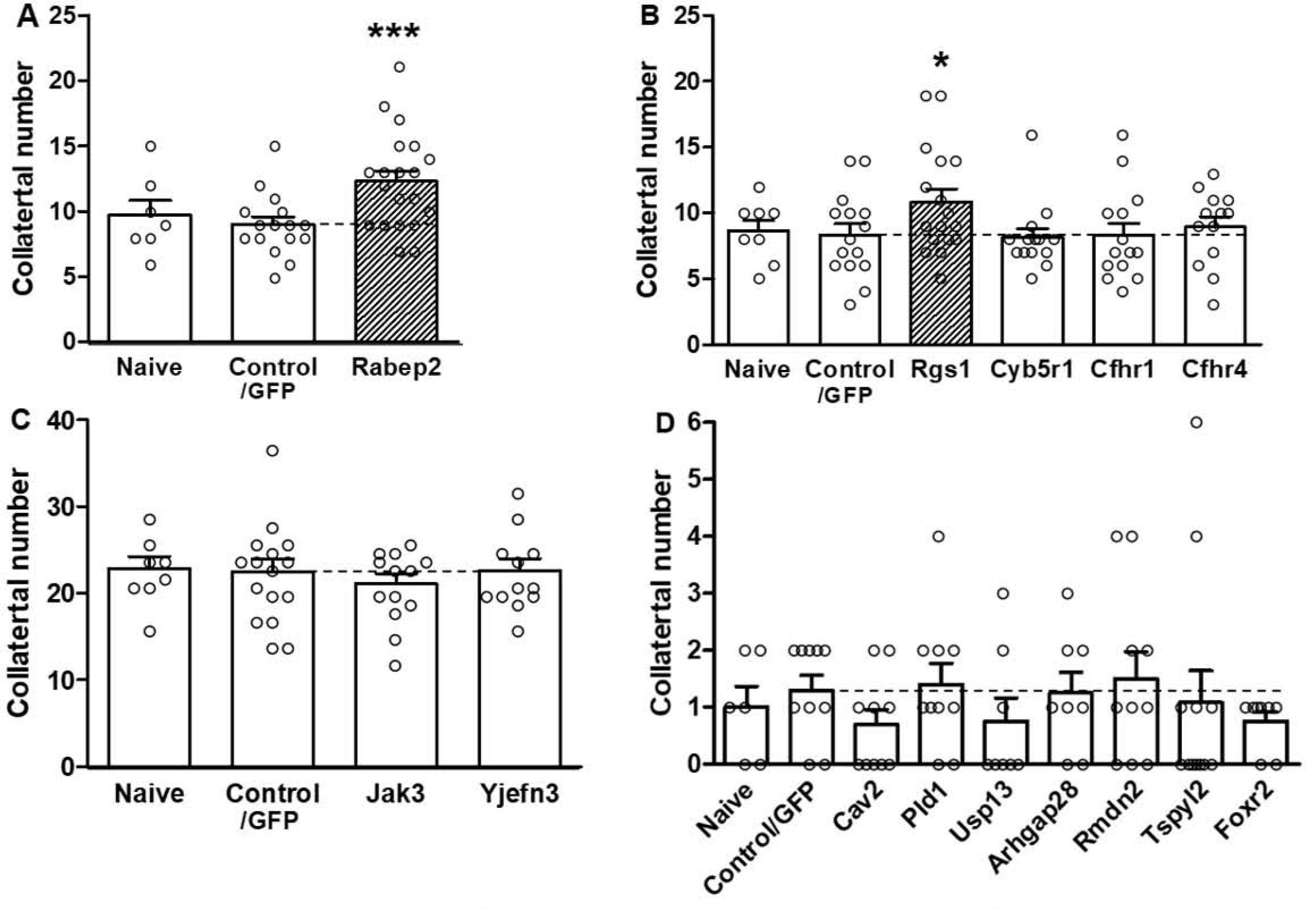
In vivo analysis of candidate genes. AAV9-expression constructs for *Rabep2* (**panel A**, control for the assay) and selected high-priority candidates underlying *Canq5* (**B**), *Canq6* (**C**) and *Canq7-10* (**D**) were injected iv on postnatal day-zero before blood brain barrier closure into *Rabep2^-/-^*(A), CC016 (B), CC032 (C) and CC036 (D) mice. Number of collaterals (Supplemental figure III gives diameter) between the MCA and ACA trees of both hemispheres were measured at 6 weeks-age. Naïve, no injection; Control/GFP, injection of AAV9 construct for enhanced green fluorescent protein only. Number of animals for each bar: Panel A: 7,16,20; B: 8,15,17,14,14,13; C: 8,16,13,12; D: 6,10,7,10,8,8,10,9,8. *p<0.05, **p<0.01, ***p<0.002 vs. Control/GFP by 1-sided *t*-tests.

Unexpectedly, difficulty was encountered in obtaining the many hundreds each of CC036 and CC053 PO pups needed for the above assays: CC036 trios were slow to yield pregnant females, the number of pups/litters was small (mean ∼2.5 in successful pregnancies), dams killed their pups or provided inadequate nurturing—which was not remediated using foster-dams but improved after a given dam’s second pregnancy—and females stopped breeding after the fourth pregnancy. Many CC032 pups (the rationale for use of this strain is given below) proved fragile and died shortly after injection for unknown reasons. Thus, litters yielded at most 3 pups at P0 that gained weight normally after injection, which qualified them for phenotyping. While the above difficulties made testing *Canq7-10* candidates a protracted 3-year process, a greater difficulty was encountered: CC053 mice were exhibiting poor breeding after completing the crosses in Figure 4 that progressed to breeding failure (extinction) by the time we had completed developing the above assay. In fact, many CC strains among the more than 500 original CC strains exhibited poor breeding vigor compared to classical strains, leading to extinction due to incompatibilities between alleles affecting fertility among the 8 founder strains’ different subspecific origins.^48^ Extinction of CC053 required us to substitute the CC016 strain in the assays of *Canq5* genes, based on it having adequate breeding vigor, a similarly low collateral number (Figure 1), the same haplotype block at *Canq5* as CC055, and the closest agreement in haplotype structures for the *Canq6-Canq10* intervals to CC053 among the extant CC strains shown in Figure 1.

An additional difficulty was encountered in the evaluation of candidate genes for *Canq6*: Only ORFs for the B6 mouse “reference” strain are available for construction of expression plasmids. This was not a problem for testing genes underlying *Canq5* and *Canq7-10* because, for a given deleterious SNP listed in Table 1, the B6 strain shared the same allele as the high-collateral parental strain in the F2 crosses yielding *Canq5* and *Canq7-10* [CC049 and CC055 (and its CC032 substitute—see below), respectively] (Figure 5, Table 1, Supplemental table I). In contrast, the alleles for the low-collateral deleterious SNPs within the candidate genes underlying *Canq6* were conferred by the B6 founder strain (Figure 5, Table 1). This difficulty required that the *Canq6* candidates be tested with AAV9 constructs with B6 ORFs for a given candidate in the CC055 high-collateral strain according to the hypothesis that the causal gene(s) will reduce collateral number. However, like CC053, CC055 had undergone extinction by the time assay development was completed. This required us to substitute CC032 for CC055, based on the same criteria given above for CC016 (including in this case that the high collateral number in CC032 is similar to CC055, Figure 1).

### Regulator of G-protein signaling 1, *Rgs1,* is a potential causal gene for *Canq5*

Administration of control AAV9 (ORF for *GFP* only) to CC016, CC032 and CC036 mice at P0 had no effect on collateral number when examined at 6 weeks-age when compared to untreated mice at the same age (“naïve”, **Figure 6**), confirming our above-mentioned lack of effect of AAV9 or the injection procedure per se. Among the candidates underlying *Canq5* for low collateral number (Table 1) only AAV9-*Rgs1* increased collateral number. No additional effect occurred in mice receiving a 3-fold higher dose (n=3), in agreement with that observed for AAV9-*Rabep2*. As expected, qPCR of *Rgs1* in lung of 6 weeks-old CC016 mice after administration of 10 μl of AAV9-*Rgs1* at P0 increased expression 4.9-fold (3.85±1.66 control AAV, 19.16±3.04 *Rgs1-*AAV9; n=5 each), indicating similar efficacy of the vector for *Rgs1* as seen for AAV9-*Rabep2*. AAV9 for *Cyb5r1, Cfhr1 and Cfhr4* were without effect. We were unable to evaluate *Sox13* because all 19 treated pups died between P2-P16 for unclear reasons (autopsies could not be performed at or close to the time of death). *Aspm* was not tested because its ORF (3122 amino acids) exceeded AAV9’s maximum cargo size. Lentivirus can accommodate larger plasmids but was not used for the reasons stated earlier.

### Testing candidate genes underlying *Canq6-Canq10* did not identify potential causal genes

AAV9-*Jak3* had no effect on collateral number at the standard dose. Given evidence for its involvement in angiogenesis (see “Bioinformatics analysis” in Supplemental material) and for other reasons mention above, we tested a 3-fold higher dose which also was without effect (n=10). AAV9-*Yjefn3* also had no effect. CC036 mice treated with AAV9 for *Canq7-9* candidates *Cav2, Pld1, Usp13, Arhgap28, Rmdn2, Tspy12 and* Foxr2 did not evidence a change in collateral number (Table 1, Figure 6C,D). For unclear reasons, AAV9-*Cav2* caused pups to die immediately after injection (n=6). A 3-fold lower dose, which had no effect on viability, did not affect collateral number (n=7). We did not test the last 6 genes listed in Table 1 for *Canq6* nor *Iqub*, *GM1527, Ankrd12, Xdh* and *Mtcl1* underlying *Canq7-10* because each was deemed unlikely to be involved in collaterogenesis based on the following: highly restricted function and expression in cell types not involved in collaterogenesis (see “Bioinformatics analysis” in Supplement); and no references were returned by the Medline search terms: angiogenesis, vascular development, endothelial cell function, and endocytic vesicle trafficking. In addition, *Iqub* has two predicted LoF SNPs in the high-collateral strain, CAST (Table 1), which would confound interpretation of results, and the ORFs for *Cilp2, Fat4* and *Myom1* (1162, 4981 and1667 amino acids) exceeded the size-limit for AAV9.

**Supplemental table II** lists genes underlying *Canq5-10* with LoF SNPs in the haplotype block of the high-collateral parental strain for the crosses shown in Figure 4. One might a priori nominate these genes as potential “anti-collaterogenesis genes” that restrain collaterogenesis according to the hypothesis that a missense or nonsense mutation results in impaired protein function or amount, leading to increased collaterogenesis and thus collateral number in the adult. However unlike the genes in Table 1, we deemed these not high-priority candidate genes and thus did not conduct AAV9-expression assays for them because there is no evidence for—and it is unlikely that—collateral formation during development relies on “anti-”collaterogenesis genes/signaling pathways providing a restraint on “pro”-collaterogenesis genes (see Discussion), and because the extensive effort required to test the pro-collaterogenesis gene candidates in Table 1 made doing so unfeasible.

The data in Figures 3 and 5 show that collateral number is a polygenic trait where contributions of the individual causal variants differ according to genetic background. It is therefore unlikely that examining B6 mice with a targeted mutation for a gene listed in Table 1 or Supplemental table II will be informative. Nevertheless, we examined two high-priority candidates using this approach. *Cyb5r1* is among the top candidates for *Canq5*. Because deletion of B6.*Cyb5r1* is embryonic lethal, we obtained floxed B6 mice and crossed them with ubiquitous and EC-specific *Cre* mice (see Methods). Unfortunately, pups were dead on inspection post-delivery from unknown cause. We also examined *Igfn1*, the highest ranked gene for *Canq5* in Supplemental table II. Collateral number did not differ among 8-week old *Igfn1*^-/-^, *Igfn1^+/-^* and C57BL6/NJ wildtype mice: 19.6±1.5 n=11, 19.0±0.8 n=7, 18.9±1.5 n=9.

**Supplemental table III** lists genes underlying *Canq5-10* that lack nonsense or missense SNPs but have functions known to be involved or associated with collaterogenesis, angiogenesis, vascular development, endothelial cell function, or endocytic vesicle trafficking and thus could affect collaterogenesis depending on level of expression. Owing to logistical limitations, we confined testing to only *Cxcr4* and *C1galt1*, the highest ranking genes for the two largest QTL, *Canq5* and Canq6, according to the hypothesis that a mutation in a regulatory element(s) within the QTL interval of either gene in the low-collateral strain results in reduced expression leading to reduced collaterogenesis, suggesting that the gene is causal. Treatment of CC016 P0 pups with AAV9-*Cxcr4* was without effect on collateral number (control=8.3±0.8, n=9 AAV9-*Cxcr4*=7.0±0.9, n=15) or diameter (control=12.6±1.2, n=9; AAV9-*Cxcr4*=14.5±0.5, n=15). Since deletion of *C1galt1* is embryonic lethal,^48^ we obtained B6 mice with the floxed allele (B6.*C1galt1*) and crossed them with EC-specific *Cre* mice (Methods). Unfortunately and for unknown reasons, 4 litters from different breeder trios yielded cyanotic pups on P0 that did not survive to P1. For completeness, collateral diameter was also determined in the above assays (**Supplemental figure III**). Consistent with their effect to increase collateral number (Figure 7), *Rabep2* and *Rgs1* also increased collateral diameter (p<0.01; n=20, n=17, respectively); other genes were without effect, except *Cfhr4* which increased diameter for unclear reasons based on the literature (see Bioinformatic analysis in Supplement). Lastly, we performed in silico analysis of the human orthologs of the above genes and provide a comprehensive list of their ancestry-associated variants predicted to be damaging (**Supplemental table IV**).

## Discussion

The Collaborative Cross genetic reference panel can be viewed as a cohort of 60 healthy young-adult individuals with widely diverse genetic backgrounds. Despite that collateral number and diameter varied by 47-fold and 3-fold, we were unable to identify any significant large-effect QTL in a population-wide analysis (Figure 3). This indicates that collateral abundance is a highly polygenic trait. Collateral number and diameter correlated weakly with each other, and none of the 8 non-significant QTL with LOD > 4.5 for number and diameter were co-localized. This suggests that many of the polymorphic genes responsible for differences in collateral number and diameter are not shared, ie, that the signaling pathways and mechanisms that determine them differ. This is plausible since collaterogenesis and collateral maturation are distinct processes that occur at different times (see Introduction and Results).^19–21^ Unlike collaterogenesis, collateral maturation is likely specified by the same or a similar pathway activated by fluid shear and circumferential wall stresses that drives the increase in diameter of arteries and arterioles that occurs during postnatal tissue growth.^50, 51^

The values we calculated for heritability for variation in collateral number and diameter among the CC (82 and 78 percent) probably underestimate the contribution of genetics, since measurement error is essentially absent and environmental and stochastic variation during development can be assumed to be the same among the strains. Also, collateral number and diameter do not correlate with body weight, brain weight, or sex.^18, 23, 52^ Collateral number in the 10 week-old CC founder strains (Figure 1) compare with those reported by Lee et al^25^ for 3 week-old founders, with the exception that number was ∼25% greater in WSB/EiJ and CAST/EiJ mice in their study. This may reflect that postnatal pruning of a fraction of the newly formed collaterals, which is complete by 3-4 weeks in C57BL/6J and BALB/cByJ mice^19^—and apparently also in PWK/PhJ, 129S1/SvlmJ, NOD/ShiLtJ and NZO/HlLtJ mice since our and Lee et al’s values for these strains agree—presumably had not reached completion in their 3 week-old WSB/EiJ and CAST/EiJ mice.

As expected, infarct volume after permanent MCA occlusion correlated inversely with collateral number and diameter (Figure 2A). The phenotyping data in Figure 1 identify strains with high, intermediate, and low collateral number and diameter for use in future studies, for example respectively, CC044 or 049, CC027 or 039, and CC036 or CC058, that in contrast to similarly varying but closely related classical strains like B6, A/J, BALB/cBy,^13, 18^ have much wider variation in their genetic backgrounds. They therefore provide more genetically diverse models of the “outbred” human population (see UNC SGCF or Jackson Laboratories for information about obtaining specific animal numbers for CC strains). However a limitation of CC mice is that, like most classical strains other than the B6 (C57BL/6) reference strain for *Mus musculus*, almost no targeted deletions or transgenic modifications of genes are available for them. As well, genetic-dependent variation in factors in addition to collateral number and diameter exists among both CC and classical strains and contribute to variation in collateral blood flow. These include collateral length, perfusion pressure, viscosity, size of the occluded tree, and—in the dependent territory—dynamic changes in hydraulic resistance, leukocyte-platelet mechanisms, interstitial pressure, and neuro-glial sensitivity to hypoxia/ischemia.^18, 47^ ^and^ ^references^ ^therein^ This is supported by the observation that the inverse relationship between infarct volume and collateral number/diameter does not always track closely when comparing two strains with the same values for the latter (eg, B6 and CC039, Figures 1 and 2A, see also Lee et al^25^).

We previously found that 4 QTL account for almost all of the 20-fold variation in collateral number between B6 and BALB/cBy mice, and that the largest-effect QTL (*Canq1,* LOD 25) is responsible for ∼70% of this variation and is caused by a LoF variant in *Rabep2*.^22^ We were therefore surprised to find that no QTL, including none marking *Rabep2*, reached significance on mapping the 60 CC strain-set (Figure 3) despite its large 47-fold variation in collateral number and the robust sample sizes that were employed. However, in retrospect this is not altogether unexpected: Primary angiogenic proteins, including VEGF-A, Flk1, Notch, DLL4, EphrinB2, and EphB4 that direct formation of the capillary plexus and arterial-venous trees during development, are required to be fully functional and expressed at precise times and concentrations for normal fetal and postnatal growth.^53^ Accordingly, their genes are presumably under strong selective pressure against damaging alleles that would reduce protein activity or expression enough to impair formation of the correct vessel densities, diameters and branch-patterning that are required to optimize hydraulic conductance within tissue vasculatures. In contrast, deleterious polymorphisms within the primary genes and regulatory elements within the collaterogenesis pathway are less likely to be under such negative selection pressure. We hypothesize this based in part on the fact that cohorts of BALB/cBy^12, 16–18^ and CC036 (Figure 1A,B, 2B-D) mice—which have on average one collateral and will include a fraction with no collaterals in their brain and other tissues—evidence no apparent developmental, anatomical or physiological limitations (cardiovascular or otherwise) compared to strains with abundant collaterals. This indicates that collaterals are not required to assure adequate blood flow to support fetal development, growth to adulthood, and tissue function under normal conditions (this may not be the case when oxygen availability is limited^15, 35, 54^). This conclusion is supported by the observation that little or no net blood flow occurs across the collateral network in the absence of arterial obstruction.^55^ ^and^ ^references^ ^therein^ In other words, the morphogenesis of the arterial-capillary-venous circulations across classical^12, 18, 24^ and CC strains (Figures 1A,2B-D) are presumably comparable since development and post-natal growth of their brains (and other tissues) are comparable, yet their average pial collateral number can vary by ∼50-fold.

That collateral number is a highly polygenic trait contrasts with the conclusion suggested by our previous studies^24^ that only a small number of QTL (polymorphic genes/genetic elements) are responsible for most of the variation. However, this conclusion was based on mapping within an F2 population of B6 and BALB/cBy mice^23^ and 16 additional classical strains^24^ that share up to ∼70 of their genomes in common.^27–29^ Indeed prior to the availability of the CC, mapping studies were confined to using these and other classical strains despite their inherent limited genetic diversity and blindness to a large fraction of the mouse genome. This explains why despite polymorphic *Rabep2*’s large contribution to variation in collateral formation between B6 and Balb/cBy, we did not find a QTL reaching significance over *Rabep2* (Figure 3). That is, while *Rabep2* is a primary causal gene for variation in collaterogenesis among B6, BALB/cBy and 16 other relatively closely related strains,^22^ other genes have more prominent roles in the genetically diverse CC and—presumably by inference—the mouse as a species.

Since no significant loci emerged on mapping the CC strain-set, we examined two F2 populations derived from four parental CC strains selected for their informative power. This was done to reduce the number of allelic combinations of polymorphic genes responsible for differences in collaterals between a pair of strains and thus increase the likelihood of identifying significant QTL. Among the 8 QTL thus identified, 6 had intervals that were sufficiently defined to allow an analysis of protein-coding genes within the 10 Mb interval surrounding the QTLs’ peaks for presence of predicted deleterious mutations. We thus identified 28 candidate genes, based on the assumption that the presence of a damaging polymorphism will lead to impaired collaterogenesis and contribute to the reduced collateral number phenotype in the low-collateral parental strain (Table 1). Among the 13 top candidates amenable to evaluation using our in vivo assay, Regulator of G-protein signaling 1 (*Rgs1*) emerged as a potential causal gene for the single large QTL, *Canq5*, identified on mapping the CC049xCC053 F2 population. However, no potential causal genes for the five QTL mapped for the CC055xCC036 population were identified on in vivo assay (see Limitations section below).

Rgs1 is a GTPase-activating protein that limits G_∝_ and G_β_γ signaling from certain G protein-coupled receptors (GPCRs).^56^ Little information exists regarding its role in vascular development.^57^, ^see^ ^also^ ^Bioinformatics^ ^analysis^ ^in^ ^Supplement^ Knockout/down of *Rgs1* reduced expression of VEGF, proliferation of ECs, and suppressed angiogenesis and neovascularization.^58, 59^ These reports are consistent with our finding that wildtype (B6 allele) AAV9-*Rgs1* induced collateral formation in collateral-deficient CC016 mice and conclusion that deficient Rgs1 activity contributes to their reduced collaterogenesis/sparse collaterals. However, findings from other studies could be interpreted to suggest that increased signaling by certain GPCRs that may contribute to collaterogenesis, resulting from interacting with a deficient Rgs1, would be expected to lead to *more* abundant collaterals in CC016, and therefore that wildtype AAV9-*Rgs1* should have had no effect.^56–69: see Bioinformatics analysis in Supplement for details^ Future in vivo studies will be required to resolve this discrepancy, for example using conditional cell-specific gene editing in CC049 and CC016 mice to determine if and how, based on our findings, Rgs1 acts to augment collateral formation during development, while potentially restraining new collateral formation induced by arterial occlusion or exposure to prolonged hypoxemia reported in previous studies.^56–60^

This study has several limitations. Our in vivo assay did not identify a causal gene among the top candidates underlying *Canq6-10* (Table 1). It is possible the assay was not robust enough, for example if one or more of these proteins contributes to collaterogenesis, but—unlike Rgs1 and Rabep2 (the latter served as the assay’s positive control)—is not able to re-activate the process when given intravenously by an AAV9 expression vector on postnatal day-zero (the time point for which the assay was restricted). We were unable to develop a better-targeted method, eg, in utero delivery of antisense or transgenic sequences to the watershed regions during the last trimester when collaterogenesis is underway. Also, no in vitro model of collaterogenesis (eg, employing microfluidics) has been devised. Developing one or both of the above remain challenges for the future. As detailed in Results, we did not test certain genes in Table 1 for three reasons: because of their highly restricted expression, including not being expressed in vascular wall cells or other cells likely to be involved in collaterogenesis; because ORFs for three genes exceeded the packaging limit for AAV9; and because of the significant resources required to test the thirteen top-ranked genes. Several other limitations were encountered (detailed in Results), including that CC mice with targeted mutant genes are lacking, and that evaluating any candidate gene available on the B6 background is unlikely to be informative (we tried this approach for several genes without success—see Results). This limitation was expected, given our finding that collaterogenesis is a polygenic trait and thus that the effect size of any targeted allele is unlikely to be significant if not explicitly tested in a mouse strain where that arm of the collaterogenesis pathway is perturbed. Given the above limitations, all of the genes in Table 1 for *Canq6-10* remain candidates for future study.

We did not test the genes listed in Supplemental table II with predicted deleterious SNPs in the haploblock of the two high-collateral parental strains for two reasons: the considerable resources required to test the pro-collaterogenesis genes in Table 1; and the unlikelihood that collateral formation relies to a significant degree on “anti-”collaterogenesis genes providing a restraint on “pro”-collaterogenesis genes. This reasoning extends from evidence, detailed earlier in Discussion, that the presence of collaterals vessels in tissues is not required under normal physiological conditions, which in turn leads to the assumption that the process of collaterogenesis has not evolved counter-regulatory “balancing” mechanisms and/or compensatory redundancies that assure that collateral number and diameter do not vary significantly among individuals. This contrasts with the well-known complex interactions that exist between angiogenic and anti-angiogenic genes, pathways and compensatory mechanisms that are required to assure the anatomic constancy present in the general artery-capillary-vein circulations of tissues.^70^ Notwithstanding the aforementioned, we nevertheless identified with SNP analysis the genes listed in Supplemental table II to aid future investigations, since the above argument rests on theoretical grounds rather than experimental evidence. Likewise, the genes listed in Supplemental table III, which lack predicted damaging SNPs between the high and low-collateral parental strains but have functions potentially involved in collaterogenesis, were also identified for future studies because polymorphisms, if present in their regulatory regions, could cause differences in expression. Such studies will require obtaining one or more of the following from E14.5-E18.5 embryos:^19–21^ whole-mount mRNA and protein localization, and isolation of RNA and protein from the cell types within the pial watershed regions that are involved in collateral formation. Future studies might also investigate genes underlying *Canq5-10* harboring variant types in addition to coding and regulatory mutations (eg, functional splice region variants) for possible involvement in the collaterogenesis pathway.

In conclusion, we report using the genetically diverse Collaborative Cross mouse strain-set that collateral number and diameter evidence wide variation and are highly polygenic traits that result in large differences in infarct volume after MCA occlusion. This variation in pial collaterals extends to collaterals in other tissues of the same individual. The distribution of collateral number across the 60 CC strains partitioned into 14% with poor, 25% with poor-to-intermediate, 47% with intermediate-to-good, and 13% with good collaterals (Figure 1C). Pre-procedural scores for collateral blood flow in patients with acute large-vessel ischemic stroke show a similar distribution.^1–6^ This congruence suggests that genetic background-dependent differences in collateral abundance are a primary contributor to variation in stroke severity in humans. We also identify several novel genetic loci and a set of underlying genes including *Rgs1* as potential contributors, along with *Rabep2* identified previously,^22^ to variation in collaterals for future investigation. Such information is needed to better understand the signaling proteins within the collaterogenesis pathway and aid studies investigating the genetic basis for differences in collaterals among humans.^71–73^ It is also needed for translation going forward, given recent studies showing that new collaterals form in brain and heart following acute arterial occlusion as well as after several weeks of systemic hypoxemia.^35, 54, 63^ Identification of candidate “collateral genes” in mice will enable their investigation using single-gene SNP association analysis, with its significant statistical advantage over GWAS, against collateral score (and/or other ordinal or continuous-variable surrogate endophenotypes) in brain^1–6^ and collateral flow index in heart and lower extremities^10, 11^ in patients with vascular obstruction, chronic hypoxemia and hemoglobinopathies, and in healthy individuals adapted to reduced oxygen availability.^15^ Toward this end, we provide a comprehensive list of ancestry-associated single-nucleotide variants predicted to be damaging in human orthologs of the above-identified murine genes (Supplemental table IV). Testing the human orthologs of the candidates genes identified in our study using single-gene analysis provides a way forward, in lieu of the development of a robust collaterogenesis assay (s), toward identifying which of them are causal.

## Non-standard abbreviations and acronyms

ACA: anterior cerebral artery
AAV9: adenovirus-associated virus-9
B6: C57BL/6J mouse strain
*Canq*: <u>c</u>ollateral <u>a</u>rtery <u>n</u>umber <u>Q</u>TL (genetic locus associated with a difference in number of collaterals)
CC: Collaborative Cross mouse genetic reference panel of inbred strains
Chr: chromosome
EC: endothelial cell
GFP: (enhanced) green fluorescent protein
LOD: log of the odds ratio
LoF: Loss of function DNA variant/polymorphism
ORF: open reading frame
QTL: quantitative trait locus/loci
MCA: middle cerebral artery
SNP: single nucleotide polymorphism

## Acknowledgements

The authors thank Rachel Lynch and Darla Miller of the UNC SGCF for advice and assistance with animal husbandry, and Baldev Desai, Kathryn Conlon and Gabriel Gong for assistance with collateral diameter morphometry.

## Author contributions

JF: conceptualization, experimental design, project supervision, in silico SNP and gene analyses, methodological development, writing, funding acquisition; HZ: methodology development, data acquisition, statistical analysis, animal husbandry; JX: QTL mapping; TB and PH: genotyping; FPMV: advice and discussion; MF: strain selection for F2 generation, QTL mapping, advice and discussion, edits to the manuscript; WR: collateral diameter morphometry. All authors read and approved the manuscript.

## Supplemental material

Supplemental material for this paper is available online.

