## Supplemental Tables and figures for "Large differences in collateral blood vessel abundance among individuals arise from multiple genetic variants"

1. Supplemental figure I. *Canq5* and *Canq6* exhibit additive (independent) effects.
2. Supplemental figure II. Collateral diameter weakly correlates with collateral number in 60 Collaborative Cross strains.
3. Supplemental figure III. In vivo analysis of candidate genes for effect on collateral diameter.
4. Supplemental table I. Characteristics of QTL for variation in collateral number.
5. Supplemental table II. Genes underlying *Canq5*-to-*Canq10* with SNP(s) in the haplotype block of the high-collateral strains in the crosses shown in Table 1.
6. Supplemental table III. Genes within ± 10Mb of QTL peak that lack nonsense or missense SNPs but have functions known to be involved in or associated with collaterogenesis (a), angiogenesis (b), vascular development (c), endothelial cell function (d), or endocytic vesicle trafficking (e).
7. Supplemental table IV. Coding and nonsense variants present in human orthologs of mouse genes known to ("Known genes") and "Candidate genes" predicted to impair collaterogenesis.
8. Bioinformatics analysis of candidate genes' function, expression, and potential involvement in the collaterogenesis pathway.

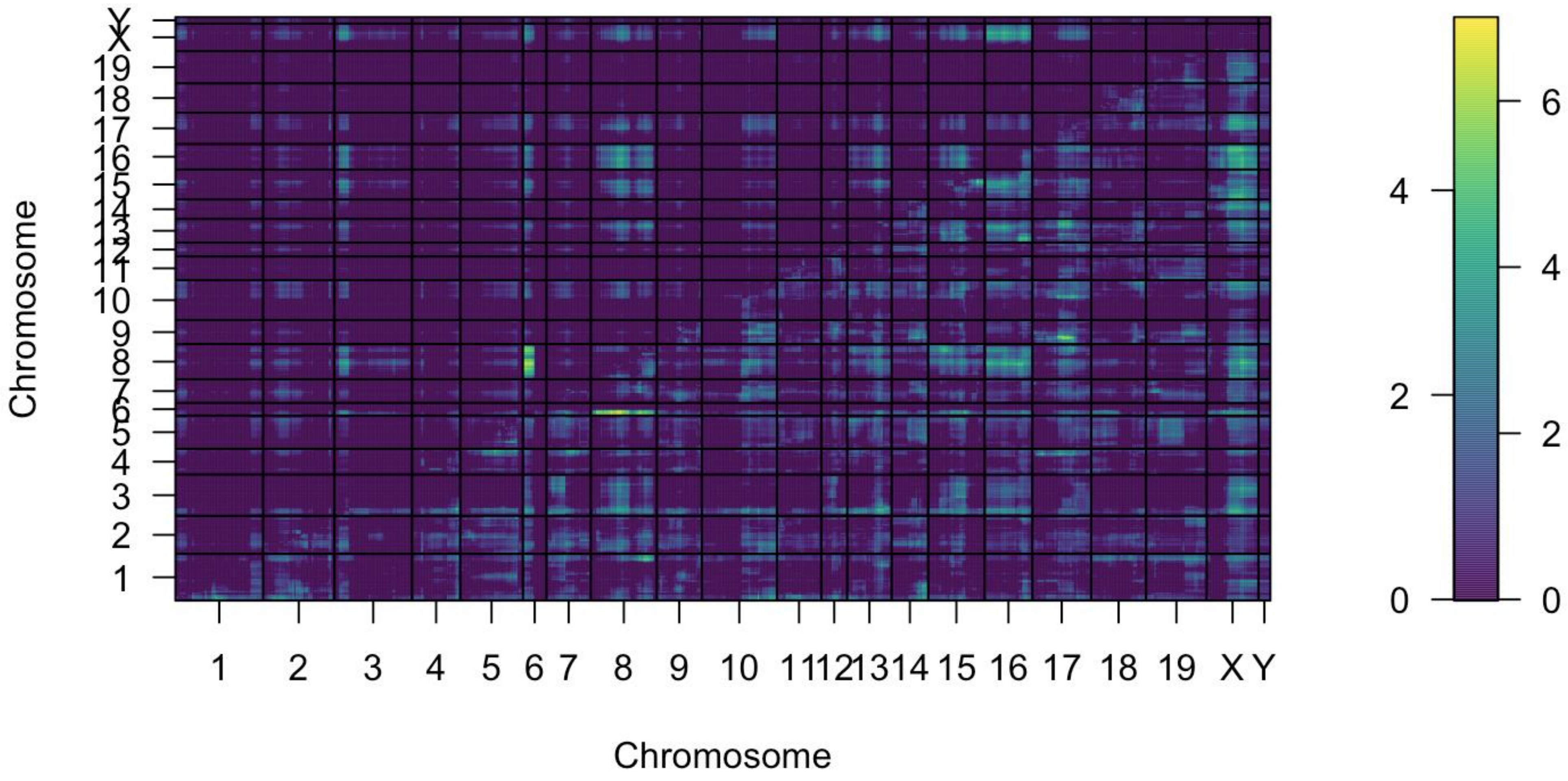


**Supplemental figure I.** Supplemental Figure 2: Two-QTL scan for CC036xCC055. To investigate possible pairwise joint effects of QTL, we performed a two-dimensional, two-QTL genome scan using the function scantwo() in R/qtl. In this plot, the marginal improvement in LOD score from the two QTL models with interactions versus null models with no QTL are visualized in the lower right triangle. The marginal improvements in LOD score from the full two-QTL models with interactions versus additive two-QTL models without interactions are visualized in the upper left triangle. This plot provides evidence for the additive effects of most loci in these models, and some evidence for an interactive effect between the QTL on chromosomes 6 and 8.

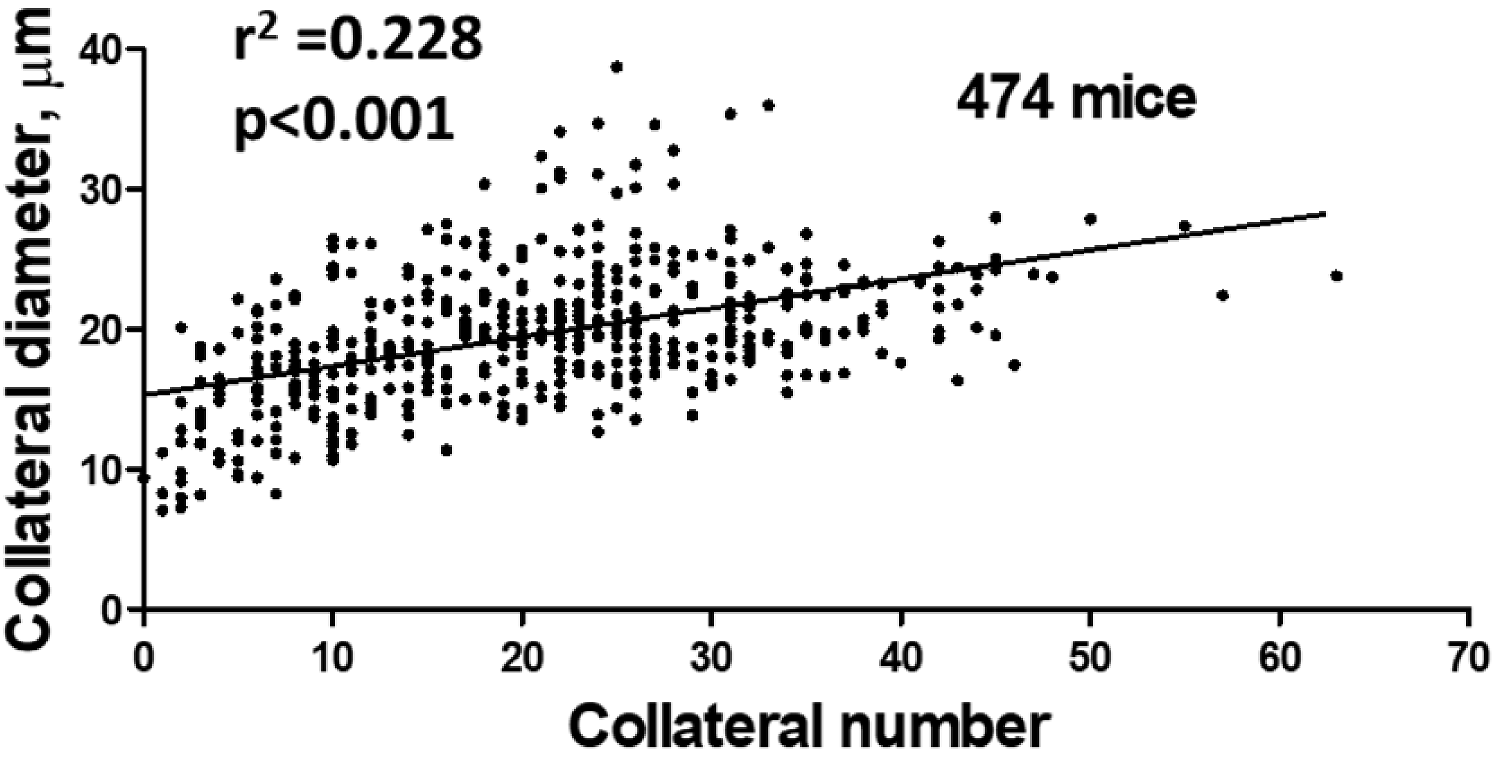


**Supplemental figure II. Collateral diameter weakly correlates with collateral number in 60 CC strains.** Data are taken from Figure 1. The low coefficient of determination for the 3.4-fold range in diameter and 47-fold range in number across the 60 CC strains shown in Figure 1 is consistent with previous findings (see Results and Discussion).

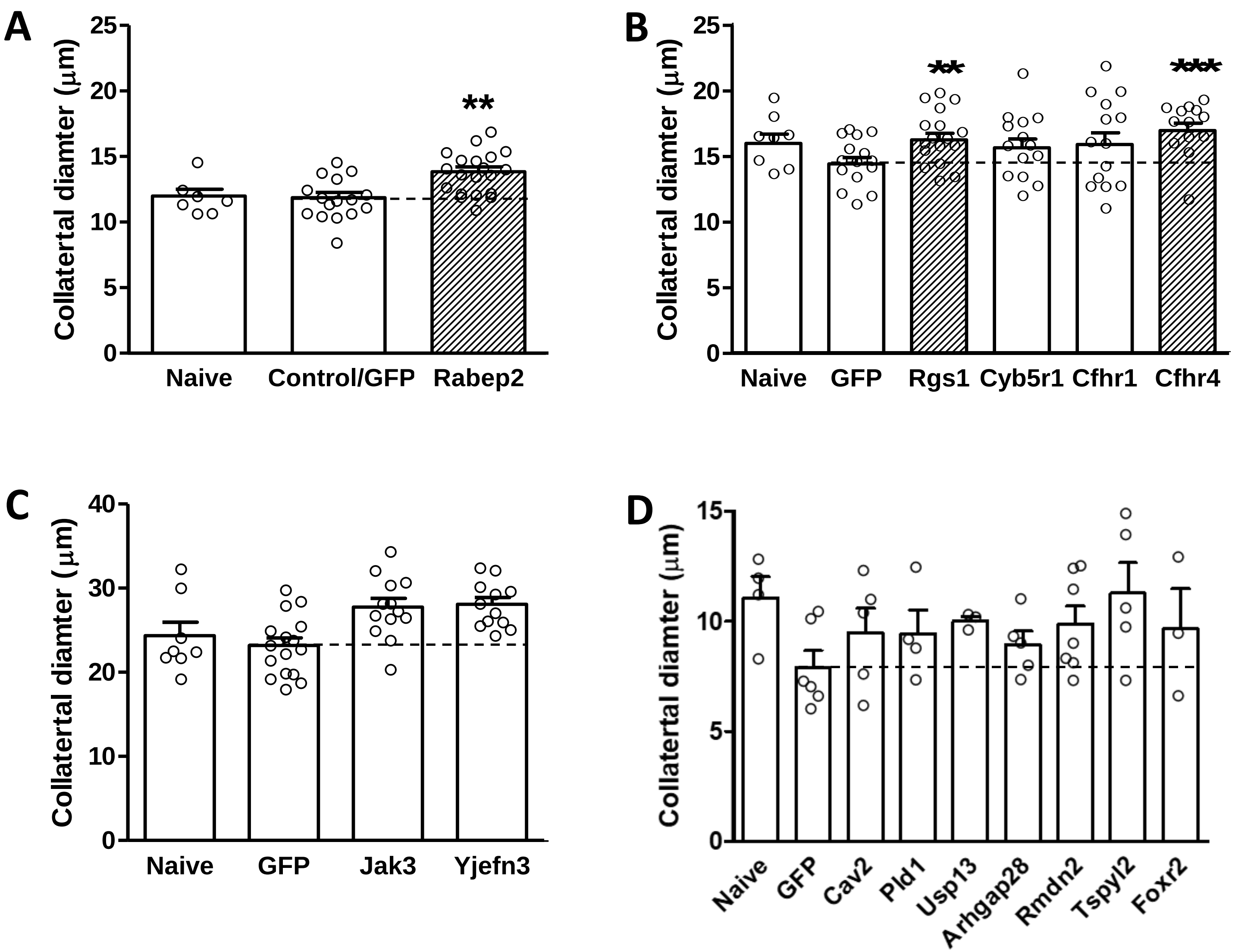


**Supplemental figure III. In vivo analysis of candidate genes for effect on collateral diameter.** AAV9-expression constructs for *Rabep2* (**panel A**) and selected high-priority candidate genes underlying *Canq5* (**B**), *Canq6* (**C**) and *Canq7-10* (**D**) were injected iv on postnatal day-zero before blood brain barrier closure into *Rabep2*$^{-/-}$ (A), CC016 (B), CC032 (C) and CC036 (D) mice. Average diameter for all collaterals between the MCA and ACA trees of both hemispheres was measured for each mouse at 6 weeks-age. Naïve, no injection; Control/GFP, injection of AAV9 construct for enhanced green fluorescent protein only. Number of animals for each bar: Panel A: 7,16,20; B: 8,15,17,14,14,13; C: 8,16,13,12; D: 4,6,5,4,3,5,7,5,3 (smaller n-sizes than in Figure 6D reflect animals with no collaterals). *p<0.05, ***p<0.001 vs. Control/GFP by 1-sided *t*-tests.

Supplemental table I. Characteristics of QTL for variation in collateral number

| Cross | QTL | Chr | Marker | Position | LOD, | P-value | Interval (M) | Haplotype blocks |
|---|---|---|---|---|---|---|---|---|
| 049 x 053 | *Canq5* | 1 | gUNC1733690 | 135,464,050 | 10.88 | <0.001 | 127.48-157.13 | WSB high, A/J low |
| 055 x 036 | *Canq6* | 8 | mJAX00671437 | 68,343,478 | 10.39 | <0.001 | 55.82-78.90 | 129S1 high, C57 low |
| 055 x 036 | *Canq7* | 6 | gJAX00603740 | 15,909,737 | 6.90 | <0.001 | 4.03-26.49 | CAST high, WSB low |
| 055 x 036 | *Canq8* | 3 | mUNC4993850 | 32,940,787 | 5.79 | <0.001 | 29.47-45.57 | CAST high, A/J low |
| 055 x 036 | *Canq9* | 17 | UNC28266829 | 70,113,081 | 5.84 | <0.001 | 60.00-80.94 | A/J high, WSB low |
| 055 x 036 | *Canq10* | X | gUNC31504684 | 157,934,381 | 5.85 | <0.001 | 141.50-164.42 | PWK high, 129S1ow |
| 055 x 036 | *Canq11* | 13 | mUNC130149286 | 85,808,377 | 4.78 | 0.008 | 3.56-109.60 | |
| 055 x 036 | *Canq12* | 16 | mUNC26808608 | 50,710,699 | 5.08 | 0.004 | 25.09-95.80 | |

Cross, parental CC strains; Chr, chromosome; Marker, Position, marker SNP and peak location on chromosome; 95% confidence interval, Haplotype blocks: “high” denotes haplotype block harboring locus for high collateral number (protective allele) in CC049 or CC055, “low” denotes haplotype block harboring locus for low collateral number (deleterious allele) in CC053 or CC036. P values are based on 1000 permutations.

Supplemental table II. Genes underlying *Canq5*-to-*Canq10* with SNP(s) in the haplotype block of the high-collateral strains in the crosses shown in Figure 4A.

| Locus | Symbol | Description | RS number | Coding change | Allele | P-value for deleterious effect | | |
|---|---|---|---|---|---|---|---|---|
| | | | | | | SIFT | SIFT-P | PROV* |
| *Canq5* | | | | | | | | |
| | *Igfn1* | immunoglobulin-like and fibronectin type III domain containing 1 | rs30911401 | P607S | W | 0.020 | 0.012 | -4.43 |
| | | | rs253370915[a] | S633P | W | 0.030 | 0.001 | -1.89 |
| | *Crb1* | crumbs family member 1 | rs30924175 | P269L | W | 0.000 | 0.002 | -8.50 |
| | *Cacna1s* | calcium channel, voltage-dependent, L type, alpha 1S subunit | rs223895038 | P3L | W | 0.000 | 0.000 | -1.30 |
| | *Il19* | interleukin 19 | rs33497741 | S105I | W | 0.000 | 0.003 | -3.82 |
| | | | rs33495994 | T8P | W | 0.020 | 0.014 | -0.82 |
| | *Kif14* | kinesin family member 14 | rs238546255 | E426G | W | 0.010 | 0.090 | -3.31 |
| | *Etnk2* | ethanolamine kinase 2 | 133,365,816[b] | Stop | W | | | |
| | *Ppp1r15b* | protein phosphatase 1, regulatory subunit 15B | rs33493063 | P398S | W | 0.020 | 0.002 | -2.84 |
| | *Cntn2* | contactin 2 | rs36402706 | R64H | W | 0.000 | 0.002 | -2.54 |
| | *Lax1* | lymphocyte transmembrane adaptor 1 | rs32582773 | P67S | W | 0.080 | 0.048 | -6.72 |
| | *Pigr* | polymeric immunoglobulin receptor | rs241665934 | P607S | W | 0.010 | 0.030 | -2.86 |
| | | | rs30820548 | T39M | W | 0.060 | 0.030 | -1.08 |
| | *Thsd7b* | thrombospondin, type I, domain containing 7B | rs264931691 | R1154Q | W | 0.020 | 0.001 | -2.73 |
| | *Acmsd* | amino carboxymuconate semialdehyde decarboxylase | rs46700071 | T69I | W | 0.020 | 0.012 | -3.65 |
| | *Zranb3* | zinc finger, RAN-binding domain containing 3 | rs222620320 | R317C | W | 0.010 | 0.027 | -4.00 |
| | *Lct* | lactase | rs50723357 | T939M | W | 0.000 | 0.001 | -2.40 |
| | *Map3k19* | mitogen-activated protein kinase kinase kinase 19 | rs46700071 | T69I | W | 0.020 | 0.012 | -3.65 |
| *Canq6* | | | | | | | | |
| | *Nek1* | NIMA (never in mitosis gene a)-related expressed kinase 1 | rs239230033 | R39S | 1 | 0.050 | 0.050 | -2.71 |
| | *Sh3rf* | SH3 domain containing ring finger 1 | rs33135970 | G84V | 1 | 0.020 | 0.004 | -3.84 |
| | *Gdf1* | growth differentiation factor 1 | rs32650885 | S145C | 1 | 0.020 | 0.014 | -2.16 |
| | *Prmt9* | protein arginine methyltransferase 9 | rs45636255 | Q824H | P | 0.050 | 0.066 | -1.59 |
| | *Nwd1* | NACHT and WD repeat domain containing 1 | rs237177178 | Y1386C | P | 0.010 | 0.020 | -2.56 |
| | *Sugp2* | SURP and G patch domain containing 2 | rs32650665[c] | Stop | 1 | | | |
| | *Zfp963* | zinc finger protein 963 | rs33361474 | S185N | 1 | 0.050 | 0.038 | -1.20 |
| | | | rs32816029 | E49K | 1 | 0.030 | 0.027 | -2.06 |
| *Canq7* | | | | | | | | |
| | *Tfec* | transcription factor EC (E-box containing) | rs37583912 | R306Q | C | 0.020 | 0.029 | -2.75 |
| | | | rs241258046 | D9E | C | 0.010 | 0.008 | -0.01 |
| | *Ppp1r3a* | protein phosphatase 1, regulatory subunit 3A | rs33469811 | D870H | C | 0.000 | 0.001 | -2.02 |
| | *Ptprz1* | protein tyrosine phosphatase, receptor type Z, polypeptide 1 | rs240106830 | L8F | C | 0.000 | 0.002 | -0.83 |
| | | | rs33747302 | L843P | C | 0.030 | 0.040 | -1.35 |
| | | | rs33748034 | T893A | C | 0.010 | 0.070 | -1.20 |
| | | | rs33748042 | A1181T | C | 0.000 | 0.001 | -0.05 |
| | *Vwde* | Willebrand factor D and EGF domains | rs46632271 | L9M | C | 0.020 | 0.037 | -0.34 |
| | *Tmem168* | transmembrane protein 168 | rs212279097 | N332D | C | 0.070 | 0.042 | -0.45 |
| | *Pot1a* | protection of telomeres 1A | rs226408196 | V551I | C | 0.030 | 0.033 | -0.39 |
| | *Cttnbp2* | cortactin binding protein 2 | rs36251448 | P1213T | C | 0.090 | 0.087 | -1.52 |
| *Canq8* | | | | | | | | |
| | *Plk4* | polo like kinase 4 | rs49839395 | S601A | C | 0.040 | 0.028 | -2.37 |
| | *Skil* | SKI-like | rs29841544 | P367A | C | 0.040 | 0.030 | -1.32 |
| | | | rs29814968 | L463F | C | 0.060 | 0.021 | -1.15 |
| | *Intu* | inturned planar cell polarity protein | rs255853181 | R273W | C | 0.010 | 0.011 | -0.80 |
| | *Actrt3* | actin related protein T3 | rs246289103 | V149M | C | 0.000 | 0.000 | -2.56 |
| | *Samd7* | sterile alpha motif domain containing 7 | rs252671701 | P45L | C | 0.020 | 0.068 | -0.86 |
| | | | rs261109659 | P140S | C | 0.080 | 0.008 | -4.47 |
| | *Lrriq4* | leucine-rich repeats and IQ motif containing 4 | rs253521416 | I31N | C | 0.010 | 0.003 | -0.68 |
| | *1810062G17Rik* | RIKEN cDNA 1810062G17 gene | rs259087812 | Stop[d] | C | | | |
| | *Mfn1* | mitofusin 1 | rs29864746 | A641V | C | 0.070 | 0.016 | -1.91 |
| | *Ccdc144b* | coiled-coil domain containing 144B | rs49535369 | V68E | C | 0.020 | 0.382 | -1.52 |
| | *Ccna2* | cyclin A2 | rs229616401 | A144D | C | 0.030 | 0.011 | 0.78 |
| *Canq9*[e] | | | | | | | | |
| | *NdufaF7* | NADH:ubiquinone oxidoreductase complex assembly factor 7 | rs50621970 | R336H | A | 0.000 | 0.031 | -3.59 |
| *Canq10* | | | | | | | | |
| | *Gja6* | gap junction protein, alpha 6 | rs29305426 | I283T | P | 0.020 | 0.013 | 0.89 |
| | *Wnk3* | WNK lysine deficient protein kinase 3 | rs29293581 | R725Q | P | 0.000 | 0.000 | -3.71 |
| | *Scml2* | Scm polycomb group protein like 2 | rs248836551 | P628A | P | 0.030 | 0.006 | -2.20 |
| | *Ofd1* | OFD1, centriole and centriolar satellite protein | rs31559944 | F1010C | P | 0.050 | 0.006 | -1.91 |
| | *Zrsr2* | zinc finger (CCCH type) | rs252840144 | S424C | P | 0.040 | 0.030 | -2.45 |
| | *Magea5*[f] | MAGE family member A5 | rs583713186 | K185N | P | 0.040 | 0.014 | -4.63 |
| | *Asb11* | ankyrin repeat and SOCS box-containing 11 | rs31454203 | Q3P | P | 0.020 | 0.009 | -0.21 |

| | | | | | | | |
|---|---|---|---|---|---|---|---|
| *Frmpd4* | FERM and PDZ domain containing 4 | rs254540233 | M42I | P | 0.050 | 0.023 | -0.57 |
| *Cdkl5* | cyclin-dependent kinase-like 5 | rs29305426 | I283T | P | 0.020 | 0.013 | 0.89 |
| *Adgrg2* | adhesion G protein-coupled receptor G2 | rs257666586 | I670T | P | 0.030 | 0.16 | 1.73 |
| *Tro* | Trophinin | rs29297514 | A1267G | P | 0.000 | ----- | -0.93 |

Table lists the genes in the high-collateral strain, their SNP rs numbers, coding change, allele and P-values (See legend to Table 1 for additional information). A,W,C,B,1,P, abbreviations for strain-specific haplotype allele harboring SNP for, respectively: A/J, WSB/EiJ, CAST/EiJ, C57BL/6J, 129S1/SvlmJ, PWK/PhJ. [a], Five additional SNPs (W allele) are < 0.1 by SIFT, 3/5 < 0.05 by SIFT-P, 5/5 -0.21 to -2.24 by PROVEAN. [b], Stop is in first of 8 exons. [c], Stop at amino acid position 58 of 158. [d], Stop at amino acid position 10 of 129. [e], interval analysis was confined to -5 → +10 Mb due to haplotype heterozygosity. [f], *Magea5* has a private missense snp (C/T) at location 155089004 that is predicted deleterious ($p < 0.05$).

Supplemental table III. Genes within ± 10Mb of QTL peak that lack nonsense or missense SNPs but have functions known or potentially involved in collaterogenesis (a), angiogenesis (b), vascular development (c), endothelial cell function (d), or endocytic vesicle trafficking (e).

| QTL | Symbol | Description |
|---|---|---|
| *Canq5* | *Cxcr4*[a,b,c,f] | chemokine (C-X-C motif) receptor 4 |
| | *Atp2b4*[b,d] | ATPase, Ca++ transporting, plasma membrane 4 |
| | *Optc*[b,d] | opticin |
| | *Cfh*[b,d] | complement component factor h |
| | *Pik3c2b*[e] | phosphatidylinositol-4-phosphate 3-kinase catalytic subunit type 2 beta |
| | *Dennd1b*[e] | DENN/MADD domain containing 1B |
| *Canq6* | *C1galt1*[a,b,c,d] | core 1 synthase, glycoprotein-N-acetylgalactosamine 3-beta-galactosyltransferase, 1 |
| | *Hmox1*[b,c,d] | heme oxygenase 1 |
| | *Apela*[b,c,d] | apelin receptor early endogenous ligand |
| | *Lpar2*[b,c,d,e] | lysophosphatidic acid receptor 2 |
| | *Fkbp8*[b,d] | FK506 binding protein 8 |
| | *Rab3a*[b,d,e] | RAB3A, member RAS oncogene family |
| | *Plvap*[b,d] | plasmalemma vesicle associated protein |
| | *Fcho1*[e] | FCH domain only 1 |
| | *Rab8a*[e] | RAB8A, member RAS oncogene family |
| | *Ap1m1*[e] | adaptor-related protein complex AP-1, mu subunit 1 |
| | *Klf2*[b,c,d] | kruppel-like factor 2 |
| *Canq7* | *Cav1*[b,c,d,e] | caveolin 1 |
| | *Wnt2*[b,c,d] | wingless-type MMTV integration site family, member 2 |
| *Canq8* | *Pik3ca*[b,d] | phosphatidylinositol-4,5-bisphosphate 3-kinase catalytic subunit alpha |
| *Canq9* | *Rab31*[e] | RAB31, member RAS oncogene family |
| | *Twsg1*[c,d] | twisted gastrulation BMP signaling modulator 1 |
| | *Washc1*[e] | WASH complex subunit 1 |
| | *Rab12*[e] | RAB12, member RAS oncogene family |
| | *Ptprm*[d] | protein tyrosine phosphatase, receptor type, M |
| | *Lama1*[c,d] | laminin, alpha 1 |
| | *Ehd3*[e] | EH-domain containing 3 |
| | *Ltbp1*[c,d] | latent transforming growth factor beta binding protein 1 |
| | *Crim1*[c,d] | cysteine rich transmembrane BMP regulator 1 |
| | *Cyp1b1*[b,c,d] | cytochrome P450, family 1, subfamily b, polypeptide 1 |
| *Canq10* | *Vegfd*[b,c,d] | vascular endothelial growth factor D |
| | *Rab9*[d,e] | RAB9, member RAS oncogene family |

Genes are listed under each locus in order of position (-10 -to- +10Mb, respectively). [e], The association with endocytic vesicles arises as a criteria for inclusion in the table based on the known role of the endocytic protein, *Rabep2*, in collaterogenesis in certain classical strains of mice.[22,25,26 in manuscript] [f], See In Silico Analysis in Supplement for references linking Cxcr4 to collaterogenesis.

Supplemental table IV. Coding and nonsense variants present in human orthologs of mouse genes known to ("Known genes") and "Candidate genes" predicted to impair collaterogenesis. Single nucleotide variants (SNVs) obtained from the Genome Aggregation Database (gnomAD v3.1.2) are listed with their physical position on the indicated chromosome (Chr), rs number, amino acid substitution, and overall frequency of occurrence (Freq; see gnomAD for ancestry-specific frequencies). SNVs are listed if present in at least 10 individuals (Counts) out of the total Number of individuals genotyped and if predicted to be "probably damaging" by at least one of—and "possibly damaging" by at least one of: Polyphen2 (PP), SIFT or MetaLR (Meta) *in silico* algorithms per Ensemble (v109, 2/2023). Values calculated by Mutation Assessor (MutA), Revel and CADD prediction algorithms are also listed. If Polyphen2 values were not available, values for another algorithm were substituted for PP's criterion (eg, see *RGS1*). Red font, probably damaging; orange, possibly damaging; green, tolerated; blue, uncertain. AA, African/African American; AJ, Ashkenazi Jewish; AM, Amish; EA, East Asian; FN, Finnish; LT, Latino/Admixed American; ME, Middle Eastern; SA, South Asian. Positions of residues of stops gained are given. Superscripted numbers attached to each "Known Gene" refer to the references given below the table that identified the gene using QTL analysis and CRISPR (*Rabep2*)[1] or reverse-genetic approaches (gene targeting, etc) for other genes.[2-8] "Candidate Genes" are from Table 1.

| Position | rs number | Coding SNV | Count | Number | Freq | PP | SIFT | Meta | MutA | Revel | CADD | Ancestry counts |
|---|---|---|---|---|---|---|---|---|---|---|---|---|
| **Known Genes** | | | | | | | | | | | | |
| ***RABEP2***[1] Chr 16 | | | | | | | | | | | | |
| 28905025 | rs527458355 | p.Arg543His | 24 | 152166 | 1.58e-4 | 0.994 | 0.00 | 0.673 | 0.763 | 0.647 | 28 | 23 AA, 1 SA |
| 28905481 | rs200118396 | p.Arg508Ser | 94 | 152154 | 6.18e-4 | 0.986 | 0.00 | 0.230 | 0.700 | 0.267 | 25 | 57 EU, 18 LT, 7 SA |
| 28905727 | rs184144701 | p.Arg490Trp | 12 | 152098 | 7.89e-5 | 0.997 | 0.00 | 0.216 | 0.781 | 0.367 | 29 | 8 EA, 3 AA, 1 AM |
| 28906121 | rs750752217 | p.Arg441Cys | 12 | 152182 | 7.89e-5 | 0.997 | 0.00 | 0.255 | 0.685 | 0.412 | 25 | 8 EU, 2 AA, 2 AJ |
| 28911143 | rs373485868 | p.Arg311Cys | 13 | 152178 | 8.54e-5 | 0.769 | 0.00 | 0.128 | 0.338 | 0.209 | 24 | 12 AA, 1 LT |
| 28914373 | rs200278634 | p.Arg253Cys | 14 | 152192 | 9.20e-5 | 0.825 | 0.03 | 0.164 | 0.314 | 0.138 | 24 | 11 AA, 3 EU |
| 28914519 | rs769480150 | p.Ser204Leu | 49 | 152196 | 3.22e-4 | 0.978 | 0.01 | 0.209 | 0.280 | 0.099 | 25 | 44 FN, 5 EU |
| ***VEGFA*** [2,3] Chr 6 | | | | | | | | | | | | |
| 43770809 | rs1055085365 | p.Glu35Ter | 16 | 152154 | 1.05e-4 | Stop gained 35/395 | | | | | | 14 AA, 1, EU, 1 other |
| 43770920 | rs1004481569 | p.Gly72Trp | 11 | 152180 | 7.23e-5 | 0.984 | 0.000 | 0.061 | ----- | 0.125 | 25 | 11 EU |
| ***KDR***[3] Chr 4 | | | | | | | | | | | | |
| 55080075 | rs151242590 | p.Asp1313Asn | 10 | 152224 | 6.57e-5 | 0.984 | 0.000 | 0.556 | ----- | 0.000 | 24 | 5 AA, 4 EU, 1 other |
| 55088923 | rs373562441 | p.Thr1152Met | 33 | 152168 | 2.17e-4 | 0.898 | 0.010 | 0.816 | 0.697 | 0.000 | 28 | 32 AA, 1 LT |
| 55088939 | rs121917766 | p.Pro1147Ser | 12 | 152164 | 7.89e-5 | 0.998 | 0.010 | 0.847 | 0.536 | 0.000 | 26 | 8 EU, 3 AA, 1 LT |
| 55089802 | rs56302315 | p.Ala1065Thr | 62 | 152106 | 4.08e-4 | 0.955 | 0.010 | 0.663 | 0.089 | 0.000 | 32 | 42 AJ, 19 EU, 1 AA |
| 55094936 | rs140041720 | p.Arg946His | 37 | 152144 | 2.43e-4 | 0.980 | 0.020 | 0.464 | 0.153 | 0.000 | 28 | 17 EA, 8 AA, 7 LT, 5 EU |
| 55097760 | rs140761530 | p.Pro839Leu | 12 | 151660 | 7.91e-5 | 1.000 | 0.020 | 0.993 | 0.038 | 0.000 | 24 | 6 AA, 6 EA |
| 55098158 | rs145905001 | p.Pro830Ser | 11 | 152102 | 7.23e-5 | 1.000 | 0.010 | 0.792 | 0.560 | 0.000 | 26 | 6 EU, 4 LT, 1 AA |
| 55098263 | rs564385300 | p.Gly795Arg | 13 | 152160 | 8.54e-5 | 1.000 | 0.998 | 0.523 | 0.570 | 0.000 | 26 | 8 SA, 4 ME, 1 other |
| 55114137 | rs587778429 | p.Pro263Thr | 10 | 152068 | 6.58e-5 | 0.936 | 0.000 | 0.270 | 0.942 | 0.000 | 24 | 9 FN, 1 EU |
| ***ADAM17***[3] Chr 2 | | | | | | | | | | | | |
| No predicted deleterious SNVs | | | | | | | | | | | | |
| ***DLL4***[4] Chr 15 | | | | | | | | | | | | |
| 40931580 | rs145914797 | p.Asp158Asn | 19 | 152220 | 1.25e-4 | 0.326 | 0.030 | 0.871 | 0.769 | 0.000 | 26 | 19 AA |
| 40936273 | rs370339149 | p.Arg429His | 12 | 152220 | 7.88e-5 | 0.723 | 0.020 | 0.196 | 0.418 | 0.000 | 27 | 9 AA, 3 EU |
| 40936539 | rs201981567 | p.Glu518Lys | 33 | 152244 | 2.17e-4 | 0.986 | 0.010 | 0.826 | 0.912 | 0.000 | 28 | 20 FN, 9 EU, 2 AJ, 1 AA |
| 40936645 | rs375688749 | p.Arg553Gln | 13 | 152218 | 8.54e-5 | 0.992 | 0.060 | 0.775 | 0.615 | 0.000 | 26 | 9 EU, 2 FN, 1 AA, 1 SA |
| ***NOTCH1***[3,4] Chr 9 | | | | | | | | | | | | |
| 136502059 | rs201779159 | p.Leu1805Pro | 15 | 152080 | 9.86e-5 | 0.937 | 0.010 | 0.676 | 0.869 | 0.648 | 27 | 12 EU, 1 AA, 1 LT, 1 SA |
| 136502383 | rs373841359 | p.Arg1758His | 16 | 152240 | 1.05e-4 | 0.937 | 0.000 | 0.651 | 0.886 | 0.546 | 27 | 9 LT, 5 EU, 1 AA |
| 136504793 | rs375018022 | p.Arg1633His | 88 | 152208 | 5.78e-4 | 0.860 | 0.010 | 0.517 | 0.454 | 0.425 | 24 | 81 AA, 5 LT, 1 EU, 1 SA |
| 136504896 | rs543770603 | p.Val1599Met | 11 | 152200 | 7.23e-5 | 0.706 | 0.000 | 0.679 | 0.446 | 0.430 | 23 | 5 LT, 3 EU, 3 AA |
| 136505359 | rs765844768 | p.Gly1513Ser | 15 | 152216 | 9.85e-5 | 0.343 | 0.010 | 0.659 | 0.605 | 0.503 | 24 | 11 AA, 2, EU, 2 LT |
| 136506571 | rs149057410 | p.Val1324Met | 67 | 152154 | 4.40e-4 | 0.992 | 0.020 | 0.704 | 0.161 | 0.543 | 23 | 61 AA, 2 LT, 1 EU, 1 SA |
| 136509030 | rs201163739 | p.Ser1004Leu | 24 | 152260 | 1.58e-4 | Stop gained 1004/2555 | | | | | 19 | EU, 3 AJ, 2 LT |
| 136509839 | rs370797169 | p.Arg955Cys | 15 | 152238 | 9.85e-5 | 0.891 | 0.000 | 0.690 | 0.813 | 0.474 | 28 | 11 EU, 2 EA, 1 AA, 1 ME |
| 136510659 | rs201620358 | p.Arg912Trp | 333 | 152244 | 2.19e-3 | 0.846 | 0.000 | 0.573 | 0.239 | 0.402 | 25 | 235EU, 45LT, 25AA, 17 FN |
| 136510719 | rs578228721 | p.Arg892Cys | 46 | 152244 | 3.02e-4 | 0.952 | 0.000 | 0.778 | 0.621 | 0.715 | 24 | 40 LT, 4 AA, 1 EU, 1 other |

| | | | | | | | | | | | | |
|---|---|---|---|---|---|---|---|---|---|---|---|---|
| 136511244 | rs559917218 | p.Pro832Leu | 56 | 152138 | 3.68e-4 | 0.999 | 0.010 | 0.757 | **0.603** | 0.680 | 28 | 47 LT, 5 AA, 3 EU, 1 other |
| 136513054 | rs201620755 | p.Gly812Arg | 39 | 151732 | 2.57e-4 | 1.000 | 0.030 | 0.743 | 0.443 | 0.724 | 25 | 36 AA, 3 EU |
| 136513129 | rs199672693 | p.Asn787Asp | 16 | 152062 | 1.05e-4 | 0.999 | 0.010 | 0.802 | **0.578** | 0.616 | 27 | 13 LT, 1 AA, 2 other |
| 136514511 | rs201662530 | p.Gly736Arg | 20 | 152234 | 1.31e-4 | 0.189 | 0.010 | 0.626 | **0.657** | 0.696 | 13 | 12 AA, 7 EU, 1 LT |
| 136517898 | rs200562991 | p.Thr432Met | 19 | 152072 | 1.25e-4 | 1.000 | 0.000 | 0.891 | 0.503 | 0.835 | 26 | 6 EU, 5 AJ, 4 SA, 3LT, 1 EA |
| 136519457 | rs376104770 | p.Pro284Leu | 19 | 152244 | 1.25e-4 | 0.910 | 0.180 | 0.877 | 0.448 | 0.743 | 23 | 14 EU, 5 LT |
| 136519469 | rs367825691 | p.Asn280Ser | 24 | 152172 | 1.58e-4 | 0.999 | 0.040 | 0.601 | 0.037 | 0.517 | 24 | 20 EU, 3 LT, 1 AA |
| 136522891 | rs150737112 | p.Arg234His | 63 | 152172 | 4.14e-4 | 0.952 | 0.600 | 0.515 | 0.100 | 0.462 | 23 | 32 EU, 24 LT, 6 AA, 1 SA |
| 136522918 | rs553542677 | p.Ser225Trp | 10 | 152228 | 6.57e-5 | 0.999 | 0.010 | 0.744 | 0.461 | 0.736 | 29 | 10 AA |
| 136523138 | rs750242131 | p.Gly152Ser | 22 | 152222 | 1.45e-4 | **0.742** | 0.020 | 0.909 | **0.809** | 0.788 | 24 | 14 AA, 8 EU |

***ADAM10***[3] Chr 15

No predicted deleterious SNVs

***EPHA4***[5] Chr 2

| | | | | | | | | | | | | |
|---|---|---|---|---|---|---|---|---|---|---|---|---|
| 221426033 | rs141387215 | p.Val986Ile | 93 | 151988 | 6.12e-4 | 0.994 | 0.000 | 0.514 | 0.692 | 0.328 | 26 | 87 AA, 5, LT, 1 EU |
| 221436595 | rs370645355 | p.Arg717Ile | 17 | 152210 | 1.12e-4 | 0.942 | 0.000 | 0.526 | 0.501 | 0.577 | 26 | 15 EU, 1 AA, 1 LT |

***CLIC4***[6,7] Chr 1

No predicted deleterious SNVs

***GJA4***[8] Chr 1

| | | | | | | | | | | | | |
|---|---|---|---|---|---|---|---|---|---|---|---|---|
| 34794466 | rs373204338 | p.Val85Ile | 18 | 152108 | 1.18e-4 | 0.992 | 0.000 | 0.973 | **0.543** | 0.784 | 27 | 13 SA, 5 EU |
| 34794745 | rs138853660 | p.Gly178Ser | 12 | 152164 | 7.89e-5 | 0.999 | 0.000 | 0.957 | **0.833** | 0.952 | 28 | 12 EU |
| 34794836 | rs147387551 | p.Ile208Thr | 42 | 152092 | 2.76e-4 | **0.462** | 0.000 | 0.917 | 0.948 | 0.934 | 27 | 36 EU, 6 AA |
| 34794925 | rs201475159 | p.Arg238Trp | 10 | 152148 | 6.57e-5 | 0.167 | 0.010 | 0.847 | 0.314 | 0.313 | 21 | 7 AA, 3 EA |
| 34795178 | rs151235389 | p.Arg322His | 10 | 151836 | 6.59e-5 | **0.837** | 0.010 | 0.852 | **0.560** | 0.339 | 23 | 5 AA, 2 FN, 2 LT, 1 EU |

### Candidate Genes

***SOX13*** Chr 1

| | | | | | | | | | | | | |
|---|---|---|---|---|---|---|---|---|---|---|---|---|
| 204116516 | rs1284301034 | p.Glu143Gly | 10 | 152222 | 6.57e-5 | 0.994 | 0.00 | 0.965 | **0.773** | 0.847 | 30 | 10 LT |
| 204122265 | rs756405105 | p.Ala297Asp | 11 | 152150 | 7.23e-5 | 0.976 | 0.01 | 0.967 | **0.866** | 0.696 | **25** | 10 F, 1 EU |
| 204122282 | rs199517579 | p.Glu303Lys | 41 | 151944 | 2.70e-4 | **0.665** | 0.01 | 0.922 | **0.853** | 0.482 | 26 | 28 Lat, 11 EU, 2 AA |
| 204122959 | rs201772428 | p.Arg377His | 150 | 152180 | 9.86e-4 | 0.996 | 0.00 | 0.944 | **0.832** | 0.495 | 24 | 111 EU, 12 AA, 10 F, 10 SA |
| 204123205 | rs561040843 | p.Asp410Tyr | 12 | 152242 | 7.88e-5 | **0.678** | 0.00 | 0.930 | **0.835** | 0.760 | 33 | 11 SA, 1 EA |
| 204123685 | rs771489813 | p.Arg419Gln | 23 | 152140 | 1.51e-4 | 0.992 | 0.01 | 0.954 | **0.875** | 0.643 | 32 | 23 AA |

***CYB5R1*** Chr 1

| | | | | | | | | | | | | |
|---|---|---|---|---|---|---|---|---|---|---|---|---|
| 202963091 | rs368686887 | p.Trp240Cys | 17 | 152154 | 1.12e-4 | 1.000 | 0.00 | 0.849 | 0.959 | 0.860 | 31 | 16 EA, 1 SA **[stop gained 240/305]** |
| 202964651 | rs2232845 | p.Arg174Ter | 12 | 152174 | 7.89e-5 | Stop gained 174/305 | | | | | | 10 AA, 2 F |
| 202966750 | rs199636712 | p.Thr55Met | 13 | 152178 | 8.54e-5 | **0.877** | 0.00 | 0.762 | **0.899** | 0.502 | 33 | 13 LT |

***RGS1*** Chr 1

| | | | | | | | | | | | | |
|---|---|---|---|---|---|---|---|---|---|---|---|---|
| 192579159 | rs150745219 | p.Arg156Gln | 31 | 151926 | 2.04e-4 | ------ | 0.000 | 0.288 | 0.821 | 0.391 | 31 | 20 EU, 8 AA |

***ASPM*** Chr 1

| | | | | | | | | | | | | |
|---|---|---|---|---|---|---|---|---|---|---|---|---|
| 197089983 | rs542967760 | p.Pro3311Ala | 11 | 151616 | 7.26e-5 | 0.001 | **0.650** | 0.321 | **0.731** | 0.459 | 27 | 11 LT |
| 197090384 | rs368682161 | p.Leu3214Ser | 13 | 152022 | 8.55e-5 | 0.999 | 0.020 | 0.164 | **0.795** | 0.371 | 28 | 9 EU, 2 LT, 1 AA |
| 197090909 | rs140679756 | p.Arg3193Cys | 14 | 151970 | 9.21e-5 | 1.000 | 0.030 | 0.714 | **0.848** | 0.585 | 26 | 7 AA, 7 EU |
| 197090947 | rs193251130 | p.Gln3180Pro | 199 | 152122 | 1.31e-3 | 0.890 | 0.010 | 0.287 | **0.735** | 0.282 | 21 | 184 SA, 5 AA, 5 EU, 3 LT |
| 197091912 | rs150906798 | p.Ile3147Val | 12 | 152018 | 7.89e-5 | 0.968 | 0.050 | 0.437 | 0.474 | 0.097 | 23 | 11 EU, 1 LT |
| 197091956 | rs36004306 | p.Leu3132Pro | 14 | 152004 | 9.21e-5 | 0.924 | 0.000 | 0.355 | 0.338 | 0.336 | 22 | 12 EU, 1 SA, 1 LT |
| 197093059 | rs146444278 | p.Arg3096Gln | 13 | 151868 | 8.56e-5 | 0.998 | 0.010 | 0.446 | **0.739** | 0.477 | 25 | 7 EU, 6 AA |
| 197093074 | rs147005963 | p.Arg3091His | 85 | 151900 | 5.60e-4 | 0.997 | 0.000 | 0.560 | **0.739** | 0.668 | 23 | 81 AA, 2 EU, 2 LT |
| 197094116 | rs753961004 | p.Lys3018Glu | 10 | 151856 | 6.59e-5 | **0.643** | 0.010 | 0.684 | 0.322 | 0.415 | 25 | 10 AA |
| 197096010 | rs149690383 | p.Leu2992Pro | 39 | 151868 | 2.57e-4 | 0.950 | 0.110 | 0.753 | 0.158 | 0.419 | 11 | 33 EU, 4 AA, 1 FN, 1 LT |
| 197096099 | rs533985055 | p.Gln2962His | 15 | 151850 | 9.88e-5 | 1.000 | 0.000 | 0.846 | 0.964 | 0.764 | 24 | 15 AA |
| 197100727 | rs112946633 | p.Arg2842Trp | 13 | 151754 | 8.57e-5 | **0.833** | 0.000 | 0.417 | **0.766** | 0.370 | 22 | 12 AA, 1 LT |
| 197101048 | rs372416792 | p.Phe2735Val | 71 | 151798 | 4.68e-4 | **0.875** | 0.020 | 0.462 | **0.838** | 0.303 | 17 | 69 SA, 2 AA |
| 197101970 | rs752506858 | p.Phe2427Leu | 25 | 151764 | 1.65e-4 | 0.948 | 0.020 | 0.537 | **0.645** | 0.498 | 26 | 17 FN, 8 EU |
| 197102137 | rs190693455 | p.Arg2372Gly | 48 | 151816 | 3.16e-4 | **0.778** | 0.020 | 0.288 | **0.673** | 0.294 | 19 | 42 EA |
| 197102145 | rs372076208 | p.Gln2369Pro | 14 | 151786 | 9.22e-5 | **0.830** | 0.000 | 0.426 | 0.330 | 0.354 | 18 | 14 AA |

| | | | | | | | | | | | | |
|---|---|---|---|---|---|---|---|---|---|---|---|---|
| 197102158 | rs371212366 | p.Gln2365Glu | 15 | 151670 | 9.89e-5 | 0.990 | 0.000 | 0.732 | 0.905 | 0.489 | 24 | 15 AA |
| 197102400 | rs371321801 | p.Ser2284Phe | 18 | 151778 | 1.19e-4 | 0.945 | 0.000 | 0.475 | 0.887 | 0.260 | 22 | 17 AA, 1 EU |
| 197102455 | rs200800956 | p.Ala2266Thr | 13 | 151698 | 8.57e-5 | 0.999 | 0.000 | 0.718 | 0.944 | 0.462 | 23 | 8 EU, 2 AJ, 1 SA, 1 EA |
| 197103304 | rs141715950 | p.Met1983Leu | 175 | 151904 | 1.15e-3 | 0.933 | 0.000 | 0.234 | 0.956 | 0.235 | 23 | 131 EU, 15 AA, 15 LT |
| 197103354 | rs140834556 | p.Gln1966Pro | 19 | 151932 | 1.25e-4 | 0.712 | 0.010 | 0.367 | 0.772 | 0.299 | 14 | 13 EU, 6 AA |
| 197103467 | rs554548758 | p.Gln1928His | 12 | 151888 | 7.90e-5 | 0.999 | 0.000 | 0.714 | 0.935 | 0.581 | 20 | 9 SA, 1 EU, 1 AA |
| 197103667 | rs527602809 | p.Lys1862Glu | 37 | 151950 | 2.44e-4 | 0.861 | 0.020 | 0.490 | 0.893 | 0.292 | 22 | 18 LT, 16 SA |
| 197103741 | rs144969324 | p.Asn1839Ser | 29 | 151844 | 6.58e-6 | 0.973 | 0.030 | 0.677 | 0.972 | 0.592 | 24 | 25 EA, 1 SA, 3 LT |
| 197103799 | rs41299625 | p.Arg1818Cys | 156 | 151900 | 1.03e-3 | 0.998 | 0.000 | 0.617 | 0.988 | 0.538 | 25 | 122 EU, 14 AA, 2 SA, 1 EA |
| 197104113 | rs141297873 | p.Arg1713His | 12 | 151814 | 7.90e-5 | 0.965 | 0.010 | 0.589 | 0.907 | 0.586 | 19 | 6 LT, 3 EU, 2 ME, 1 AA |
| 197104114 | rs143911853 | p.Arg1713Cys | 16 | 151852 | 1.05e-4 | 0.995 | 0.020 | 0.619 | 0.854 | 0.626 | 24 | 10 AA, 5 EU, 1 EA |
| 197104266 | rs147839989 | p.Ile1662Ser | 16 | 151894 | 1.05e-4 | 0.999 | 0.000 | 0.770 | 0.973 | 0.838 | 26 | 13 EU, 2 LT |
| 197104276 | rs560847421 | p.Val1659Phe | 14 | 151830 | 9.22e-5 | 0.959 | 0.000 | 0.523 | 0.909 | 0.390 | 17 | 14 EA |
| 197104709 | rs375778017 | p.Ile1514Met | 12 | 151928 | 7.90e-5 | 0.944 | 0.010 | 0.434 | 0.808 | 0.321 | 17 | 12 AA |
| 197104755 | rs140119882 | p.Arg1499Leu | 68 | 151794 | 4.48e-4 | 0.999 | 0.000 | 0.858 | 0.957 | 0.740 | 26 | 66 AA,2 LT |
| 197104849 | rs189918817 | p.Ala1468Thr | 25 | 151870 | 1.65e-4 | 1.000 | 0.010 | 0.684 | 0.906 | 0.483 | 24 | 18 EU, 4 AA, 2 LT, 1 FN |
| 197105035 | rs145662604 | p.Ala1406Thr | 36 | 151804 | 2.37e-4 | 1.000 | 0.040 | 0.217 | 0.926 | 0.0284 | 24 | 36 AA |
| 197105077 | rs762353508 | p.Arg1392Ter | 10 | 151786 | 6.59e-5 | Stop gained | 1392/3477 | | | | | 7 LAT, 2 FN, EA |
| 197105103 | rs375240502 | p.Met1383Thr | 16 | 151948 | 1.05e-4 | 0.970 | 0.20 | 0.248 | 0.499 | 0.305 | 24 | 14 AA, 2 LT |
| 197105116 | rs768625023 | p.Ser1379Pro | 14 | 151950 | 9.21e-5 | 0.980 | 0.000 | 0.425 | 0.938 | 0.457 | 25 | 14 AA |
| 197117899 | rs376541100 | p.Ala1319Ser | 11 | 152060 | 7.23e-5 | 0.703 | 0.000 | 0.649 | 0.935 | 0.282 | 20 | 5 EA, 2 EU, 1 LT, 1 SA, 1AJ |
| 197122013 | rs142214506 | p.Ala1258Thr | 42 | 151896 | 2.77e-4 | 0.494 | 0.020 | 0.469 | 0.657 | 0.346 | 24 | 42 AA |
| 197133493 | rs375901171 | p.Arg759Gln | 16 | 152094 | 1.05e-4 | 0.981 | 0.000 | 0.388 | 0.467 | 0.375 | 25 | 12 EA, 4 EU |
| 197133494 | rs777867809 | p.Arg759Trp | 90 | 152112 | 5.92e-4 | 0.997 | 0.000 | 0.365 | 0.467 | 0.366 | 27 | 88 LT, 2 AA |
| 197139844 | rs112997359 | p.Ser650Cys | 26 | 152160 | 1.71e-4 | 0.538 | 0.010 | 0.150 | 0.518 | 0.019 | 16 | 26 AA |
| 197142535 | rs144049904 | p.Arg573Trp | 317 | 152070 | 2.08e-3 | 0.613 | 0.000 | 0.302 | 0.560 | 0.115 | 22 | 176 EU, 107LT, 15AA, 7AJ |
| ***CFHR1*** Chr 1. No predicted damaging variants | | | | | | | | | | | | |
| ***CFHR4*** Chr 1 | | | | | | | | | | | | |
| 196902480 | rs199547603 | p.Arg41Ter | 122 | 151486 | 8.05e-4 | Stop gained | 41/578 | | | | | 118 AA, 3 LT, 1 SA |
| 196902563 | rs201480125 | p.Trp68Ter | 65 | 151082 | 4.30e-4 | Stop gained | 68/578 | | | | | 54 AA, 5 EU, 3 LT |
| 196907017 | rs199731883 | p.Ser199Tyr | 31 | 151196 | 2.05e-4 | 0.989 | 0.000 | 0.501 | 0.915 | 0.250 | 17 | 31 AA |
| 196907017 | rs199731883 | p.Ser199Phe | 16 | 151196 | 1.06e-4 | 0.989 | 0.000 | 0.501 | 0.905 | 0.250 | 17 | 8 EU, 7 AJ, 1 LT |
| 196907468 | rs183705632 | p.Gly257Arg | 373 | 150964 | 2.47e-3 | 0.632 | 0.000 | 0.426 | 0.946 | 0.292 | 19 | 347 EU, 22 FN, 3 AJ |
| 196912826 | rs77776928 | p.Tyr362His | 241 | 151618 | 1.59e-3 | 0.970 | 0.010 | 0.354 | 0.971 | 0.234 | 21 | 223 AA, 11 LT, 2 EU, 1 ME |
| 196914648 | rs376896638 | p.Trp445Ter | 45 | 150272 | 2.99e-4 | Stop gained | 445/578 | | | | | 41 SA, 3 AA |
| 196915062 | rs575550227 | p.Gln488His | 19 | 151572 | 1.25e-4 | 0.749 | 0.010 | 0.249 | 0.522 | 0.190 | 14 | 17 AA, 2 LT |
| 196915069 | rs373767199 | p.Tyr491Asp | 12 | 151644 | 7.91e-5 | 0.968 | 0.000 | 0.684 | 0.989 | 0.482 | 21 | 11 AA, 1 LT |
| 196918321 | rs201727614 | p.Lys551Thr | 54 | 151532 | 3.56e-4 | 0.600 | 0.020 | 0.462 | 0.645 | 0.361 | 14 | 37 EU, 13 AA, 4 FN |
| ***JAK3*** Chr 19 | | | | | | | | | | | | |
| 17826901 | rs200580168 | p.Leu1073Phe | 50 | 152210 | 3.28e-4 | 0.946 | 0.000 | 0.372 | 0.251 | 0.478 | 25 | 29 EU, 18 FN |
| 17832539 | rs148688786 | p.Arg887His | 15 | 152218 | 9.85e-5 | 0.956 | 0.060 | 0.611 | 0.103 | 0.511 | 25 | 5 EU, 3 SA, 3 AJ, 2 EA |
| 17832681 | rs200077579 | p.Arg840Cys | 36 | 152208 | 2.37e-4 | 0.863 | 0.000 | 0.681 | 0.477 | 0.581 | 25 | 12 EU, 11 AM, 8 LT, 2 SA |
| 17834630 | rs149982493 | p.Pro764Leu | 15 | 150934 | 9.94e-5 | 0.619 | 0.000 | 0.535 | 0.908 | 0.387 | 25 | 14 AA, 1 EU |
| 17841389 | rs373046546 | p.Thr381Asn | 17 | 152188 | 1.12e-4 | 0.586 | 0.010 | 0.366 | 0.775 | 0.311 | 25 | 17 AA |
| 17842528 | rs202167678 | p.Val217Met | 16 | 152258 | 1.05e-4 | 0.905 | 0.020 | 0.456 | 0.694 | 0.362 | 22 | 14 AJ, 2 EU |
| 17843115 | rs149579831 | p.Gly160Cys | 15 | 152204 | 9.86e-5 | 0.508 | 0.000 | 0.403 | 0.574 | 0.360 | 26 | 13 EU, 1 LT |
| 17844395 | rs145500023 | p.Thr8Met | 83 | 152162 | 5.45e-4 | 0.973 | 0.060 | 0.518 | 0.179 | 0.293 | 22 | 63 EU, 10 AA, 8 LT, 1 FN |
| ***YJEFN3*** Chr19 | | | | | | | | | | | | |
| 19532678 | rs61749571) | p.Gly88Glu | 122 | 152208 | 1.84e-4 | 0.926 | 0.010 | 0.148 | 0.590 | 0.293 | 28 | 114 AA, 4 LT, 3 EU, 1 SA |
| 19535408 | rs141151340 | p.Cys167Trp | 14 | 152190 | 9.20e-5 | 0.906 | 0.010 | 0.251 | 0.831 | 0.426 | 15 | 9 EA, 3 SA, 1 EU, 1 LT |
| 19535632 | rs146733860 | p.Thr216Arg | 804 | 152200 | 5.28e-3 | 0.991 | 0.000 | 0.146 | 0.480 | 0.289 | 23 | 619 EU, 72 AA, 53FN, 38LT |
| 19535658 | rs200227282 | p.Val225Met | 231 | 152192 | 1.52e-3 | 0.960 | 0.000 | 0.281 | 0.882 | 0.263 | 26 | 222 AA, 6 LT, 1 EA |
| 19537411 | rs545964787 | p.Phe263Leu | 12 | 152212 | 7.88e-5 | 0.971 | 0.010 | 0.200 | 0.612 | 0.211 | 24 | 9 SA, 1 EU, 1 AA |
| 19537487 | rs1162772387 | p.Pro288Leu | 24 | 152246 | 1.58e-4 | 0.998 | 0.000 | 0.267 | 0.803 | 0.415 | 28 | 24 EU |
| ***CILP2*** Chr 19 | | | | | | | | | | | | |
| 19540306 | rs201714553 | p.Val89Gly | 68 | 152154 | 4.47e-4 | 0.742 | 0.000 | 0.154 | 0.927 | 0.000 | 27 | 62 FN, 6 EU |

| | | | | | | | | | | | | |
|---|---|---|---|---|---|---|---|---|---|---|---|---|
| 19540374 | rs200679414 | p.Arg112Cys | 316 | 152208 | 2.08e-3 | 0.958 | 0.040 | 0.092 | **0.709** | 0.000 | 26 | 185FN, 108EU, 12AA, 4LT |
| 19542944 | rs146789008 | p.Gly317Arg | 16 | 152192 | 1.05e-4 | 1.000 | 0.000 | 0.512 | 0.980 | 0.000 | 12 | 16 EU |
| 19544001 | rs146322371 | p.Glu486Lys | 42 | 152228 | 2.76e-4 | 0.984 | 0.000 | 0.245 | **0.828** | 0.000 | 24 | 37 AA, 3 EU, 2 LT |
| 19544086 | rs139539877 | p.Pro514Leu | 40 | 152266 | 2.63e-4 | **0.804** | 0.000 | 0.287 | **0.772** | 0.000 | 20 | 31 EU, 4 AA, 3 LT, 1 ME |
| 19544610 | rs758468698 | p.Gly689Ser | 10 | 152182 | 6.57e-5 | 1.000 | 0.010 | 0.321 | **0.809** | 0.000 | 27 | 10 EU |
| 19544925 | rs532951417 | p.Pro794Ser | 14 | 152214 | 9.20e-5 | 0.991 | 0.000 | 0.078 | 0.508 | 0.000 | 23 | 14 SA |
| ***D13RIK*** | Not annotated in human | | | | | | | | | | | |
| ***BABAM1*** | Chr 19 | | | | | | | | | | | |
| 17268634 | rs774465344 | p.Leu2Phe | 73 | 152012 | 4.80e-4 | In non-can. transcript; uncertain tests | | | | | | 56 EU, 8 LT, 7 AA |
| 17268640 | rs182898955 | p.Arg4Ser | 239 | 152040 | 1.57e-3 | In non-can. transcript; uncertain tests | | | | | | 235 AA, 3 LT, 1 EU |
| 17268756 | rs138467832 | p.Gly13Arg | 11 | 152114 | 7.23e-5 | In non-can. transcript; 0.098 PP | | | | | | 11 EA |
| 17271654 | rs115366940 | p.Gly115Ser | 499 | 152204 | 3.28e-3 | **0.812** | 0.010 | 0.237 | **0.657** | 0.000 | 18 | 457 AA, 21 LT, 13 EU, 1 SE |
| 17276515 | rs374131612 | p.Pro197Leu | 16 | 152154 | 1.05e-4 | 1.000 | 0.000 | 0.346 | **0.802** | 0.000 | 26 | 7 EU, 6 AA, 2 LT |
| 17276536 | rs550805861 | p.Thr204Met | 50 | 152194 | 3.29e-4 | 0.967 | 0.000 | 0.116 | **0.707** | 0.000 | 22 | 47 SA, 2, EU, 1 EA |
| 17276581 | rs200016717 | p.Arg219His | 52 | 152196 | 3.42e-4 | 0.999 | 0.000 | 0.315 | **0.818** | 0.000 | 32 | 26 EU, 19 AZ, 4 AA, 2 LT |
| ***ZFP709*** | Not annotated in human | | | | | | | | | | | |
| ***OLFR374*** | Not annotated in human | | | | | | | | | | | |
| ***ZFP372*** | Not annotated in human | | | | | | | | | | | |
| ***CAV2*** | Chr 7 | | | | | | | | | | | |
| 116499905 | rs374156174 | p.Pro42Ser | 11 | 152078 | 7.23e-5 | **0.900** | 0.000 | 0.907 | **0.875** | 0.828 | 27 | 11 EU |
| 116500365 | rs149320804 | p.Lys86Glu | 19 | 152192 | 1.25e-4 | 0.965 | 0.000 | 0.897 | **0.848** | 0.819 | 28 | 9 AJ, 5 LT, 4 EU |
| ***IQUB*** | Chr 7 | | | | | | | | | | | |
| 123452867 | rs146017906 | p.Tyr751Cys | 11 | 152068 | 7.23e-5 | ----- | 0.000 | 0.219 | **0.873** | 0.500 | 28 | 6 AA, 2 EA, 3 SA, 1 EU |
| 123457529 | rs145461420 | p.Ala682Val | 374 | 151988 | 2.46e-3 | ----- | 0.050 | 0.094 | **0.837** | 0.110 | 24 | 286 EU, 58 AA, 19 LT, 4 FN |
| 123461451 | rs60409072 | p.Arg638Gln | 480 | 151774 | 3.16e-3 | ----- | 0.010 | 0.208 | **0.887** | 0.317 | 26 | 457 AA, 14 LT, 2 EU, I ME |
| 123461494 | rs373146060 | p.Arg624Trp | 17 | 151688 | 1.12e-4 | ----- | 0.010 | 0.142 | **0.657** | 0.153 | 22 | 12 SA, 2 EU, 1 AA, 1 EA |
| 123464949 | rs1431266872 | p.Met548Val | 24 | 151880 | 1.58e-4 | ----- | 0.010 | 0.142 | **0.697** | 0.120 | 22 | 24 LT |
| 123464949 | rs199522362 | p.Leu547Phe | 88 | 151726 | 5.80e-4 | ----- | 0.020 | 0.511 | **0.869** | 0.528 | 26 | 31 AJ, 25 LT, 20 EU, 6 AA |
| 123469222 | rs74488511 | p.Thr525Ala | 16 | 152114 | 1.05e-4 | ----- | 0.010 | 0.234 | **0.649** | 0.328 | 25 | 16 EA |
| 123469294 | rs149316783 | p.Ile501Phe | 36 | 152194 | 2.37e-4 | ----- | 0.000 | 0.327 | **0.847** | 0.473 | 24 | 34 AA, 1 LT |
| 123479859 | rs144590492 | p.Gly449Val | 11 | 151998 | 7.24e-5 | ----- | 0.000 | 0.296 | **0.808** | 0.332 | 24 | 10 AA, 1 LT |
| 123479868 | rs148490443 | p.Ala446Val | 52 | 152062 | 3.42e-4 | ----- | 0.000 | 0.318 | **0.818** | 0.248 | 26 | 39 EU, 5 LT, 4 FN, 2 AA |
| 123479896 | rs143618132 | p.Leu437Phe | 30 | 152044 | 1.97e-4 | ----- | 0.000 | 0.448 | **0.864** | 0.498 | 24 | 26 EU, 2 LT, 1 AA |
| 123479960 | rs201714151 | p.Gln415His | 10 | 152136 | 6.57e-5 | ----- | 0.010 | 0.138 | **0.750** | 0.132 | 23 | 10 LT |
| 123496858 | rs370004563 | p.Val358Ile | 19 | 152006 | 1.25e-4 | ----- | 0.010 | 0.256 | **0.760** | 0.011 | 22 | 19 AA |
| 123502618 | rs201786274 | p.Tyr334Ter | 15 | 152130 | 9.86e-5 | Stop gained 334/791 | | | | | | 13 AJ, 2 EU |
| 123502948 | rs760765844 | p.Thr288Met | 15 | 151946 | 9.87e-5 | ----- | 0.000 | 0.467 | **0.921** | 0.481 | 25 | 4 EU, 4 AA, 2 LT, 2FN, 2EA |
| 123503030 | rs372053513 | p.Thr261Ala | 13 | 152202 | 8.54e-5 | ----- | 0.000 | 0.377 | **0.849** | 0.439 | 25 | 6 AJ, 4 LT, 1 AA. 1 SA |
| 123512135 | rs140652989 | p.Ser69Ter | 15 | 152164 | 9.86e-5 | Stop gained 69/791 | | | | | | 11 EU, 2 AM, 1 AA, 1 FN |
| ***PLD1*** | Chr 3 | | | | | | | | | | | |
| 171603170 | rs140343065 | p.Val1045Met | 19 | 152140 | 1.25e-4 | 0.999 | 0.000 | 0.186 | 0.955 | 0.371 | 27 | 18 AA, 1 LT |
| 171612393 | rs567323716 | p.Arg923His | 14 | 152182 | 9.20e-5 | 0.994 | 0.000 | 0.398 | **0.557** | 0.435 | 27 | 11 SA, 3 EA |
| 171620433 | rs191205035 | p.Tyr894Ser | 65 | 152124 | 4.27e-4 | 0.981 | 0.000 | 0.275 | 0.957 | 0.557 | 31 | 42 LT, 18 EU, 3 AA |
| 171644913 | rs143219477 | p.Tyr847Cys | 30 | 152206 | 1.97e-4 | 0.988 | 0.000 | 0.263 | 0.964 | 0.449 | 27 | 29 AA, 1 LT |
| 171644953 | rs142188788 | p.Gly834Ser | 12 | 152156 | 7.89e-5 | 0.999 | 0.000 | 0.177 | 0.443 | 0.138 | 25 | 8 AA, 4 EU |
| 171659247 | rs540439098 | p.Asp799Asn | 10 | 152216 | 6.57e-5 | 0.969 | 0.000 | 0.210 | **0.908** | 0.308 | 25 | 8 AA, 1 EU, 1 LT |
| 171659251 | rs140049612 | p.Ile797Met | 19 | 152284 | 1.25e-4 | 0.938 | 0.000 | 0.388 | 0.965 | 0.372 | 16 | 17 EU, 2 AA |
| 171662164 | rs745485055 | p.Arg746Cys | 11 | 152120 | 7.23e-5 | 0.956 | 0.000 | 0.269 | 0.934 | 0.625 | 32 | 7 LT, 3 EU, 1 AA |
| 171676758 | rs146885315 | p.Arg691Leu | 52 | 152224 | 3.42e-4 | 0.972 | 0.000 | 0.225 | **0.652** | 0.584 | 27 | 45 EU, 7 FN |
| 171676764 | rs139363214 | p.Ala689Val | 10 | 152198 | 6.57e-5 | 0.972 | 0.000 | 0.166 | **0.570** | 0.276 | 27 | 8 EU, 1 SA, 1 LT |
| 171676807 | rs202167212 | p.Arg675Trp | 11 | 152202 | 7.23e-5 | 0.999 | 0.000 | 0.181 | **0.729** | 0.420 | 32 | 8 LT, 2 EU, 1 FN |
| 171688711 | rs80085541 | p.Arg502Trp | 26 | 152106 | 1.71e-4 | 0.930 | 0.000 | 0.189 | **0.638** | 0.236 | 29 | 17 EU, 5 EA, 2 AA |
| 171688812 | rs374719323 | p.Val468Asp | 13 | 152188 | 8.54e-5 | 0.938 | 0.000 | 0.462 | 0.978 | 0.632 | 27 | 6 EU, 6 LT |
| 171688863 | rs141086000 | p.Pro451Leu | 10 | 152082 | 6.58e-5 | 0.996 | 0.000 | 0.294 | 0.958 | 0.597 | 26 | 9 EU, 1 AA |
| 171699746 | rs201602698 | p.Ala409Gly | 12 | 152214 | 7.88e-5 | 0.999 | 0.000 | 0.507 | **0.892** | 0.592 | 33 | 9 EU, 1 AA, 1 FN |
| 171709672 | rs371154750 | p.Arg317Trp | 11 | 152112 | 7.23e-5 | **0.666** | 0.000 | 0.590 | **0.768** | 0.740 | 29 | 6 EU, 2 AA, 2 SA, 1 ME |

| | | | | | | | | | | | | |
|---|---|---|---|---|---|---|---|---|---|---|---|---|
| 171713981 | rs370939132 | p.Val275Ile | 14 | 152110 | 9.20e-5 | **0.900** | 0.000 | 0.468 | **0.735** | 0.230 | 25 | 9 EA, 4 EU |
| 171724745 | rs149535568 | p.Gly237Cys | 139 | 152200 | 9.13e-4 | 0.985 | 0.000 | 0.932 | **0.609** | 0.809 | 24 | 116 EU, 23 AA |
| 171724759 | rs147193827 | p.Gly232Glu | 13 | 152174 | 8.54e-5 | **0.872** | 0.000 | 0.929 | **0.837** | 0.870 | 25 | 13 EA |
| ***FAT4*** | Chr 4 | | | | | | | | | | | |
| 125316596 | rs201009019 | p.Gly62Ala | 103 | 152170 | 6.77e-4 | 0.991 | 0.010 | 0.753 | 0.446 | 0.711 | 28 | 79 EU, 10 AA, 9 SA, 2 LT |
| 125318238 | rs370300509 | p.Asp609Glu | 40 | 152182 | 2.63e-4 | 0.167 | 0.000 | 0.404 | 0.972 | 0.475 | 15 | 37 AA, 2 LT, 1 EU |
| 125319589 | rs200529379 | p.Arg1060Cys | 10 | 152120 | 6.57e-5 | 0.990 | 0.000 | 0.402 | **0.630** | 0.474 | 26 | 6 FN, 4 EU |
| 125319916 | rs190175933 | p.Arg1169Trp | 16 | 152096 | 1.05e-4 | **0.594** | 0.000 | 0.404 | 0.446 | 0.467 | 25 | 7 EU, 7 AA, 1 FN, 1 LT |
| 125320411 | rs368607709 | p.Val1334Met | 19 | 152100 | 1.25e-4 | 0.995 | 0.000 | 0.449 | **0.825** | 0.349 | 26 | 10 AA, 7 EU, 1 EA |
| 125320580 | rs201934636 | p.Ile1390Thr | 24 | 152224 | 1.58e-4 | 0.127 | 0.000 | 0.188 | 0.946 | 0.384 | 25 | 20 EU, 4 AA |
| 125320694 | rs772537061 | p.Ser1428Phe | 16 | 152206 | 1.05e-4 | 0.969 | 0.020 | 0.024 | 0.382 | 0.360 | 25 | 16 LT |
| 125406973 | rs201887525 | p.Arg1801Trp | 20 | 152004 | 1.32e-4 | 0.998 | 0.000 | 0.019 | 0.391 | 0.289 | 23 | 15 EU, 3 EA, 1 SA, 1 AA |
| 125408666 | rs139716832 | p.Tyr1931Cys | 90 | 152066 | 5.92e-4 | 0.994 | 0.010 | 0.024 | **0.603** | 0.518 | 27 | 74 EU, 10 AA, 4 Lt, 1 FN |
| 125408675 | rs200026034 | p.Ile1934Thr | 13 | 152118 | 8.55e-5 | **0.892** | 0.000 | 0.330 | **0.763** | 0.338 | 25 | 13 AA |
| 125416574 | rs559079176 | p.Arg2324Trp | 22 | 152062 | 1.45e-4 | **0.594** | 0.000 | 0.029 | **0.923** | 0.409 | 22 | 19 LT, 1 EU, 1 SA |
| 125434305 | rs534302520 | p.Asn2360Ser | 26 | 152186 | 1.71e-4 | 0.992 | 0.000 | 0.526 | **0.841** | 0.441 | 25 | 16 AA, 7 LT, 2EU |
| 125446357 | rs139924242 | p.Ser2422Cys | 16 | 152106 | 1.05e-4 | 0.980 | 0.000 | 0.316 | **0.538** | 0.390 | 27 | 16 AA |
| 125446405 | rs376545643 | p.Val2438Phe | 12 | 152016 | 7.89e-5 | 0.926 | 0.010 | 0.232 | 0.154 | 0.296 | 24 | 9 EU, 2 AA |
| 125449646 | rs781160819 | p.Tyr2879Cys | 10 | 152160 | 6.57e-5 | 0.998 | 0.000 | 0.586 | 0.982 | 0.719 | 27 | 10 AA |
| 125449741 | rs148655455 | p.Tyr2911His | 25 | 152176 | 1.64e-4 | 0.104 | 0.000 | 0.583 | 0.976 | 0.727 | 26 | 25 EU |
| 125450461 | rs200702071 | p.Ala3151Thr | 41 | 152102 | 2.70e-4 | **0.594** | 0.020 | 0.014 | 0.127 | 0.198 | 24 | 23 SA, 8 EU, 5 LT, 2AA, 1AJ |
| 125450974 | rs181641041 | p.Val3322Met | 12 | 152208 | 7.88e-5 | 0.999 | 0.000 | 0.459 | 0.889 | 0.502 | 27 | 12 AA |
| 125451346 | rs756625118 | p.Ile3446Val | 10 | 152140 | 6.57e-5 | **0.511** | 0.040 | 0.325 | **0.772** | 0.290 | 24 | 10 EU |
| 125451386 | rs190233104 | p.Pro3459Leu | 11 | 152056 | 7.23e-5 | 0.226 | 0.000 | 0.204 | **0.718** | 0.342 | 23 | 5 SA, 5 EU 1 EA |
| 125451653 | rs79726583 | p.Tyr3548Cys | 23 | 152156 | 1.51e-4 | 0.924 | 0.010 | 0.498 | 0.940 | 0.335 | 27 | 23 AA |
| 125452163 | rs139635339 | p.Arg3718His | 19 | 152144 | 1.25e-4 | 0.987 | 0.040 | 0.303 | 0.508 | 0.303 | 25 | 13 EU, 4 AA, 1 Lt 1 SA |
| 125452390 | rs201859188 | p.Arg3794Trp | 28 | 152184 | 1.84e-4 | 0.990 | 0.000 | 0.432 | 0.486 | 0.626 | 24 | 14 LT, 8 EU, 4 AA, 1 FN |
| 125452472 | rs142857910 | p.Arg3821Gln | 11 | 152194 | 7.23e-5 | 0.942 | 0.010 | 0.370 | 0.091 | 0.387 | 23 | 9 AA, 1 LT, 1 SA |
| 125468638 | rs149250709 | p.Ala4011Val | 30 | 152060 | 1.97e-4 | **0.800** | 0.000 | 0.641 | **0.806** | 0.790 | 25 | 19 AA, 6 LT, 4 EU |
| 125468676 | rs138019311 | p.Arg4024Trp | 488 | 152088 | 3.21e-3 | 0.932 | 0.020 | 0.400 | 0.467 | 0.489 | 23 | 353EU,42AJ 33LT,10 other |
| 125468691 | rs140544054 | p.Ala4029Thr | 13 | 152094 | 8.55e-5 | **0.563** | 0.040 | 0.642 | **0.570** | 0.658 | 28 | 11 AJ, 2 EU |
| 125477197 | rs150804471 | p.Ile4114Met | 70 | 149916 | 4.67e-4 | **0.675** | 0.040 | 0.336 | 0.237 | 0.290 | 23 | 63 AA, 3 LT, 4 other |
| 125481563 | rs148170326 | p.Arg4216His | 61 | 152148 | 4.01e-4 | **0.725** | 0.030 | 0.344 | 0.490 | 0.404 | 23 | 54 EU, 3 AA, 2 LT, 1 FN |
| 125490017 | rs773916430 | p.Arg4401Cys | 10 | 151952 | 6.58e-5 | **0.809** | 0.000 | 0.579 | **0.821** | 0.582 | 25 | 8 AA, 2 EU |
| 125491107 | rs552252173 | p.His4764Arg | 12 | 152154 | 7.89e-5 | 0.978 | 0.000 | 0.208 | 0.179 | 0.141 | 15 | 11 AA, 1 LT |
| 125491128 | rs201633380 | p.Arg4771His | 31 | 152136 | 2.04e-4 | 0.994 | 0.000 | 0.546 | 0.179 | 0.286 | 27 | 28 LT, 2 AA, 1 EU |
| 125491178 | rs138173652 | p.Gly4788Arg | 14 | 152084 | 9.21e-5 | **0.896** | 0.000 | 0.811 | 0.202 | 0.619 | 24 | 6 LT, 6 EU, 1 AA, AJ |
| 125491582 | rs752311257 | p.Lys4922Asn | 10 | 152182 | 6.57e-5 | 0.058 | 0.000 | 0.333 | **0.615** | 0.106 | 11 | 8 FN, 2 EU |
| ***USP13*** | Chr 3 | | | | | | | | | | | |
| 179707000 | rs141550831 | p.Trp182Arg | 41 | 152118 | 2.70e-4 | ----- | 0.000 | 0.256 | **0.852** | 0.683 | 29 | 38 EU, 3 AA |
| 179708886 | rs200478981 | p.Ala245Val | 16 | 152082 | 1.05e-4 | ----- | 0.000 | 0.552 | 0.996 | 0.591 | 25 | 15 EU, 1 LT |
| 179740305 | rs776225594 | p.Pro438Leu | 12 | 152094 | 7.89e-5 | ----- | 0.010 | 0.537 | 0.508 | 0.534 | 25 | 11 EU, 1 AA |
| 179742254 | rs200761055 | p.Arg480Cys | 132 | 152190 | 8.67e-4 | ----- | 0.010 | 0.283 | 0.933 | 0.414 | 27 | 51 SA, 34 LT, 39 EU, 2 AA |
| 179754824 | rs200079140 | p.Arg672His | 12 | 152140 | 7.89e-5 | ----- | 0.000 | 0.207 | **0.909** | 0.368 | 32 | 5 EU, 4 AA, 2 EA, 1 SA |
| 179765710 | rs199954201 | p.Asn754Ser | 20 | 152262 | 1.12e-4 | ----- | 0.000 | 0.153 | **0.856** | 0.385 | 27 | 17 EU, 3 AA |
| ***GM1527*** | Not annotated in human | | | | | | | | | | | |
| ***ARHGAP28*** | Chr 18 | | | | | | | | | | | |
| 6824836 | rs188975691 | p.Arg66His | 28 | 152122 | 1.84e-4 | 0.989 | 0.040 | 0.221 | **0.603** | 0.000 | 27 | 18 EA, 5 AA, 1SA, 1EU, 1LT |
| 6882201 | rs186052265 | p.His452Arg | 14 | 152228 | 9.20e-5 | 0.362 | 0.010 | 0.085 | **0.790** | 0.165 | 24 | 11 Lt, 2 EU, 1 AA |
| 6890539 | rs74414891 | p.Arg615Gln | 74 | 152194 | 4.86e-4 | 0.980 | 0.020 | 0.092 | **0.826** | 0.179 | 25 | 71 AA, 3 LT |
| 6890539 | rs74414891 | p.Arg615Leu | 358 | 152190 | 2.35e-3 | 0.980 | 0.020 | 0.092 | **0.826** | 0.179 | 25 | 119SA, 94EU, 87EA, 18AA |
| 6898484 | rs756134118 | p.Trp641Ter | 15 | 151968 | 9.87e-5 | Stop gained 641/670 | | | | | | 15 AA |
| 6912083 | rs142653243 | p.Ala707Ser | 20 | 152118 | 1.31e-4 | **0.552** | 0.020 | 0.032 | **0.625** | 0.000 | 25 | 18 AA, 2 other |
| ***RMDN2*** | Chr 2 | | | | | | | | | | | |
| 37929288 | rs1049449941 | p.Ser4Tyr | 21 | 152148 | 1.38e-4 | **0.839** | 0.000 | 0.232 | **0.799** | 0.111 | 23 | 21 AA |
| 37929701 | rs199499480 | p.Ser142Gly | 62 | 152238 | 4.07e-4 | 0.959 | 0.000 | 0.307 | **0.881** | 0.199 | 25 | 52 EU, 9 AA, 1 LT |

| | | | | | | | | | | | | |
|---|---|---|---|---|---|---|---|---|---|---|---|---|
| 37951261 | rs140191801 | p.Arg16Ter | 391 | 151994 | 2.57e-3 | Stop gained 16/573 | | | | | | 360 AA, 4 EU, 3 ME |
| 37951541 | rs370260752 | p.Asp109Gly | 16 | 152096 | 1.05e-4 | 0.952 | 0.010 | 0.088 | ----- | 0.041 | 17 | 15 EU, 1 other |
| 37951915 | rs78569430 | p.Asp234Tyr | 1078 | 151994 | 7.09e-3 | 0.711 | 0.000 | 0.079 | ----- | 0.031 | 17 | 1005 AA, 40 LT, 12 SA |
| 37974065 | rs140735845 | p.Glu160Ter | 43 | 152150 | 2.83e-4 | Stop gained 338/573 | | | | | | 42 AA, 1 EU |
| 37975239 | rs143149132 | p.Arg219Ter | 10 | 152062 | 6.58e-5 | Stop gained 397/573 | | | | | | 8 EU, 1 AA, 1 LT |
| 37997430 | rs779349930 | p.Ser320Arg | 10 | 152110 | 6.57e-5 | 0.816 | 0.000 | 0.240 | 0.873 | 0.353 | 24 | 10 SA |
| 38004146 | rs147688126 | p.Asp370Val | 21 | 152168 | 1.38e-4 | 0.302 | 0.000 | 0.263 | 0.840 | 0.244 | 31 | 11 LT, 6 EU, 3 SA |
| ***MYOM1*** | Chr 18 | | | | | | | | | | | |
| 3067528 | rs764839969 | p.Gly1598Arg | 19 | 152212 | 1.25e-4 | ----- | 0.000 | 0.808 | 0.938 | 0.889 | 25 | 17 EU, 2 SA |
| 3067534 | rs375965840 | p.Val1596Met | 13 | 152176 | 9.20e-5 | ----- | 0.000 | 0.666 | 0.816 | 0.558 | 25 | 11 SA 3 AA |
| 3071887 | rs373419422 | p.Arg1571Cys | 26 | 152142 | 1.71e-4 | ----- | 0.010 | 0.324 | 0.879 | 0.568 | 33 | 20 AA, 5 LT, 1 EU |
| 3083991 | rs730880167 | p.Ile1459Lys | 61 | 152044 | 4.01e-4 | ----- | 0.000 | 0.268 | 0.760 | 0.377 | 32 | 59 EU, 2 FN |
| 3086046 | rs190368385 | p.Ile1415Val | 29 | 152220 | 1.91e-4 | ----- | 0.000 | 0.636 | 0.503 | 0.552 | 23 | 28 AA, 1 LT |
| 3086108 | rs374142554 | p.Asp1394Gly | 10 | 152218 | 6.57e-5 | ----- | 0.000 | 0.532 | 0.696 | 0.664 | 32 | 9 LT, 1 AA |
| 3086112 | rs371239754 | p.Lys1393Glu | 50 | 152230 | 3.28e-4 | ----- | 0.000 | 0.629 | 0.668 | 0.625 | 28 | 35 AA, 10 LT, 2 EU |
| 3102501 | rs147050513 | p.Arg1183Gln | 36 | 152094 | 2.37e-4 | ----- | 0.000 | 0.711 | 0.850 | 0.517 | 24 | 19 AJ, 7 EU, 6 LT, 3 AA |
| 3116411 | rs371066861 | p.Val1075Met | 17 | 152172 | 1.12e-4 | ----- | 0.000 | 0.439 | 0.807 | 0.519 | 24 | 7 EU, 6 AA, 3 LT |
| 3119890 | rs377512048 | p.Glu1033Lys | 24 | 152194 | 1.58e-4 | Stop gained 1033/1685 | | | | | | 23 AA, 1 LT |
| 3119949 | rs557671408 | p.Ala1013Val | 115 | 152152 | 7.56e-4 | ----- | 0.000 | 0.617 | 0.973 | 0.808 | 29 | 111 SA, 1AA, 1 LT |
| 3135618 | rs18388166 | p.Arg713His | 11 | 152078 | 7.23e-5 | ----- | 0.000 | 0.426 | 0.891 | 0.525 | 27 | 7 AA, 3 EU, 1 ME |
| 3135619 | rs375858580 | p.Arg713Cys | 14 | 151996 | 9.21e-5 | ----- | 0.000 | 0.463 | 0.532 | 0.535 | 32 | 11 EU, 1 AA, 1 EA |
| 3135624 | rs548747720 | p.Arg711His | 12 | 152096 | 7.89e-5 | ----- | 0.000 | 0.507 | 0.898 | 0.751 | 27 | 8 EU, 3 AA, 1 EA |
| 3141941 | rs200179081 | p.Lys675Glu | 44 | 152166 | 2.89e-4 | ----- | 0.010 | 0.390 | 0.896 | 0.499 | 32 | 37 EU, 5 AA, 1 LT, 1 FN |
| 3151801 | rs185573271 | p.Ser579Phe | 45 | 152174 | 2.96e-4 | ----- | 0.000 | 0.357 | 0.518 | 0.601 | 31 | 25 EU, 18 LT, 1 AA |
| 3154963 | rs370063864 | p.Gly543Arg | 22 | 152110 | 1.45e-4 | ----- | 0.000 | 0.389 | 0.846 | 0.599 | 31 | 20 LT, 2 AA |
| 3154984 | rs367903122 | p.Asp536Asn | 13 | 152202 | 8.54e-5 | ----- | 0.000 | 0.472 | 0.785 | 0.620 | 26 | 8 AJ, 4 AA, 1 EU |
| 3164292 | rs1278590083 | p.Tyr496Cys | 17 | 152234 | 1.12e-4 | ----- | 0.000 | 0.624 | 0.851 | 0.697 | 29 | 17 LT |
| 3164421 | rs368424215 | p.Ser453Ter | 17 | 152248 | 6.57e-6 | Stop gained 453/1685 | | | | | | 17 AA |
| 3174156 | rs1435418077 | p.Gly359Arg | 14 | 152218 | 9.20e-5 | ----- | 0.000 | 0.821 | 0.972 | 0.882 | 26 | 14 LT |
| ***ANKRD12*** | Chr 18 | | | | | | | | | | | |
| 9254838 | rs199503455 | p.Ser524Cys | 12 | 151964 | 7.90e-5 | 0.996 | 0.010 | 0.864 | 0.557 | 0.687 | 24 | 9 EU, 2 AA |
| 9255011 | rs187421957 | p.Arg582Gly | 13 | 151912 | 8.56e-5 | 0.962 | 0.000 | 0.866 | 0.727 | 0.593 | 24 | 13 EU |
| 9256635 | rs549560546 | p.Lys1123Thr | 11 | 152158 | 7.23e-5 | 0.994 | 0.000 | 0.588 | 0.742 | 0.543 | 26 | 8 AA, 2 LT, 1 EU |
| 9258816 | rs190468609 | p.Arg1850His | 14 | 152078 | 9.21e-5 | 0.996 | 0.010 | 0.466 | 0.224 | 0.598 | 27 | 5 EU, 5 LT, 3 other, 1 AA |
| 9258843 | rs369087314 | p.Gln1859Arg | 10 | 152206 | 6.57e-5 | 0.969 | 0.000 | 0.366 | 0.224 | 0.517 | 26 | 10 AA |
| ***XDH*** | Chr 2 | | | | | | | | | | | |
| 31339527 | rs142329784 | p.Arg1246Cys | 66 | 152202 | 4.34e-4 | 0.963 | 0.000 | 0.428 | 0.913 | 0.312 | 29 | 59 EU, 7 AA |
| 31339596 | rs116290580 | p.Arg1223Cys | 15 | 152202 | 9.86e-5 | 0.943 | 0.000 | 0.260 | 0.762 | 0.388 | 28 | 13 AA 2 EU |
| 31339598 | rs148235835 | p.Thr1222Ile | 41 | 152208 | 2.69e-4 | 0.995 | 0.000 | 0.470 | 0.945 | 0.769 | 27 | 34 EU, 6 LT, 1 AA |
| 31339668 | rs770434783 | p.Ala1199Val | 11 | 152228 | 7.23e-5 | 0.929 | 0.000 | 0.481 | 0.941 | 0.480 | 26 | 7 FN, 4 EU |
| 31341378 | rs139515054 | p.Ile1179Thr | 329 | 152166 | 2.16e-3 | 0.976 | 0.000 | 0.338 | 0.844 | 0.707 | 28 | 221 EU, 56 LT, 27AA, 19FN |
| 31348282 | rs142402298 | p.Thr1045Ala | 16 | 152180 | 1.05e-4 | 0.820 | 0.000 | 0.399 | 0.946 | 0.766 | 27 | 16 EU |
| 31349744 | rs148921536) | p.Glu971Lys | 13 | 152206 | 8.54e-5 | 0.649 | 0.000 | 0.167 | 0.372 | 0.259 | 26 | 11 EU, 1 AA, 1 LT |
| 31350118 | rs776794071 | p.Arg913Trp | 15 | 152184 | 9.86e-5 | 1.000 | 0.000 | 0.716 | 0.979 | 0.782 | 26 | 8 LT, 6 EU, 1 AA |
| 31365991 | rs148585342 | p.Ala814Val | 67 | 152156 | 4.40e-4 | 0.957 | 0.010 | 0.295 | 0.912 | 0.420 | 27 | 47 EU, 16 LT, 3 AA |
| 31366019 | rs140007233 | p.Arg805Trp | 11 | 152156 | 7.23e-5 | 0.998 | 0.000 | 0.362 | 0.942 | 0.442 | 27 | 10 EU, 1 AA |
| 31366070 | rs141050887 | p.Ile788Phe | 122 | 152064 | 8.02e-4 | 0.795 | 0.000 | 0.319 | 0.846 | 0.341 | 25 | 108 EU, 10 AA, 3 LT, 1 SA |
| 31366072 | rs149717617 | p.Arg787Gln | 45 | 152190 | 2.96e-4 | 0.972 | 0.000 | 0.209 | 0.433 | 0.274 | 28 | 25EU, 12LT, 3SA, 2FN, 1AA |
| 31366073 | rs148904866 | p.Arg787Trp | 13 | 152108 | 8.55e-5 | 0.999 | 0.000 | 0.380 | 0.913 | 0.506 | 25 | 5 EA, 4 EU, 4 AA |
| 31366946 | rs755854585 | p.Cys749Tyr | 12 | 152250 | 7.88e-5 | 0.918 | 0.000 | 0.259 | 0.513 | 0.297 | 25 | 6 EU, 6 LT |
| 31368050 | rs142848703 | p.Ile703Lys | 16 | 152208 | 1.05e-4 | 0.935 | 0.000 | 0.259 | 0.695 | 0.523 | 27 | 15 AA, 1 LT |
| 31368056 | rs527413763 | p.Asp701Val | 18 | 152148 | 1.18e-4 | 0.898 | 0.020 | 0.328 | 0.806 | 0.450 | 33 | 17 AA, 1 LT |
| 31372247 | rs201668777 | p.Arg613Trp | 11 | 152244 | 7.23e-5 | 0.803 | 0.000 | 0.071 | 0.873 | 0.159 | 23 | 5 EU, 4 LT, 2 AA |
| 31372358 | rs139772558 | p.Arg576Trp | 234 | 152204 | 1.54e-3 | 0.767 | 0.000 | 0.072 | 0.872 | 0.277 | 26 | 220 AA, 10LT, 1EU 3 other |
| 31375459 | rs148603901 | p.Arg508Gln | 28 | 152176 | 1.84e-4 | 1.000 | 0.000 | 0.379 | 0.928 | 0.692 | 26 | 21 AA, 6 EU, 1 ME |
| 31379899 | rs372033714 | p.Leu404Met | 26 | 152224 | 1.71e-4 | 0.779 | 0.020 | 0.158 | 0.926 | 0.278 | 24 | 24 AA, 2 LT |

| | | | | | | | | | | | | |
|---|---|---|---|---|---|---|---|---|---|---|---|---|
| 31383002 | rs142388231 | p.Ala346Val | 40 | 152142 | 2.63e-4 | 0.938 | 0.000 | 0.358 | 0.946 | 0.620 | 33 | 25 EU, 8 AJ, 4 AA, 2 LT |
| 31383080 | rs200384255 | p.Pro320Arg | 19 | 152198 | 1.25e-4 | 0.935 | 0.000 | 0.246 | 0.881 | 0.361 | 24 | 17 EU, 2 AA |
| 31383782 | rs138674014 | p.Leu287Val | 82 | 152160 | 5.39e-4 | 0.754 | 0.000 | 0.313 | 0.922 | 0.344 | 23 | 65 EU, 6 SA, 3 LT, 2 AA |
| 31383785 | rs375277410 | p.Glu286Gln | 29 | 152196 | 1.91e-4 | 0.985 | 0.010 | 0.267 | 0.931 | 0.421 | 24 | 28 AA, 1 LT |
| 31386510 | rs149487595 | p.Arg233Cys | 11 | 152106 | 7.23e-5 | 0.983 | 0.000 | 0.205 | 0.934 | 0.432 | 25 | 8 EU, 3AA |
| 31387845 | rs201332711 | p.Pro206Leu | 28 | 152150 | 1.84e-4 | 1.000 | 0.000 | 0.247 | 0.956 | 0.351 | 28 | 18 EU, 8 AJ, 1 AA, 1 LT |
| 31397681 | rs200810102 | p.Arg161Gln | 13 | 152158 | 8.54e-5 | 0.710 | 0.010 | 0.293 | 0.129 | 0.431 | 27 | 11 LT, 2 EU |
| 31397700 | rs145413551 | p.Pro155Thr | 56 | 152218 | 3.68e-4 | 1.000 | 0.030 | 0.309 | 0.492 | 0.478 | 23 | 49 AA, 5 LT, 2 EU |
| 31398605 | rs377748577 | p.Pro134Leu | 18 | 152206 | 1.18e-4 | 1.000 | 0.010 | 0.351 | 0.709 | 0.576 | 25 | 17 FN, 1 EU |
| 31398642 | rs768170843 | p.Met122Leu | 10 | 152160 | 6.57e-5 | 0.997 | 0.000 | 0.303 | 0.601 | 0.476 | 25 | 9 EU 1 AA |
| 31398678 | rs147286954 | p.Gly110Ser | 15 | 152160 | 9.86e-5 | 0.965 | 0.000 | 0.596 | 0.738 | 0.503 | 26 | 15 LT |
| 31414634 | rs773456900 | p.Asn11Lys | 10 | 152154 | 6.57e-5 | 0.881 | 0.000 | 0.372 | 0.985 | 0.357 | 22 | 8 EU, 1 AA, 1 LT |
| ***MTCL1*** | Chr 18 | | | | | | | | | | | |
| 8705986 | rs1000002519 | p.Asp109Gly | 22 | 148310 | 1.48e-4 | 0.911 | 0.030 | ----- | 0.461 | ----- | 28 | 16 EU, 3 AJ, 1 LT |
| 8718626 | rs139379863 | p.Arg59Gln | 24 | 152198 | 1.64e-4 | 0.943 | 0.000 | 0.656 | 0.775 | 0.455 | 31 | 14 EU, 10 LT, 1 AA |
| 8720458 | rs150009615 | p.Lys107Gln | 16 | 152226 | 1.05e-4 | 0.596 | 0.000 | 0.585 | 0.742 | 0.373 | 25 | 14 AJ, 1 EU, 1 other |
| 8783737 | rs554525790 | p.Leu209Val | 15 | 152140 | 9.86e-5 | 0.808 | 0.000 | 0.077 | 0.333 | 0.231 | 19 | 14 AA, 1 LT |
| 8783938 | rs145413531 | p.Arg276Cys | 18 | 152172 | 1.18e-4 | 0.999 | 1.000 | 0.158 | 0.871 | 0.332 | 25 | 15 AA, 2 LT, 1 SA |
| 8784010 | rs200271750 | p.Arg300Trp | 95 | 152178 | 6.24e-4 | 0.978 | 0.000 | 0.108 | 0.717 | 0.150 | 28 | 60 FN, 31 AA, 4 EU |
| 8784055 | rs780158820 | p.Arg315Trp | 14 | 152166 | 1.05e-4 | 0.761 | 0.000 | 0.042 | 0.065 | 0.112 | 24 | 14 AA, 2 EU |
| 8784259 | rs367857172 | p.Arg383Trp | 15 | 152158 | 9.86e-5 | 0.978 | 0.000 | 0.182 | 0.932 | 0.207 | 23 | 10 AA, 3 EU, 1 LT, 1 SA |
| 8784316 | rs149027252 | p.Arg402Cys | 15 | 152132 | 9.86e-5 | 0.961 | 0.000 | 0.225 | 0.797 | 0.224 | 23 | 10 AA, 3 EU, 1 LT, 1 other |
| 8784335 | rs372469977 | p.Arg408Gln | 12 | 152170 | 7.89e-5 | 0.983 | 0.000 | 0.207 | 0.862 | 0.344 | 25 | 10 AJ, 2 AA |
| 8784389 | rs140267953 | p.Ser426Leu | 88 | 152132 | 5.78e-4 | 0.941 | 0.000 | 0.229 | 0.799 | 0.339 | 23 | 59EU, 13LT, 9AA, 4SA, 1EA |
| 8784413 | rs201851779 | p.Ser434Leu | 13 | 152136 | 8.54e-5 | 0.996 | 0.010 | 0.239 | 0.799 | 0.278 | 25 | 7 EU, 3 EA, 2 AA, 1 LT |
| 8784512 | rs147236744 | p.Thr467Met | 43 | 152164 | 2.83e-4 | 0.990 | 0.010 | 0.146 | 0.625 | 0.111 | 22 | 36 AA, 4 EU, 3 LT |
| 8784520 | rs201898483 | p.Arg470Cys | 18 | 152160 | 1.18e-4 | 0.920 | 0.020 | 0.268 | 0.668 | 0.289 | 21 | 11 LT, 7 EU |
| 8784530 | rs143105042 | p.Thr473Met | 61 | 152162 | 4.01e-4 | 0.840 | 0.030 | 0.098 | 0.609 | 0.038 | 20 | 58 LT, 2 LT, 1 other |
| 8784656 | rs146651585 | p.Ile515Thr | 15 | 152140 | 9.86e-5 | 0.647 | 0.000 | 0.178 | 0.609 | 0.253 | 25 | 12 AA, 2LT, 1 other |
| 8784736 | rs114113780 | p.Pro542Ser | 17 | 152202 | 1.12e-4 | 0.708 | 0.020 | 0.139 | 0.668 | 0.142 | 23 | 15 EA, 2 SA |
| 8784793 | rs150423289 | p.Arg561Trp | 35 | 152214 | 2.30e-4 | 0.993 | 0.000 | 0.409 | 0.669 | 0.409 | 26 | 24 EU, 6 SA, 3 LT, 2 AA |
| 8784836 | rs375168888 | p.Asp575Gly | 16 | 152092 | 9.20e-5 | 0.928 | 0.010 | 0.373 | 0.625 | 0.348 | 27 | 12 EU, 1 AA, 1 LT |
| 8786041 | rs770066213 | p.Gly613Arg | 10 | 151910 | 6.58e-5 | 0.760 | 0.000 | 0.248 | 0.654 | 0.331 | 23 | 4 EU, 4 LT, 1 AA, 1 EA |
| 8796356 | rs150964075 | p.Arg712Gln | 35 | 152142 | 2.30e-4 | 0.938 | 0.000 | 0.338 | 0.766 | 0.307 | 27 | 30 EU, 4 AA, 1 other |
| 8807014 | rs532812464 | p.Ala853Val | 12 | 152184 | 7.89e-5 | 0.991 | 0.000 | 0.366 | 0.856 | 0.187 | 26 | 9 EU, 2 AA, 1 SA |
| 8813183 | rs139744939 | p.Arg937Trp | 11 | 152148 | 7.23e-5 | 0.999 | 0.000 | 0.428 | 0.773 | 0.487 | 29 | 9 AA, 1 EU, 1 SA |
| 8824734 | rs367681723 | p.Arg1075His | 10 | 152132 | 6.57e-5 | 0.994 | 0.000 | 0.357 | 0.773 | 0.292 | 28 | 5 EU, 3 AA, 1 LT, 1 other |
| 8824768 | rs149983879 | p.Lys1086Asn | 27 | 152152 | 1.77e-4 | 0.575 | 0.000 | 0.106 | 0.661 | 0.136 | 23 | 24 EU, 3 AA |
| 8824977 | rs138372811 | p.Tyr1156Cys | 12 | 152180 | 7.89e-5 | 0.949 | 0.020 | 0.128 | 0.678 | 0.231 | 25 | 12 AA |
| 8825022 | rs149277971 | p.Pro1171Leu | 14 | 152234 | 9.20e-5 | 0.705 | 0.000 | 0.147 | 0.709 | 0.352 | 26 | 12 EU, 2 AA |
| 8825108 | rs781258348 | p.Arg1200Cys | 14 | 152188 | 9.20e-5 | 0.968 | 0.030 | 0.126 | 0.700 | 0.314 | 28 | 13 AA, 1 EU |
| 8825531 | rs144633551 | p.Ser1341Gly | 49 | 152182 | 3.22e-4 | 0.577 | 0.000 | 0.209 | 0.664 | 0.265 | 26 | 49 AA |
| 8825582 | rs116331286 | p.Val1358Met | 10 | 152212 | 6.57e-5 | 0.869 | 0.010 | 0.143 | 0.496 | 0.100 | 24 | 8 EU, 1 AA, 1 other |
| 8825586 | rs116291960 | p.Ser1359Leu | 75 | 152198 | 4.93e-4 | 0.988 | 0.000 | 0.276 | 0.751 | 0.385 | 24 | 45 EA, 15 AA, 8 LT, 2 SA |
| 8825720 | rs371275636 | p.Asp1404Asn | 11 | 152156 | 7.23e-5 | 0.780 | 0.000 | 0.182 | 0.712 | 0.249 | 28 | 5 AA, 3 LT, 2 EU, 1 other |
| 8825787 | rs374048104 | p.Arg1426Leu | 14 | 152196 | 9.20e-5 | 1.000 | 0.000 | 0.481 | 0.760 | 0.553 | 31 | 13 AJ, 1 EU |
| 8825858 | rs114538532 | p.Ile1450Val | 721 | 152068 | 4.74e-3 | 0.656 | 0.010 | 0.066 | 0.363 | 0.159 | 21 | 423AA,110LT,89SA,60EA, 33EU |
| 8825861 | rs146614505 | p.Asp1451Asn | 31 | 152174 | 2.04e-4 | 0.994 | 0.000 | 0.404 | 0.712 | 0.368 | 28 | 29 EU, 1 LT, 1 AA |
| 8825903 | rs61753867 | p.Arg1465Trp | 32 | 152204 | 2.10e-4 | 0.982 | 0.010 | 0.189 | 0.615 | 0.205 | 24 | 23 AJ, 8 EU, 1 AA |
| 8825915 | rs372807917 | p.Arg1469Trp | 21 | 152238 | 1.38e-4 | 0.989 | 0.000 | 0.276 | 0.723 | 0.281 | 25 | 8 EA, 9 AA, 1 EU, 1LT, 1SA |
| 8826092 | rs139202553 | p.Glu1528Lys | 15 | 152254 | 9.85e-5 | 0.999 | 0.000 | 0.307 | 0.828 | 0.499 | 29 | 8 EU, 5 LT, 1 AA, 1 FN |
| 8826105 | rs574852095 | p.Arg1532Gln | 17 | 152224 | 1.12e-4 | 0.999 | 0.000 | 0.349 | 0.828 | 0.500 | 31 | 5 SA, 5 EU, 4 EA, 2AA, 1LT |
| 8831796 | rs201909742 | p.Arg963Ter | 10 | 152152 | 6.57e-5 | Stop gained 1998/2011 | | | | | | 8 AA, 2 EU |
| ***TSPYL2*** | Chr X | | | | | | | | | | | |
| 53082628 | rs782451065 | p.Gln44Glu | 23 | 111425 | 2.06e-4 | 0.596 | 0.010 | 0.094 | 0.731 | 0.091 | 22 | 20 EU, 2 FN, 1 AA |
| 53085058 | rs781934817 | p.Phe368Ile | 12 | 112446 | 1.07e-4 | 0.994 | 0.000 | 0.436 | 0.934 | 0.565 | 26 | 12 EU |

Many in-frame deletions of unknown effect

***FOXR2*** Chr X

| | | | | | | | | | | | | |
|---|---|---|---|---|---|---|---|---|---|---|---|---|
| 55624507 | rs374320995 | p.Arg266Cys | 14 | 112660 | 1.24e-4 | 0.097 | 0.020 | 0.727 | **0.773** | 0.375 | 10 | 10 EU, 1 AA, 1 LT, 1 EA |
| 55624623 | rs745397944 | p.Leu304Phe | 10 | 112848 | 8.86e-5 | **0.582** | 0.000 | 0.680 | 0.261 | 0.237 | 18 | 5 SA, 5 EA |

---

**Bioinformatic analysis of prioritized candidate genes' functions, expression in vascular tissues/cells, and evidence suggesting potential involvement in the collaterogenesis pathway.**

The following websites (underlined) and references therein were examined for the genes in Table 1 to query their functions, mutants, expression by vascular cells/tissues, etc, ascertained in mouse (M) and human (H). We relied in particular on information provided in the sections within the sites given in brackets: Ensemble-UniProtKB: Function/Induction, Subcellular location, Expression (in vascular tissues, endothelial and smooth muscle cells) [Bgee, Genevisible (GV)], Protein-protein interactions [STRING], Genetic variation; MGI (M): Mutations, Alleles, Phenotypes; Alliance-Gene page (M): Description, Phenotypes, Disease associations, Alleles and variants, Models (KO, Tg, etc); HGCN-Genecards (H): Summaries/descriptions, Function [GWAS Catalog, Phenotypes, Animal models], Variants, Disorders, Localization, Expression in vascular tissues and cells [GTEx, HPA], Pathways; NCBI Gene (M/H); Medline/Pubmed (M/H).

***Canq5:***

Sox13, SRY-Box Transcription Factor 13, a SOX family whose members regulate embryonic development and cell fate, is involved in T-cell development and brown adipocyte differentiation, and inhibits Wnt signaling. It has recently been linked to angiogenesis in gliomas and promotes angiogenesis on forced expression in glioma endothelial cells.[1] Expression in mouse is very high in embryonic arteries[2] and embryonic lung endothelial cells and intermediate in arteries and kidney vasculature of adults, compared to other tissue/cell types, while in adult human it is very high in arteries, endothelial and smooth muscle cells (Bgee, GV, GTEx, HPA). The above findings support a potential role for Sox13 in collaterogenesis.

Cyb5r1, cytochrome b5 reductase 1, is an NADH-cytochrome b5 reductase that is localized to the plasmalemma, mitochondria and endoplasmic reticulum. Cyb5r family members are involved in desaturation and elongation of fatty acids, cholesterol biosynthesis, drug metabolism, and EMT and the infiltrative tumor cell phenotype in colon cancer.[3] Expression in mouse and human is intermediate in brain vessels, aorta, arteries, veins, endothelial and smooth muscle cells (Bgee, GV, GTEx, HPA), compared to other tissue/cell types. Cyb5r1 is also responsible for reduction of methemoglobin, which in its oxidized form is defective in binding oxygen. Loss of function variants of Cyb5r1 cause familial methemoglobinemia and susceptibility to functional anemia and tissue hypoxia. Cyb5r1 is upregulated in diabetic retinopathy which is accompanied by hypoxia-induced pathological angiogenesis.[4] These findings suggest Cyb5r1 may play a role in the hypoxia→VEGFA→Flk1→ Rabep2 component of the collaterogenesis signaling pathway. However, Cyb5r2 was also shown to be upregulated in diabetic retinopathy, but unlike above,[4] inhibited angiogenesis in vitro,[5] and to act as a tumor suppressor in nasopharyngeal carcinoma by attenuating angiogenesis.[6]

Rgs1, regulator of G-protein signaling-1, is a GTPase-activating protein that limits $G_\propto$ and $G_{\beta}\gamma$ signaling from certain G protein-coupled receptors (GPCRs).[7] Deficiency in *Rgs1* has been linked to abnormal trafficking of immunoglobulin-secreting cells and autoimmune-like disease. However little information exists regarding its role in vascular development. Expression is intermediate in arteries, ECs, and smooth muscle cells of mouse and human.[8] (see also Bgee, Genevisible) Knockout/down of *Rgs1* suppressed angiogenesis in a rat model of arthritis[9] and reduced expression of VEGF, proliferation of ECs, and neovascularization in the ischemic mouse choroid.[10] These reports are consistent with our finding that wildtype (B6 allele) AAV9-*Rgs1* induced collateral formation in CC016 mice and conclusion that deficient Rgs1 activity contributes to reduced collaterogenesis and the sparse collaterals evident in CC016 mice. However, findings from other studies could suggest that increased signaling by GPCRs that contribute to collaterogenesis, resulting from interacting with a deficient Rgs1, would be expected to lead to *more* abundant collaterals in CC016, and therefore that wildtype AAV9-*Rgs1* should have had no effect (or decreased collateral number if the latter exhibits competitive binding): SDF1→CXCR4 signaling, which is limited by Rgs1,[7] promotes formation of a collateral pathway after arterial occlusion of the dorsal aorta in zebrafish[11] and additional collaterals in the newborn heart[12] and adult brain[13] of mice exposed to prolonged hypoxemia. Also, hypoxia reportedly increased RGS1 expression through a HIF-dependent mechanism, resulting in dampening of SDF1-induced migration of human mesenchymal stem cells[14]—cells which have been shown to be involved in collateral formation in adult heart.[15] In addition, sphingosine-1-phosphate (S1P) which interacts with Rgs1[9] has been implicated in formation of arteriole anastomoses induced by an S1P receptor agonist in the mouse dorsal skin fold chamber[16] and formation of additional collaterals following MCA occlusion.[17-19] Of note, pharmacological inhibition of angiotensin converting enzyme induced the formation of arterial anastomoses in chick embryo, leading to the possibility that on or both of the GPCR activating ligands, angiotensin and kallikrein, is involved in collaterogenesis;[20] however AT1 receptor signaling is strongly restrained by Rgs8 rather than by Rgs1.[21] The above findings support a potential role for Sox13 in collaterogenesis.

Aspm, abnormal spindle microtubule assembly, which is located in the cytoplasm and nucleus, is involved in mitotic spindle regulation with a preferential role in neurogenesis. Expression of Aspm in mouse and human is low in blood vessels (Bgee, GTEx, HPA) with the exception of being very high in choroid and iris microvessel endothelial cells (GV-H). KO mice evidence severe microcephaly; their vascular status has not been examined.

Cfhr1 and Cfhr4, complement factor H-related 1 and 4, respectively, are secreted plasma proteins synthesized primarily by hepatocytes. They are involved in complement regulation, and can associate with lipoproteins and may participate in lipid metabolism. Mutations, primarily linked to age-related macular degeneration, have not been associated with any vascular phenotype; KO models has not been reported. Cfhr1 is expressed at high levels in coronary artery, aorta, aorta and smooth muscle and endothelial cells (Bgee-H, GV-H, HPA), compared to other tissues/cell types. Cfhr4 is expressed at intermediate levels in aorta, coronary and tibial arteries, endothelial cells and human blood outgrowth endothelial cells (Bgee-H, GV-H, HPA). No publications for either protein have reported involvement in vessel formation or function.

***Canq6:***

Jak3, Janus kinase 3, is a non-receptor tyrosine kinase involved in cell growth, development, differentiation, T- and B-cell development, and signaling in innate and adaptive immunity, in particular by type I interleukin receptors and other cell surface receptors to downstream effectors including STAT proteins which activate gene expression. Mutations in humans are associated with severe combined immunodeficiency disease, and point mutations in mice develop autoimmune inflammatory bowel disease and altered susceptibility to malaria and bacterial infection. Jak 3 is expressed predominantly in immune cells. Expression is low in mouse vessels (Bgee, GV) and in human is intermediate in aorta and high in temporal and coronary arteries and endothelial and smooth muscle cells (Bgee, GTEx, HPA). Jak3 negatively regulated re-endothelization, endothelial proliferation, and proliferation and alpha-actin expression in intimal cells after wire injury of the carotid artery.[22] Consistent with Jak-STAT signaling in hypoxic Hif1∝-mediated induction of several angiogenic genes, Jak3 is required for VEGF-mediated proliferation, migration and angiogenesis in human microvascular endothelial cells.[23] It also regulates adherens junctions and epithelial mesenchymal transition through β-catenin.[24] The above findings support a potential role for Jak3 in collaterogenesis.

Yjefn3, YjeF N-terminal domain containing 3 (ApoA-I binding protein 2), binds ApoA-I on HDL and accelerates cholesterol efflux from endothelial cells to high-density lipoprotein (HDL). It also activates endothelial SREBF2, the master transcription factor for cholesterol biosynthesis, which in turn transactivates Notch and promotes hematopoietic stem and progenitor cell emergence from the hemogenic endothelium.[25] Increased Yjefn3 was shown to interfere with R2 dimerization and VEGFA induced angiogenesis in mouse endothelial cells and to inhibit angiogenesis in zebrafish, whereas knockdown of Yjefn3 in the latter resulted in dysregulated sprouting/branching angiogenesis. These results indicate biphasic effects on embryonic angiogenesis depending on Yjefn3 level.[25,26] Expression in adult mice and humans is low in arteries, veins, endothelial and smooth muscle cells (Bgee, GV. GTEx, HPA). The above data suggest that Yjefn3 may play a role in collaterogenesis.

Not entries are given herein for the last 6 genes in Table 1 for *Canq6* because each was deemed very unlikely to be involved in collaterogenesis based on the following: highly restricted expression and function in cell types unlikely to be involved in collaterogenesis (Bgee, Genevisible (GV); the Pubmed search terms angiogenesis, vascular development, endothelial cell function, and endocytic vesicle trafficking did not return references.

***Canq7:***

Cav2, caveolin-2, forms a stable complex with caveolin-1 to target lipid rafts and caveolae formation which regulate endocytic vesicle trafficking. It is involved in essential cellular functions, including signal transduction, lipid metabolism, cellular growth control and apoptosis. Caveolin-2 (and caveolin-1) also binds G-protein alpha subunits and can regulate their activity, promotes Src-mediated phosphorylation and binding of Ras1, Src and Nck1, colocalizes with focal adhesions where it links integrin subunits (including β3 on endothelial cells) to the tyrosine kinase FYN, is an initiating step in coupling integrins to the Ras-ERK pathway, and binds to endothelial cell NOS3 (eNOS) and promotes NO production. Caveolin-2 also induces nuclear translocation of Stat3,[27] and modulates endothelial cell proliferation: KO mice exhibit increased proliferation of endothelial cells in lung in association with loss of an inhibitory effect of caveolin-2 on TGF-β--Smad2/3 signaling (through internalization of TGFBR1 from membrane rafts and subsequent degradation).[28,29] On the other hand, reduced angiogenesis and growth of subcutaneously implanted tumors was observed in KO mice[30]—responses also seen in caveolin-1 KO mice.[31] Although not significant, infarct volume after permanent MCA occlusion was 30% lower in *Cav2* KO than wildtype mice, whereas it was significantly increased by 88% in *Cav1* KOs (n=10-16 per group).[32] Interestingly, a SNP in the 3'UTR of *CAV2*, rs4727833, has been linked to cardioembolic stroke (P-value $2x10^{-7}$, OR 1.09, CI 1.06-1.13, MAF 0.45);[33] This SNP is in linkage disequilibrium ($r^2$ 0.96, MAF 0.45) with an intergenic SNP for CAV2 that has been linked in three other GWAS studies to atrial fibrillation (P-value $7x10^{-13}$), heart rate response to exercise (P-value $2x10^{-9}$) and heart rate response to recovery post exercise (P-value $7x10^{-10}$) (GWAS Catalog); these SNPs have eQTLs for increased expression of *CAV2* by tibial artery ($2.7x10^{-10}$ and $1.2x10^{-9}$, respectively, both with normalized effect size 0.23) (GTEx); Differences in coronary collateral status, which evidence suggests may cause differences in tissue oxygen delivery during increases in oxygen demand,[34] could contribute to these traits. Expression of

*Cav2* is very high in mouse brain microvessels and aorta (Bgee, GV) and in human arteries, veins, endothelial and smooth muscle cells (Bgee, CV, HPA). The above findings support a potential role for caveolin-2 in collaterogenesis.

Iqub, IQ motif and ubiquitin domain containing, is found in the cytoplasm and nucleoplasm, especially in sperm cells. Two publications have investigated the function of Iqub: One tested siRNA against expression of putative cilia genes in cell lines with findings suggesting it may play a role in primary cilium assembly or maintenance and in hedgehog signaling.[35] A second study found that Iqub was upregulated in breast cancer and promoted proliferation and migration of a breast cancer cell line via activating Akt/GSK3β/Wnt/β-catenin signaling.[36] In mouse and human, expression is high in sperm, ciliated respiratory and endometrial cells, very low in other tissues including brain vessels and aorta, and not detected in endothelial and smooth muscle cells (Bgee, GTEx, HPA).

***Canq8:***

Pld1, phospholipase D1, which is selective for phosphatidylcholine, is stimulated by $PIP_2$ and $PIP_3$, and to a lesser degree by RhoA, Rac1 and CDC42. Pld1 is localized to the plasma membrane, endosomes, golgi, endoplasmic reticulum, and perinuclear area, and is a critical signal in numerous cellular pathways including receptor signal transduction, vesicle trafficking at the plasma and nuclear membranes, and regulation of mitosis. Endothelial cells deficient in Pld1 show reduced PI3K-Akt and MEK-ERK signaling in response to VEGFR2 activation, reduced integrin-dependent adhesion, migration, and proliferation, and reduced angiogenesis in vitro, in tumor xenographs, hypoxic retina, and in zebrafish during development,[37-39] Expression in mouse is intermediate in carotid and brain vessels (Bgee) and in human is intermediate in aorta, carotid and tibial arteries, endothelial and smooth muscle cells, low in saphenous vein (Bgee, HPA) and is very high in blood outgrowth endothelial cells (GV). The above findings suggest that Pld1 may play a role in collaterogenesis.

Fat4, FAT atypical cadherin 4, which localizes to the plasma membrane and primary cilia, is involved in cell junctions and adhesion, planar cell polarity, receptor and YAP1-hippo signaling, neuroprogenitor and mesenchymal cell proliferation, and development including that of the lymphatic vasculature through a lymphatic endothelial cell autonomous manner.[40] KO mice show neonatal lethality with reduced body size and altered morphology in several non-vascular tissue types. Recent evidence suggests a role for Fat4 in pericyte differentiation and proliferation.[41] Expression in mouse is very high in carotid, aorta and brain vessels (Bgee) and in human is very high in aorta, popliteal, tibial and coronary arteries, endothelial and blood outgrowth endothelial cells and vascular smooth muscle cells (Bgee, GV, GTEx, HPA). The above findings suggest Fat4 may be involved in collaterogenesis.

Usp13, ubiquitin specific peptidase 13 (isopeptidase T-3), is located in the cytosol and nucleus. Usp13 deubiquitinates BECN1, MITF, SKP2, USP10 and other proteins and is involved in autophagy and endoplasmic reticulum-associated degradation. Among its binding partners is the dual specificity phosphatase PTEN. In mouse lung fibroblasts, USP13 appears to regulate stability of Smad 4 leading to ECM expression and cell differentiation, and both translocate to the cytoplasm in response to TGF-β1.[42] Expression in mouse is intermediate in aorta and carotid artery (Bgee), while in human it is high in vena cava, saphenous vein, smooth muscle and endothelial cells, intermediate in temporal and popliteal arteries, and low in coronary artery and aorta (Bgee, HPA, GTEx).

GM1527, predicted gene 1527, is predicted to activate small GTPases. Expression in mouse is restricted to morula blastoderm and sperm cells (Bgee). No information exists for this gene in human, and Pubmed lists no functional information.

***Canq9:***

Arhgap28, Rho GTPase activating protein 28, which resides primarily in the cytoplasm, negatively regulates RhoA signaling which has a number of downstream effectors including stress fiber assembly. There are no reports examining Arhgap28 in vascular wall cells. In one KO mouse model, blood vessel development was impaired and early postnatal lethality occurred. In another, KOs were viable and appeared normal, however an increase in the closely related Arhgap6 led the authors to suggest that it may have compensated for loss of Arhgap28.[43] Arhgap28 interacts in protein binding assays with Arhgap29 (STRING). Expression of the latter was recently found to be induced during lumenization of endothelial cell cords early in vascular development, and through its inhibition of RhoA, to interfere with non-muscle myosin II mediated generation of endothelial cell tension, resulting in maintenance of lumen formation (Ondine Cleaver, pers comm). It is possible that Arhgap28 may assist Arhgap29 in the above and contribute to collaterogenesis. Consistent with the above findings, *Arhgap28* is expressed at high levels in mouse carotid artery, brain vessels, endothelial cells and smooth muscle cells, compared to other tissues (Bgee). In human, it is expressed at high levels in endothelial and smooth muscle cells, and at intermediate levels in tibial, coronary and hepatic arteries and saphenous vein, compared to low levels in other tissues (excepting sperm, lungs and kidney glomerulus where expression is high) (Bgee, GV, HPA).

Rmdn2, regulator of microtubule dynamics 2, binds with microtubules and presumably influences their functions, including chromosomal segregation in *C elegans.*[44] However, little is known about Rmdn2 function aside of its association with amyotrophic lateral sclerosis and congenital glaucoma. Located in the cytoplasm, its expression in mice is low in aorta, carotid artery and brain vessels (Bgee), while in human levels are moderate in aorta, coronary artery, endothelial and smooth muscle cells (Bgee, GTEx, HPA), compared to other tissues and cell types.

Myom1, myomesin (skelemin), binds myosin, meromyosin, and titin in striated muscle where it is thought to link intermediate filaments to the M-disk. It also binds probable hydrolase PNKD which can activate the NF-kappa-B signaling pathway, binds β1/β3 integrins which are important in angiogenesis during development, and is present in endothelial cells.[45,46]. Myomesin is also found in the nucleus where it affects gene expression.[45] In mouse, *myomesin* is expressed at intermediate levels in aorta, carotid artery, and brain blood vessels (Bgee) and in human at high levels in coronary and tibial arteries, intermediate levels in aorta and vena cava, and at very high levels in striated muscle, smooth muscle, and endothelial cells (GTEx, Bgee, HPA) compared to other cell types. The above findings support a possible role for myomesin in collaterogenesis.

No entries are given herein for Ankrd12, Xdh and Mtcl1 for *Canq9* because each was deemed very unlikely to be involved in collaterogenesis based on the following: highly restricted expression and function in cell types unlikely to be involved in collaterogenesis (Bgee, Genevisible (GV); the Pubmed search terms angiogenesis, vascular development, endothelial cell function, and endocytic vesicle trafficking did not return references.

***Canq10:***

Tspyl2, Testis-specific Y-encoded-like protein 2, is subject to x-inactivation, resides in the nucleolus and at lower levels in the cytoplasm, participates in nucleosome assembly, modulates transcription including glutamate receptor expression in embryonic brain, inhibits cell-cycle progression and is involved in embryonic stem cell differentiation. It is ligated by TGFβ1, and is involved in both the canonical (Smad-based) and noncanonical TGFβ signaling pathways,[47] the latter activating MAPKs including ERK, p38, and JNK, and that it signals to PI3K-AKT and Rho-like GTPases.[48] *Tspyl2* is induced in a hypoxia-inducible factor-1α (Hif1∝)-dependent manner[49] and augments TGFβ signaling in vascular wall cells.[50] The above pathways are important in vasculogenesis and angiogenesis during development. Experimentally induced diabetes upregulated vascular Tspyl2, it is upregulated in human diabetic aorta, and deletion in diabetic mice promotes aorta aneurysm formation and reduced diabetes-induced accumulation of kidney collagen III and IV and tubular injury. Its pro-fibrotic activity is mediated through cross-talk with TGFβ and negatively regulates NF-kB signaling.[48,50-53] The above support a possible role for Tspyl2 in collaterogenesis. Reinforcing this, mouse *Tspyl2* is widely expressed among tissues from embryonic day E10.5 to P7, including in brain, and at intermediate levels in arteries and brain blood vessels in adult (Bgee, GV). Human *TSPYL2* expression is moderately high in arteries, smooth muscle, and endothelial cells (GTEx, Bgee, HPA). One KO mouse model was embryonic lethal, while another resulted in behavioral and brain morphological alterations that mirror a number of neurodevelopmental psychiatric traits.

Foxr2, forkhead box R2, a member of the Fox family of transcription factors, resides at low levels in the nucleoplasm, is expressed at low levels in all tissues and cells (Bgee, GV, GTEx, HPA), and is involved in embryogenesis, differentiation, epithelial-mesenchymal transformation and metabolism. Foxr2 induces expression of sonic hedgehog which is important in early vascular development, stimulates angiogenesis in association with increased VEGF-A expression, promotes tumor progression, and is aberrantly expressed in a variety of cancers at levels that correlate with cancer progression[54-56]. Downregulation of Foxr2 strongly reduced Shh, Gli1, and Ptch1 protein in a tumor cell line.[54] In glioma endothelial cells, Foxr2 ligated the promotors and increased expression of ZO-1, occludin, and claudin-5.[57] Foxr2 was also shown to activate FAK/SRC signaling in brain tumor cells.[58] All of the above signals are involved in vasculogenesis and angiogenesis during development, consistent with a potential contribution of Foxr2 to collaterogenesis.

**Genes in Supplemental table III that were subjected to in vivo AAV9 assay:**

Cxcr4, chemokine (C-X-C motif) receptor 4. See Rgs1, above, for rationale for this gene's rank and selection for testing.

C1galt1, core 1 synthase, glycoprotein-N-acetylgalactosamine 3-beta-galactosyltransferase 1, is plasma lemma protein that is a precursor for many extended O-glycans in glycoproteins, and has a central role in many processes, including angiogenesis and thrombopoiesis. Xia et al[55] found that expression in mice is primarily in endothelial, hematopoietic, and epithelial cells during development. They also reported that KO mice develop brain hemorrhage that was uniformly fatal by embryonic day 14, with chaotic microvascular networks in brain, distorted capillary lumens and defective association of endothelial cells with pericytes and extracellular matrix. The above findings support a possible role for myomesin in collaterogenesis.
